# Selfing-outcrossing as a gradient, not a dichotomy: propensity for selfing varies within a population of hermaphroditic animals

**DOI:** 10.1101/2021.10.27.466132

**Authors:** Anja Felmy, Alena B. Streiff, Jukka Jokela

**Author notes:** Corresponding author: Anja Felmy.

## Abstract

For mating-system evolution, individual-level variation is essential. In self-compatible hermaphrodites, individuals may vary in their lifetime propensity for selfing, which consists of individual and environmental components. According to the reproductive assurance hypothesis explaining partial selfing, a key environmental factor is mate availability, which fluctuates with population density.

We quantified individual variation in selfing propensity in a hermaphroditic snail by manipulating mate availability, recording mating behaviour, estimating selfing rates from progeny arrays, and measuring female lifetime fitness. We found substantial among-individual variation in selfing propensity, including pure outcrossers, pure selfers, and two types of plastic individuals. This variation only manifested in the laboratory; for the highly dense field population, data suggest full outcrossing. Meanwhile, experimental levels of mate availability (low *versus* moderate) neither significantly affected selfing propensities nor selfing rates.

Instead, selfing propensities had an individual, environment-independent component. Our results imply that selfing propensities are partially heritable and, when selected on, cause mean selfing rates to evolve. We propose that genetic variation in selfing propensities offers a reconciliation between the reproductive assurance hypothesis and its limited empirical support in animals: distributions of selfing propensities vary temporally and spatially, thus obscuring the relationship between population density and realised selfing rates.

## Background

The mating system structures populations and species with respect to sexual behaviour. In self-compatible hermaphrodites, it primarily describes the relative frequency of self-fertilisation (hereafter selfing) *versus* outcrossing. The mating system adopted by such hermaphrodites determines the quantity and genetic quality of offspring (1). Consequently, it has pervasive effects on individual fitness and on the evolutionary and demographic properties of populations and species (2, 3, 4).

Variation in selfing rates has often been studied at the species level (5, 6, 7, 8), where the distribution of selfing rates has been found to be bimodal, with more outcrossers and selfers than mixed maters (0.2 < selfing rate *s* < 0.8). In plants, many studies have documented variability between populations of a single species (e.g., 9, 10, 11, 12). For example, in 51% of 105 surveyed plant species, populations fell into several mating-system categories (13). Our knowledge on animal populations is sparser. Inter-population variation was observed in the androdioecious nematode *C. elegans*, whose hermaphrodites cannot mate with each other and where all outcrossing is due to males (14). In the few studies where selfing rates, or the delay in the onset of reproduction when selfing (termed “waiting time”, 15), have been estimated across populations of self-compatible hermaphrodites capable of outcrossing, inter-population variation was the norm (16, 17, 18, 19, 20, 21, 22, 23). It is therefore important to characterise species by more than one population estimate, lest crucial differences between populations may be overlooked.

Variation in selfing rates among individuals is equally essential. Evolutionary change primarily happens at the population level; among-individual variation constitutes the raw material natural selection acts upon. If individuals within isolated populations of simultaneous hermaphrodites were uniform in their ability to self and had equal opportunities to outcross, the selection gradient for the selfing rate would be zero: selection can only act on traits that have both phenotypic and genetic variance (24).

Despite the importance of among-individual variation in selfing rates, empirical studies are not usually designed to capture variation at this level. Reliable estimates of individual selfing rates require a large number of rather large progenies to be genotyped with genetic markers of sufficient resolution to unequivocally identify outcrossing events (25, 26). Existing studies have shown that selfing rates varied within populations of plants (e.g., 27, 28, 29) and animals (e.g., 18, 19, 20, 30, 31, 32, 33, 34). However, especially in studies of animals, many estimates rely on small numbers of weakly polymorphic loci, which hampers the detection of outcrossing events, and so may be biased (25, 26). Substantial differences among individuals were also found in proxies of the selfing rate, such as in animals’ waiting time (22, 35, 36) and in the presence of seeds in self-pollinated plants (37, 38).

Individual selfing rates likely depend on genetic factors, environmental conditions, and genotype-by-environment interactions. Genetic self-incompatibility systems are well-known in plants (39, 40) and have recently been discovered in ascidians (41) and oomycetes (42). Differences in waiting times among families, suggestive of genetic variation in selfing rates, have been found in snails (15, 22, 35) and flatworms (36). A key environmental determinant of individual selfing rates is the availability of mating partners. Selfing provides reproductive assurance when mates are scarce, but due to inbreeding depression and other costs of selfing (43, 44) may be unpreferred if outcrossing is possible. Yet, to date there are no direct tests of how individual selfing rates in animals respond to differential mate availability; existing studies (45, 46) lack selfing rates estimated using molecular markers. Here we aim to close this knowledge gap.

Individual variation in selfing can be assessed by estimating individuals’ lifetime propensity for selfing, here defined as a biological property of an individual (the ability and readiness to self) and its environment (mate availability), loosely following Brandon’s propensity interpretation of fitness (47). This definition allows a separate exploration of the individual (supposedly genetic) *versus* ecological determinants of variation in selfing rates, the need for which recent studies have abundantly shown (9, 13, 23, 48).

The freshwater snail *Radix balthica* is an ideal species to study within-population variation in selfing propensity. Laboratory studies have confirmed that self-compatibility is geographically widespread yet not omnipresent within populations (49, 50). There is also tentative evidence for between-and within-population variation in selfing rates based on molecular genetic markers (32, 33, 51). The population studied here is almost fully outcrossing in the field (26), in accordance with its high density.

We derived individuals’ lifetime propensity for selfing for two laboratory environments with differential mate availability. We first ascertained how readily 274 laboratory-reared, unmated snails selfed in isolation. We then assigned them to one of two mating treatments, simulating (i) a low population density and scarcity of mating opportunities, or (ii) conditions in the ancestral field population, where density was high and multiple paternity within egg clutches common (52). We recorded individual mating behaviour, the lifetime production of eggs in isolation (necessarily selfed) and thereafter (potentially outcrossed), the proportion of developing eggs, and the proportion of surviving juveniles. For 56 individuals across both treatments (32 of them once-and 24 repeatedly paired), we estimated individual selfing rates in eggs produced post-isolation from progeny arrays.

Assuming variation in post-isolation selfing rates, we had two *a priori* hypotheses: Selfing post-isolation is more common under low mate availability (hypothesis 1) and in snails that selfed in isolation (hypothesis 2). Hypothesis 1 posits that selfing provides reproductive assurance at low density but is unfavoured when outcrossing opportunities are abundant (43, 44). If supported, it suggests selfing propensities have a strong ecological component. Hypothesis 2 postulates that a period of isolation reveals classes of individuals with different propensity for selfing, which remain largely unaffected by environmental change. It thus assumes selfing propensities to be stable, fundamental properties based in genetic differences. The hypotheses are not mutually exclusive; ecological and genetic factors can jointly shape individual selfing rates. We further explored how variation in selfing propensities is maintained by investigating their association with individuals’ sexual drive, female lifetime reproductive success, and their progenies’ embryonic development and juvenile survival.

## Methods

### Study system

1. *R. balthica* is a simultaneously hermaphroditic snail inhabiting the shallow littoral zone of European lakes (51, 53). In Lake Zurich, Switzerland, it is an annual with non-overlapping generations. Snails hatch from eggs in spring, reach sexual maturity in winter, reproduce from March to May, and then die (52). During the single breeding season, individuals may copulate repeatedly in both sexual roles and lay hundreds of eggs in distinct egg clutches (52). Copulation is unilateral. Lymnaeid snails can store allosperm and should not be allospermlimited even if they do not mate continuously (54).

### Experimental snails

We caught 86 adult snails (P0 mothers) at peak breeding season (24 April 2013) in Uerikon, Lake Zurich, Switzerland. In the laboratory at Eawag-Duebendorf, Switzerland (room temperature 18°C), they were kept individually in 200 ml plastic cups filled with aged tap water and fed *ad libitum* with organic lettuce. Water was changed once a week. In these cups, P0 mothers laid egg clutches until they died of natural causes in late spring 2013 (details in reference 49). P0 mothers were not allowed to mate in the laboratory; their offspring (F1 snails) were outcrossed using allosperm stored from copulations in the field, as shown by genotyping P0 mothers and F1 snails (see Results). Previous work showed that egg clutches collected from the ancestral field population throughout the breeding season often had multiple fathers and contained many unique parental genotypes (52). This suggests that the fathers of F1 snails, while unknown, came from a large and diverse population, and that F1 offspring of the same P0 mother were often just half siblings. Hence, an overrepresentation of certain fathers among F1 snails is unlikely. Each egg clutch laid by P0 mothers was placed in a separate water-filled 40 ml plastic cup. After 17 days, when hatching was imminent, each clutch was transferred to a larger 200 ml cup. F1 snails were reared in sibling groups and individually from age 19.4 ± 4.1 weeks onwards (mean ± SD) to preserve their virginity (details in 49). This start of isolation is very early; paired control snails only began to mate at age 41.3-45.7 weeks. F1 snails were fed *Spirulina* powder mixed with finely ground chalk and flakes of fish food as juveniles, and organic lettuce from age 37.0-42.3 weeks onwards. In spring 2014, mating trials (see next section) were conducted with 274 F1 snails derived from 38 P0 mothers and 108 sibling groups. The effects of mating trials on mating behaviour, fecundity, and the population growth rate are described elsewhere (49). For a subset of 56 of these 274 F1 snails, derived from 22 P0 mothers and 38 sibling groups, we estimated selfing rates (see Genetic analysis).

### Mating trials

Mating trials began in early May 2014 when F1 snails were 52.1 ± 1.0 weeks old. By that time, all F1 snails had experienced a prolonged period of isolation and 38.7% had begun to reproduce through selfing. In the ancestral field population, breeding started in early March (52), and laboratory control snails had started to copulate several weeks earlier (see above).

A detailed description of the design of mating trials is provided in (49). In brief, two types of mating trials were conducted. In the first type, 137 snails had one mating opportunity with one partner, simulating low population density and a scarcity of mating opportunities. In the second type, 137 snails had six mating opportunities over the course of six weeks, each with a different partner, simulating moderate population density. Hence, no F1 snail remained permanently isolated; they were all paired at least once. Each mating opportunity lasted for 10.3 ± 0.8 hours, long enough for copulations in both sexual roles, and happened during daylight hours. Once-paired snails were thus allowed to mate for only ∼0.5% of the length of their natural, three-month breeding season. Additionally, having just a single mating partner creates substantial obstacles for outcrossing (*e.g.*, unsuccessful sperm transfer, cryptic female choice, sperm competition with autosperm, genetic incompatibility between sperm and egg). So, if a snail’s sole mating partner happened to be sterile or incompatible, then this was part of the low-density environment we wanted to simulate. Hence, once-paired snails were clearly mate-limited. By contrast, a choice of six mating partners may come close to conditions in the ancestral, fairly dense field population, where field-collected egg clutches had 2.1 fathers on average (range: 1–9; 52). Given that not all mate encounters will lead to copulations, and the above-mentioned obstacles to outcrossing, the average number of potential mating partners of a free-living snail throughout its life is probably a multiple of 2.1. The contrast between once-and six-times-paired snails thus enables a rigorous test of hypothesis 1 (“selfing post-isolation is more common under low mate availability”). F1 snails were euthanised one week after repeatedly paired snails had had their sixth mating opportunity, when mortality began to increase reflecting the snails’ natural, annual life cycle.

All except two mating partners were from among the 274 snails, so snails could act both as mothers and fathers. The first mating partner of each snail was size-matched, to ensure that anatomical or developmental differences did not constrain copulation. All snails were individually marked (details in 49). Mating pairs were set up randomly with respect to relatedness. Consequently, in 22/465 pairings (4.7%) involving 42 F1 snails, mating partners had the same P0 mother (*i.e.*, full or half siblings). Of these 42 snails, 37 were paired repeatedly, and so were also paired with four to five unrelated snails each, reducing potential effects of mating partner relatedness. The five snails solely paired with a sibling were excluded from all analyses, as was a snail with missing data on female reproductive output. Hence, our sample sizes are 131 snails paired once (32/56 snails with selfing rate estimates) and 137 snails paired repeatedly (24/56).

Mating trials were recorded on time-lapse movies (one frame/30 sec). From these we extracted the number of partners each snail had copulated with as a male and as a female. We also collected all egg clutches F1 snails laid both before and after being paired to measure female lifetime reproductive success. We placed clutches in individual 40 ml cups and counted the number of eggs and how many of these contained developed embryos 17 days after oviposition (details in 49). Then each F1 snail’s clutches were transferred to two 2 l plastic containers each, where selfed (clutches laid in isolation) and potentially outcrossed (clutches laid post-isolation) F2 family groups were reared separately. F2 snails were fed a mixture of *Spirulina* algae, finely ground chalk, and flakes of fish food.

### Genetic analysis

We selected those F1 snails (n = 56) for selfing rate estimation that post-isolation produced at least twelve F2 offspring that reached 12.3 ± 0.5 weeks of age. At that time, some F2 juveniles were used in a field experiment and the rest frozen for genetic analysis. As by then 53.4 ± 18.1% of F2 offspring born post-isolation had died, we here estimate “secondary” selfing rates after the expression of potential in-or outbreeding depression in early development. For logistic reasons, we did not genotype any juveniles from eggs laid shortly before the experiment was terminated. Hence, the genotyped offspring of repeatedly paired snails originate from eggs laid before mating opportunity 4 (n = 6 snails), 5 (n = 15 snails), and 6 (n = 3 snails), respectively. Offspring of these snails can thus be sired by three to five mating partners.

Of 40 F1 snails, 15 or more F2 juveniles were available for genotyping; of these we genotyped at least 15, resulting in 16.0 ± 1.9 genotyped juveniles per F1 snail (range: 15-22). Of the 16 F1 snails with fewer than 15 juveniles available for genotyping, all juveniles were genotyped. This resulted in 8.1 ± 3.6 genotyped juveniles per F1 snail (range: 3-14). We also genotyped all 86 P0 and 274 F1 snails to estimate population-level inbreeding coefficients using Genetix version 4.05.2 (55) and to ascertain whether F1 snails themselves were selfed (Table S1). Snails were genotyped for ten highly polymorphic microsatellite loci (GenBank Accession No. KX830983-KX830992) developed specifically for this population by Ecogenics GmbH (Zurich, Switzerland). The genotyping and scoring protocol is described elsewhere (26). One locus showed a non-negligible frequency of null alleles (locus Rb_3, KX830985) and was excluded from all analyses.

### Paternity analysis

Paternity analyses were performed using a custom-built R routine. We excluded 41 juveniles (5.3% of 768 juveniles genotyped) from the analysis because they did not meet the inclusion criteria (fewer than six loci of both mother and offspring genotyped successfully, OR genotypes lacking a maternal allele at more than two loci, OR genotypes with six successfully genotyped loci but two loci without a maternal allele). This left us with 13.0 ± 4.1 successfully genotyped juveniles per family (range: 3-18). For these 727 juveniles, we counted the loci consistent with being outcrossed, *i.e.*, those with one non-maternal allele. We then compared each juvenile’s genotype to those of all its candidate fathers, defined as all the potential mates of a juvenile’s mother, irrespective of whether copulations had been observed. A candidate father was considered a perfect match if he possessed a juvenile’s paternal allele at every locus successfully genotyped in the juvenile, the mother, and himself. Candidate fathers lacking a juvenile’s paternal allele at one or several loci were considered non-perfect matches and deemed increasingly unlikely fathers the higher the number of father-offspring mismatches. Additionally, we repeated all paternity analyses using COLONY version 2.0.5.9 (56, see Supporting Methods), known for its highly accurate parentage assignments (57).

Using these procedures, paternity was assigned to 717 (98.6%) juveniles with high-quality genotypes, originating from 55 F1 snails (13.0 ± 3.9 assigned offspring per F1 snail). Paternity assignments were inconclusive for all of the offspring of one F1 snail (see Table S2 for numbers of genotyped, successfully genotyped, and assigned juveniles per family). For 705 juveniles, the R routine and COLONY produced identical assignments. Of the remaining twelve juveniles, five were left unassigned by the R routine, four by COLONY, and three were assigned by both but to different fathers; here we gave preference to the R routine (details in Table S2). Allelic diversity was high across loci (Table S1), yielding powerful paternity analyses. The mean probability of being selfed for offspring identified as selfed was 0.79 ± 0.22, while that of offspring identified as outcrossed was 0.00 ± 0.00 (Table S2). Outcrossed offspring were assigned to a father with a probability of 0.93 ± 0.16 (Table S2).

### Statistical analysis

We tested whether F1 snails were more likely to produce at least some selfed juveniles postisolation (no/yes) under low mate availability (test of hypothesis 1) and upon successful selfing in isolation (test of hypothesis 2) using a GLMM with binomial errors (n = 55). Fixed effects were the experimental treatment (once-vs. repeatedly paired), the number of developed embryos produced in isolation, and the number of male and female partners snails copulated with. P0 mother identity was used as a random intercept to account for potential effects of relatedness.

We used eight GLMMs to test if snails with different propensities for selfing differed in their lifetime number of eggs (female LRS) and developed embryos, in their lifetime proportion of undeveloped embryos, and in the proportion of developed embryos that died before reaching the juvenile age of 12.3 ± 0.5 weeks, when surviving F2 juveniles were genotyped. We estimated these models for F1 snails with successfully estimated selfing rates (four models, n = 50; the five outcrossers were excluded due to low sample size) and for all female fertile snails, here defined as snails with developed embryos (four models, n = 123 or 113, depending on the trait analysed). We fitted GLMMs using negative binomial errors (nbinom 1 family) for the number of eggs and developed embryos, and using Gaussian errors for the proportion of undeveloped embryos and dead juveniles. Despite being proportions, the latter two response variables did not deviate significantly from normality (one-sample Kolmogorov-Smirnov tests: *D* ≥ 0.06, *p* ≥ 0.33). Furthermore, model assumptions were not violated, as shown by both conventional diagnostic plots and those produced by DHARMa, a simulation-based approach to assess the model fit (58). In all eight models, adult body size was used as a continuous covariate, the propensity for selfing as a factor with three levels (“selfer”, “plastic mixer”, and “plastic switcher” when n = 50, and “apparent selfer”, “apparent plastic snail”, and “apparent outcrosser” when n = 123 or 113), and P0 mother identity as a random intercept. In all models except those of the number of eggs, we also included the “pair identity” on mating opportunity 1 as a random intercept. This accounts for the non-independence of sexual functions within pairs of once-paired snails, where it was, *e.g.*, impossible for one snail to remain unmated while its sole mating partner mated. In models of the number of eggs, the likelihood of selfing postisolation (see above), and the likelihood of mating in both sexual roles among snails with estimated selfing rates (see below), we did not add the pair identity on mating opportunity 1 because doing so resulted in convergence problems. However, of 31 once-paired snails with estimated selfing rate, only six were paired with one another, while 25 were paired with repeatedly paired snails or with once-paired snails whose selfing rate was not estimated. Any non-independence in selfing rate estimates is thus likely small.

Furthermore, we tested if snails that mated in both sexual roles (no/yes) differed in their propensity for selfing using a GLMM with binomial errors and including snails with successfully estimated selfing rates only (n = 50, see above). The propensity for selfing (three levels: “selfer”, “plastic mixer”, “plastic switcher”), adult body size (continuous covariate) and P0 mother identity (random intercept) were used as predictors.

Finally, we used two-tailed Pearson’s Chi-square tests to investigate effects of the experimental treatment of F1 snails (once-vs. repeatedly paired) on multiple paternity (supporting analysis), selfing (test of hypothesis 1), and the propensity for selfing (test of hypothesis 1). Specifically, 2×2 contingency tables were used to test for treatment effects on the number of F2 progenies with multiple paternity, the number of F2 progenies with selfed offspring, and the overall number of selfed F2 offspring. Treatment effects on the propensity for selfing in all female fertile F1 snails (categories: “apparent selfer”, “apparent plastic snail”, and “apparent outcrosser”) and only in snails with selfing rate estimate (categories: “selfer”, “plastic mixer”, “plastic switcher”, and “outcrosser”) were analysed using 3×2 and 4×2 contingency tables, respectively.

Analyses were performed using R v. 4.2.2 (59). GLMMs were fitted using “glmmTMB” (60). The model fit was assessed using “DHARMa” (58). We tested the significance of random effects with log-likelihood ratio tests (full model vs. model without random effect in question). Values are given as mean ± SD. Scatter plots were prepared using “beeswarm” (61) and depict data points superimposed on boxplots. The entire code and model output is provided as an R Markdown document (62).

## Results

### Frequency of selfing and multiple paternity

Selfed offspring were found in 43.6% of genotyped families (24/55; Fig. 1a). Nine families showed low selfing rates of 20% at most and 15 families high ones of 83% or more. Families with intermediate selfing rates (0.20 < *s* < 0.80) were absent. Overall, 23.2% of offspring were identified as selfed (166/717).

**Figure 1.**
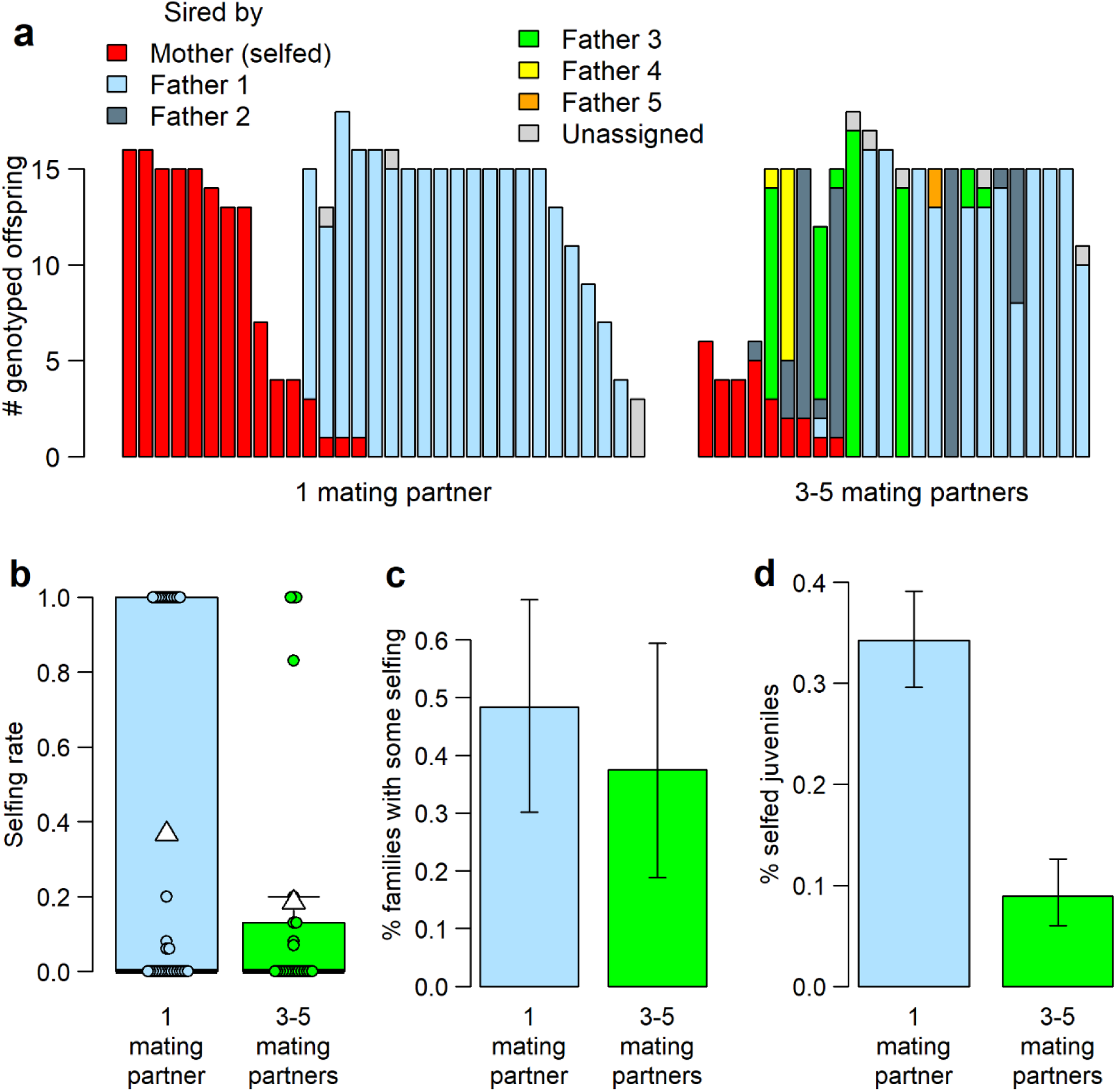
Experimental variation in mate availability did not affect individual-level selfing rates. Snails had either one mating opportunity with one partner, or three to five, each with a different partner. (a) Paternity distributions among the progenies of 56 F1 snails. Each vertical bar depicts one F2 family, with differently coloured sections corresponding to juveniles sired by different fathers. Father 1 refers to F1 snails’ first and father 5 to their last mating partner. Identical colours in different F2 families do not indicate identical father identities; all in all, F2 juveniles were sired by 68 unique fathers, with 10.5 ± 6.4 (mean ± SD) juveniles per father (min = 1, max = 30). Within pairing treatments, families are arranged according to selfing rate and number of genotyped juveniles. (b) Differences in mean selfing rates between once-(n = 31) and repeatedly paired snails (n = 24) were not statistically significant. White triangles on boxplots show group means. (c) The proportion of F2 families with non-zero selfing rates was similar among once-and repeatedly paired snails. (d) However, once-paired snails produced significantly more selfed juveniles overall. Error bars are 95% confidence intervals (79).

There was no evidence of selfing in either the F1 or P0 generation. All 274 F1 snails were outcrossed, *i.e.*, they possessed a non-maternal allele at one or more loci (mean: 4.6 ± 1.4 loci; Table S1). P0 mothers of F1 snails with genotyped offspring were likely outcrossed too: their genotypes did not show reduced heterozygosity (*F*_IS_ = 0.009, 95% CI: –0.010, 0.068; Table S1). P0 mother identity did not explain any variance in the probability that, post-isolation, F1 snails reproduced via selfing (Table 1).

**Table 1.** GLMM on the probability of selfing post-isolation. Results are shown for F1 snails (n = 55) with genotyped offspring that could be assigned to fathers. The response variable was the likelihood of F1 snails producing at least one selfed juvenile after having been paired (no/yes), modelled using binomial errors. Fixed effects were the pairing treatment (once-vs. repeatedly; test of hypothesis 1), the number of developed embryos produced in isolation (test of hypothesis 2), and the number of male (potential sperm donors) and female partners (potential sperm recipients) a snail copulated with. P0 mother identity was included as a random intercept; its effect was non-significant 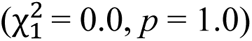. Results are provided on a logit scale. *β*: estimate, S.E.: standard error, *z*: *z*-value, *p*: *p*-value.

| Predictor | $\beta$ (S.E.) | $z$ | $p$ |
| --- | --- | --- | --- |
| Intercept | -0.22 (0.82) | -0.26 | 0.79 |
| Treatment (once- vs. repeatedly paired) | 1.75 (1.77) | 0.99 | 0.32 |
| # dev. embryos produced in isolation | 0.01 (0.01) | 2.24 | 0.0251 |
| # male mating partners | -0.82 (0.56) | -1.48 | 0.14 |
| # female mating partners | -0.19 (0.41) | -0.46 | 0.64 |

Multiple paternity was significantly more common among the offspring of repeatedly paired (45.8%) than once-paired snails (12.9%, Pearson’s Chi-squared test with Yates’ continuity correction: observed counts: 4, 27, 11, 13, expected counts: 8.5, 22.5, 6.5, 17.5, 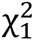 = 5.83, *p* = 0.0158; Fig. 1a); note that progenies of once-paired snails show multiple paternity when they contain both selfed and outcrossed offspring. On average, the progenies of repeatedly and once-paired snails had 1.7 ± 0.9 (max. = 4) and 1.1 ± 0.3 (max. = 2) fathers, respectively.

### Only subtle effect of mate availability on selfing

Mean selfing rates did not differ significantly among once-(0.37 ± 0.48) and repeatedly paired snails (0.19 ± 0.36; Wilcoxon rank sum test with continuity correction: *W* = 433.5, *p* = 0.25), nor did their variance (Levene’s test for homogeneity of variance: *F*_1,53_ = 2.44, *p* = 0.12; Fig. 1b). Repeated mating opportunities did not significantly reduce the likelihood of a snail producing at least some selfed offspring (Table 1), nor the proportion of families with non-zero selfing rates (once-paired: 48.4%, repeatedly paired: 37.5%, Pearson’s Chi-squared test: observed counts: 15, 16, 9, 15, expected counts: 13.5, 17.5, 10.5, 13.5, 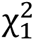 = 0.65, *p* = 0.42; Fig. 1c). We thus did not find strong support for hypothesis 1, which predicted that selfing is more common at low mate availability. However, as a group, once-paired snails produced significantly more selfed offspring than repeatedly paired snails: 34.2% vs. 8.9% (Pearson’s Chi-squared test: observed counts: 138, 265, 28, 286, expected counts: 93.3, 309.7, 72.7, 241.3, 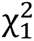 = 63.63, *p* < 0.0001; Fig. 1d). Nevertheless, the distribution of selfing rates was similarly bimodal among both treatment groups (Fig. 2a).

**Figure 2.**
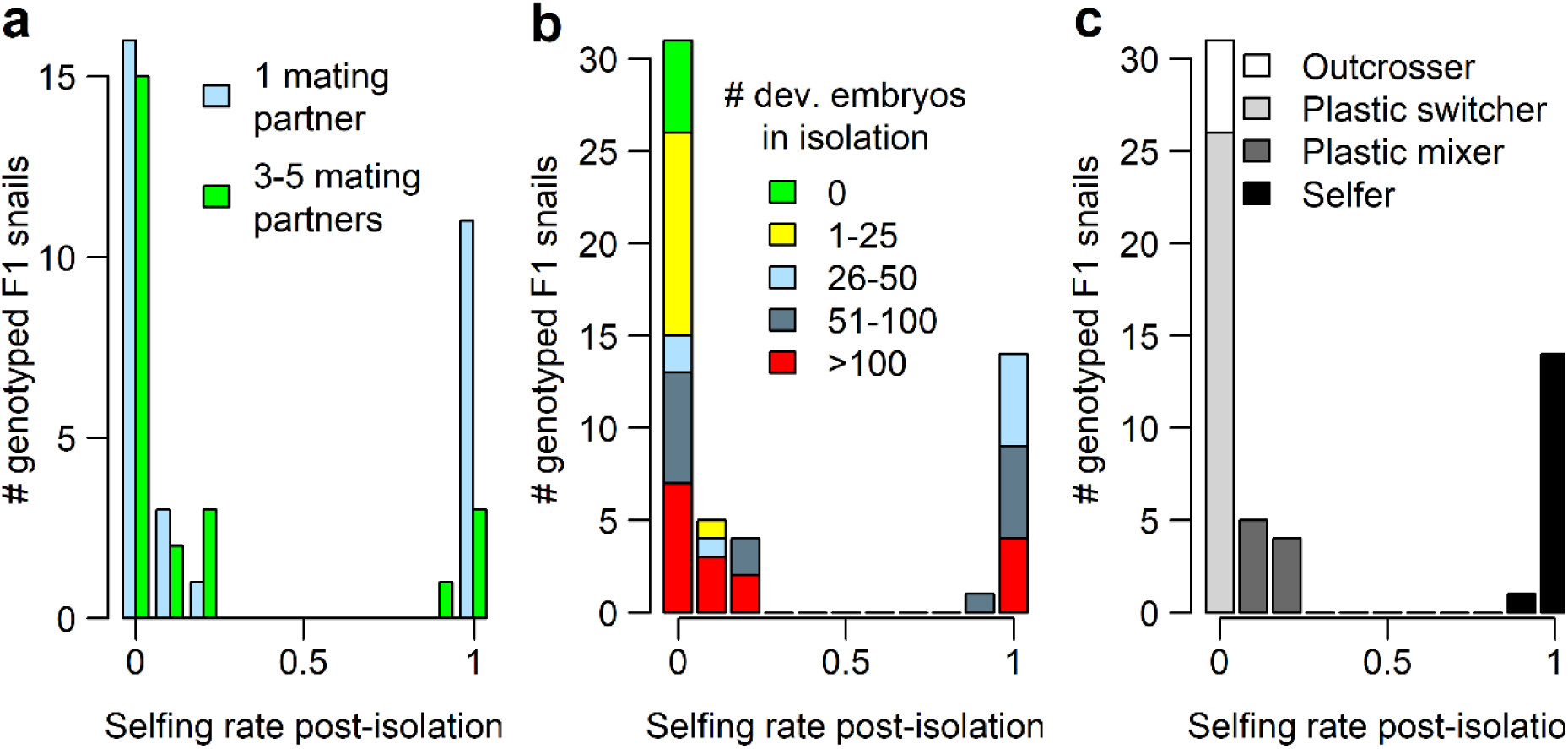
Variation in individual propensity for selfing. We ascertained the selfing propensity of snails with offspring that were successfully genotyped and assigned to a father (n = 55) based on the presence/absence of selfing in isolation and the post-isolation selfing rate. (a) The distribution of post-isolation selfing rates was bimodal among both once-and repeatedly paired snails. (b) Selfing in isolation predicted continued selfing: predominant selfing after mating trials had started only occurred in snails that had selfed successfully in isolation. (c) An a-posteriori categorisation revealed four classes of snails with differential selfing propensity: outcrossers that never reproduced via selfing, plastic switchers that selfed in isolation but outcrossed after being paired, plastic mixers that mixed selfing in isolation with a low level of selfing postisolation, and selfers that selfed fully or predominantly throughout their lives. These categories were very similar when based on the number of undeveloped embryos, or undeveloped and developed embryos combined (see Fig. S1).

### Selfing in isolation linked to selfing post-isolation

Predominant selfing after mating trials had started was restricted to snails that had selfed successfully in isolation, while snails without prior selfing exclusively outcrossed post-isolation (Fig. 2b). Consequently, selfing in isolation predicted continued selfing (Table 1); its effect remained statistically significant when an extremely prolific snail was excluded (*b* = 0.01, *z* = 2.04, *p* = 0.0418). We therefore conclude that our data support hypothesis 2, which posited that selfing post-isolation is more common in snails with a history of selfing.

### Individual-level variation in propensity for selfing

Snails varied in their propensity for selfing. For snails with estimated selfing rates (n = 55), we ascertained the selfing propensity from the presence/absence of selfing (*i.e.*, developed embryos) in isolation and the post-isolation selfing rate. This revealed four groups of snails: 9.1% “outcrossers” that never selfed, 47.3% “plastic switchers” that only selfed in isolation, 16.4% “plastic mixers” that combined selfing in isolation with predominant yet incomplete outcrossing post-isolation, and 27.3% “selfers” that selfed fully or mostly throughout (Fig. 2c). Most outcrossers did not produce any undeveloped embryos in isolation either. Consequently, their proportion (9.1%) was largely unchanged when based on the number of undeveloped (10.9%; Fig. S1a), or developed and undeveloped embryos (5.5%; Fig. S1b). Numbers of developed and undeveloped embryos were positively correlated across the whole dataset (Pearson’s product-moment correlation: *r* = 0.61, *t*_266_= 12.53, *p* < 0.0001; Fig. S1c).

The subsample with estimated selfing rate underrepresents snails that reproduced only before or only after mating trials started (Fig. S2). To estimate the frequency of selfing propensities more accurately, we thus included all female fertile snails; the selfing propensity of female infertile snails is unknown. We ascertained the apparent selfing propensity of snails without selfing rate estimates based on when they laid eggs and how they mated. Apparent outcrossers only reproduced post-isolation and mated as a female, apparent plastic snails reproduced both before and after being paired and mated as a female, and apparent selfers only reproduced in isolation, or reproduced in isolation and post-isolation yet never mated as a female. Across all female fertile snails (n = 124), this revealed 30.6% apparent outcrossers, 37.1% apparent plastic snails, and 32.3% apparent selfers

The pairing treatment did not influence the selfing propensity of snails with estimated selfing rates (Pearson’s Chi-squared test: observed counts: 11, 4, 15, 1, 4, 5, 11, 4, expected counts: 8.5, 5.1, 14.7, 2.8, 6.5, 3.9, 11.3, 2.2, 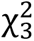 = 4.98, *p* = 0.17). However, when using the full dataset, selfing propensities differed among experimental groups (observed counts: 29, 26, 11, 11, 20, 27, expected counts: 21.3, 24.5, 20.2, 18.7, 21.5, 17.8, 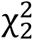 = 15.17, *p* = 0.0005). Low mate availability led to more apparent selfers (43.9% vs. 19.0%) and fewer apparent outcrossers (16.7% vs. 46.6%) yet did not affect apparent plastic snails (39.4% vs. 34.5%), lending some support to hypothesis 1.

### Reduced female LRS and increased offspring mortality in selfers

Selfing was associated with a significant decrease in female LRS (number of developed embryos) of 48.9% compared to plastic mixers and of 26.8% compared to plastic switchers (Fig. S3a, Table S3). On the one hand, this was because selfers produced significantly fewer eggs – 39.9% and 19.1% fewer than plastic mixers and switchers, respectively (Table S3). On the other hand, selfers laid more eggs that failed to develop (28.3%) than plastic mixers (16.2%); the difference to plastic switchers (21.4%) was non-significant (Fig. S3b, Table S3). Outcrossers were excluded because of low sample size (n = 5). Across all female fertile snails, female LRS in apparent selfers was not only reduced compared to apparent plastic snails (by 62.9%), but also compared to apparent outcrossers (by 52.0%; Fig. 3a, Table 2). Again, this was both due to a reduced fecundity (reductions by 57.0% and 44.9%, respectively) and increased proportion of undeveloped embryos in apparent selfers (36.8% vs. 25.4% and 28.2%, respectively; Fig. 3b, Table 2).

**Figure 3.**
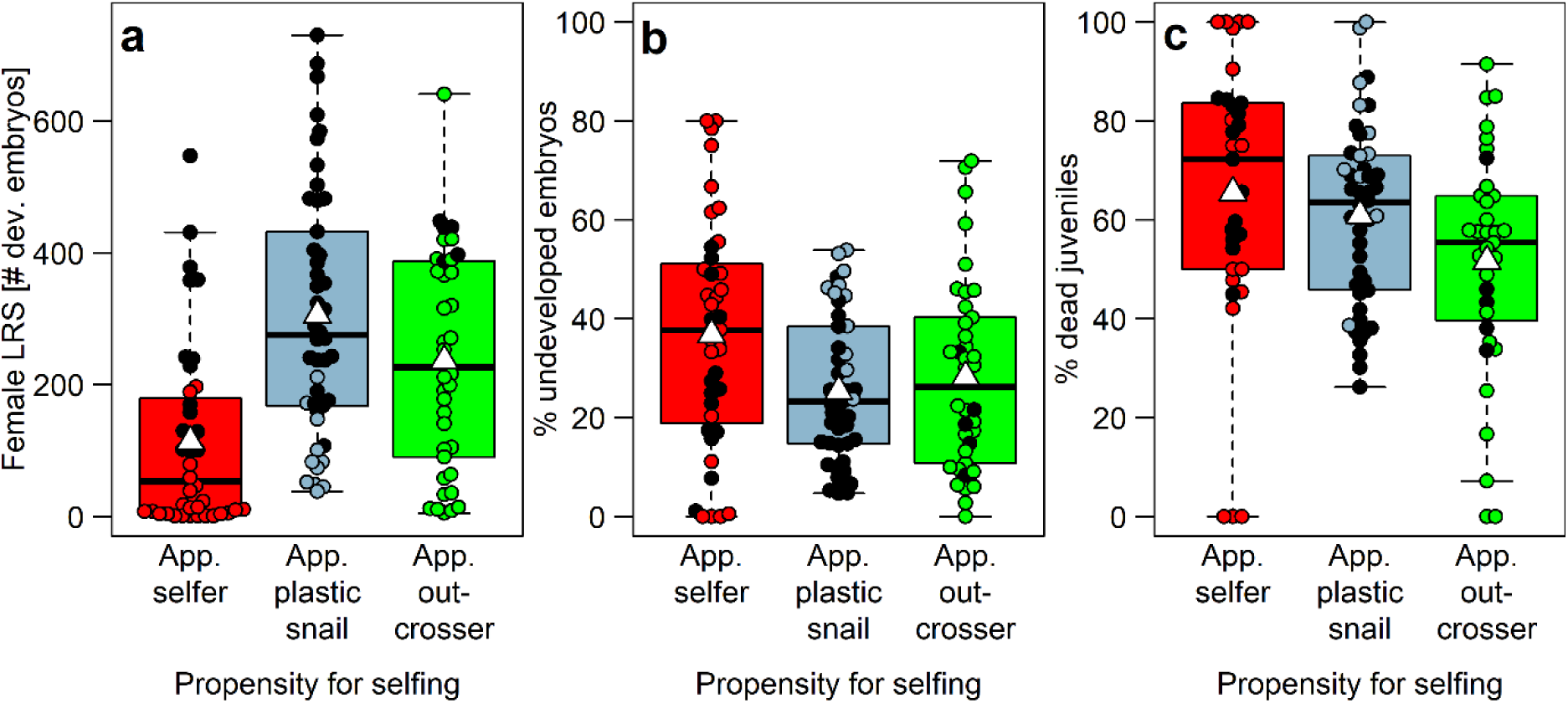
Reduced female LRS and increased offspring mortality in preferential selfers. Individuals with a strong propensity for selfing had the fewest developed embryos (a, n = 123), the highest proportion of embryos that failed to develop (b, n = 123), and the highest mortality rate among their juvenile offspring (c, n = 113). Shown are snails with (black circles) and without genotyped offspring (empty circles). We ascertained the apparent propensity for selfing of snails without genotyped offspring based on the timing of egg production and the mating behaviour. White triangles on boxplots show group means.

**Figure 4.**
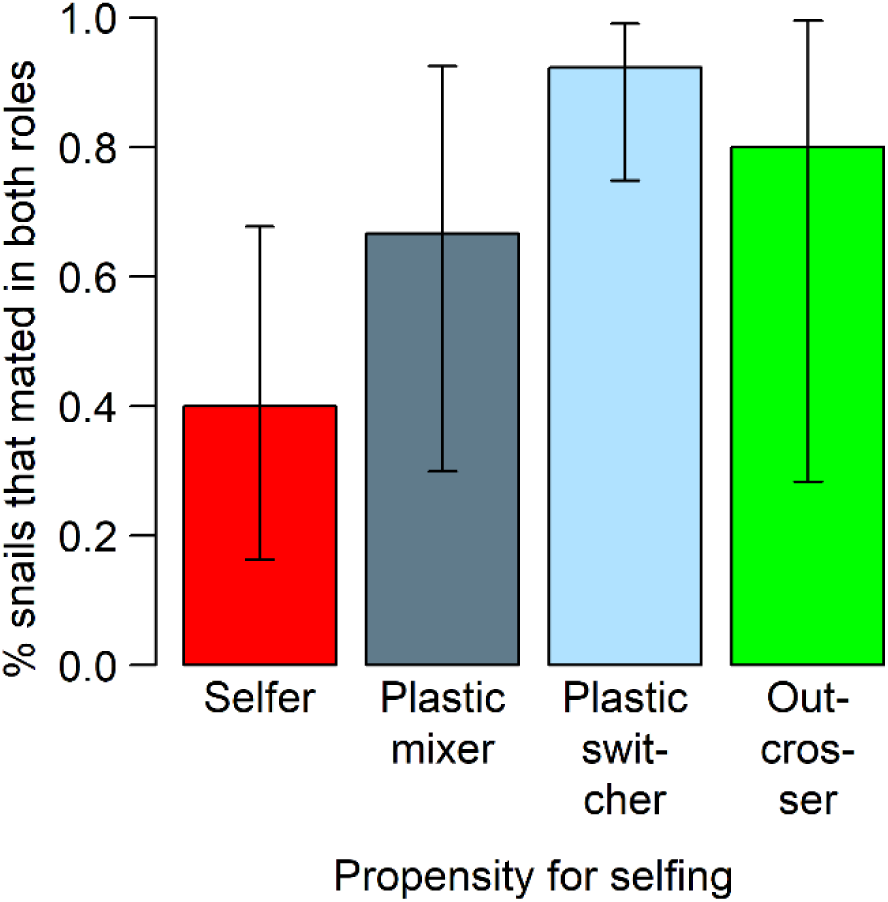
Poor correspondence between propensity for selfing and sexual drive. Although selfers (n = 15) were less likely to mate in both sexual roles than plastic switchers (n = 26), they showed considerable sexual activity, with 53.3% mating as a female and 80.0% as a male. Error bars are 95% confidence intervals (79).

**Table 2.** GLMMs on female LRS, its components, and juvenile mortality. Results are shown for four models using all female fertile F1 snails. Response variables were the lifetime number of developed embryos (female LRS) and eggs, the lifetime proportion of undeveloped embryos, and the proportion of developed embryos that died before they could be genotyped (*i.e.*, before reaching the juvenile age of 12.3 ± 0.5 weeks). For response variables 1 and 2, negative binomial errors were fitted, and results are provided on the log scale. For the two response variables that are proportions, Gaussian errors were fitted, as they did not deviate significantly from normality and as model assumptions were fulfilled. Fixed effects were propensity for selfing (reference level: apparent selfers) and body size (shell length). As random intercepts we included P0 mother identity (significant: 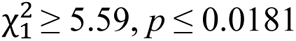, except for % dead juveniles: 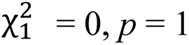) and, in all models except that of the number of eggs, pair identity on mating opportunity 1 (non-significant: 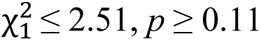). Although there were 124 female fertile F1 snails, sample size in models was reduced because one snail lacked data on body size and ten snails lacked developed embryos that were reared to the juvenile age. Results were very similar when computed for snails with estimated selfing rates only (Table S3). S.E.: standard error, PS.: propensity for selfing, app.: apparent.

| Response | Sample size | Predictor | Estimate (S.E.) | z-value | p-value |
| --- | --- | --- | --- | --- | --- |
| # dev. embryos (LRS) | 123 | Intercept | 2.59 (0.63) | 4.11 | <0.0001 |
|  |  | PS (app. plastic snail) | 1.08 (0.18) | 6.13 | <0.0001 |
|  |  | PS (app. outcrosser) | 0.89 (0.19) | 4.81 | <0.0001 |
|  |  | Body size | 0.14 (0.04) | 3.27 | 0.0011 |
| # eggs | 123 | Intercept | 2.75 (0.59) | 4.64 | <0.0001 |
|  |  | PS (app. plastic snail) | 1.02 (0.17) | 6.11 | <0.0001 |
|  |  | PS (app. outcrosser) | 0.83 (0.17) | 4.79 | <0.0001 |
|  |  | Body size | 0.15 (0.04) | 3.81 | 0.0001 |
| % undev. embryos | 123 | Intercept | 30.78 (14.87) | 2.07 | 0.0384 |
|  |  | PS (app. plastic snail) | -12.77 (3.80) | -3.36 | 0.0008 |
|  |  | PS (app. outcrosser) | -11.14 (3.92) | -2.84 | 0.0045 |
|  |  | Body size | 0.47 (1.07) | 0.44 | 0.66 |
| % dead juveniles | 113 | Intercept | 106.09 (19.51) | 5.44 | <0.0001 |
|  |  | PS (app. plastic snail) | -7.80 (4.94) | -1.58 | 0.11 |
|  |  | PS (app. outcrosser) | -16.69 (5.12) | -3.26 | 0.0011 |
|  |  | Body size | -2.71 (1.38) | -1.96 | 0.0496 |

Additionally, selfing resulted in inbreeding depression in juvenile survival. In snails with selfing rate estimates, the proportion of dead F2 juveniles was significantly higher among the progenies of selfers (69.5%) than among those of plastic switchers (53.4%); the difference to plastic mixers (64.5%) was non-significant (Fig. S2c, Table S3). Within all female fertile snails, apparent selfers had more dead juveniles (65.5%) than apparent outcrossers (51.6%), but not significantly more than apparent plastic snails (60.9%; Fig. 3c, Table 2).

These results were all corrected for effects of body size, P0 mother identity, and, where possible, pair identity on mating opportunity 1 (Table 2, Table S3).

### Mating as a female did not prevent selfing

A strong propensity for selfing was not linked to a lack of sexual drive. Neither the number of male (potential sperm donors) nor female mating partners (potential sperm recipients) was significantly associated with the likelihood of selfing post-isolation (Table 1). Many selfers (53.3%) copulated in the female role and so might have received allosperm. Four out of five selfers (80.0%) mated as a male, potentially siring outcrossed offspring. Of snails that selfed postisolation, 70.8% mated as a female.

However, the selfing propensity and sexual activity were not entirely independent. Significantly fewer selfers (40.0%) than plastic switchers (92.3%) copulated in both sexual roles (*b* = 2.88, *z* = 3.18, *p* = 0.0015; Fig. 4); the difference to plastic mixers was non-significant (66.7%, *b* = 1.12, *z* = 1.26, *p* = 0.21). Outcrossers (80.0%) were excluded from the model due to low sample size (n = 5, Fig. 4). Body size (*b* = –0.05, *z* = –0.24, *p* = 0.81) and P0 mother identity (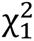 = 0.0, *p* = 1.00) did not affect the likelihood of mating in both sexual roles.

## Discussion

We found that propensities for selfing differed substantially within a snail population. Individual-level variation in selfing propensity has not received much attention in the mating-system literature, which has traditionally been focused on the level of species (5, 6, 7, 8) or populations (10, 13).

We controlled for several possible sources of error that could hamper the estimation of individual selfing propensities. Focal individuals were outcrossed, sexually mature, and paired with at least one size-matched partner that was not a close relative. The pairing treatments contrasted two biologically meaningful levels of mate availability. We estimated individual selfing rates post-isolation from progeny arrays, which are almost free of problematic assumptions (26, 63, 64). Our molecular markers were highly polymorphic, yielding reliable paternity assignments with sufficient statistical power (26). Nonetheless, it is important to note that we estimated “secondary” selfing rates, which might be biased if there is inor outbreeding depression prior to sampling. Here, we found inbreeding depression: juvenile mortality was highest when progenies were selfed, intermediate when mixed, and lowest when outcrossed. Most pre-sampling mortality thus likely affected selfed juveniles. Accordingly, there should be no bias in high selfing rates, while low selfing rates (0 < *s* < 0.21) and those of zero might be slightly too low. This does not change the high prevalence of self-fertility in our population (in fact, it underestimates it), nor the bimodal distribution of selfing rates, nor the substantial variation in selfing propensities among individuals.

The mating-system variation was cryptic in two ways. First, it only became apparent in the laboratory. By contrast, both direct and indirect selfing rate estimates of the ancestral field population were near-zero across three years and throughout the breeding season (26). Using the field results only, the population would have been deemed outcrossing and its considerable potential for selfing missed. Second, much variation would have been overlooked had we only recorded selfing during an isolation treatment, without estimating post-isolation selfing rates. This approach, used in pioneering earlier studies (e.g., 65, 66), merely reveals individuals’ ability to self but not their propensity for selfing, which must be assessed in the face of outcrossing opportunities and using genetic markers. Here we show just how variable the propensity is, comprising two types of plastic individuals, selfers, and outcrossers. Selfing-outcrossing thus is a gradient, not a dichotomy – also, and perhaps especially, at the individual level.

We found that variation in selfing propensity had a stable, environment-independent component: the better an individual was at selfing in isolation, the more likely it was to self post-isolation (hypothesis 2). In fact, those individuals that did not self while isolated invariably outcrossed thereafter. These pure outcrossers were numerous, representing nearly a third of female fertile snails. Notably, most of them did not produce any undeveloped embryos in isolation either, showing that they did not try to self and failed (perhaps due to early-acting inbreeding depression), but rather did not try at all. Furthermore, given the deliberately late start of mating trials, it is unlikely that many outcrossers would have begun to self if isolated even longer. These results hint at a genetic background to selfing propensities, which is in line with findings in other hermaphroditic animals (15, 22, 35, 36, 41). An aversion to selfing so strong as to risk reproductive failure when deprived of mates is so clearly maladaptive that a genetic self-incompatibility mechanism appears plausible. While common in plants (39, 40), such mechanisms have only recently been discovered in animals (41). Our results imply that a detailed investigation of the genetic basis of variation in self-compatibility would be worthwhile in this population.

By contrast, we found only limited support for a strong ecological component of selfing propensities: selfing post-isolation was not substantially more common under low mate availability (hypothesis 1). While selfing was indeed most prevalent when snails were isolated, it often persisted post-isolation. Moreover, pairing snails repeatedly rather than once had a surprisingly subtle effect, significantly affecting neither the mean selfing rate, the proportion of selfing individuals, nor the likelihood of selfing. This was despite large effect sizes and sufficient statistical power: doubling our sample size would not have rendered any of these effects significant (see R Markdown document). Instead, treatment effects were non-significant because variation among snails was large even within treatment groups. The only significant consequences of pairing snails repeatedly were a lower overall number of selfed offspring, showing that increased mate availability reduced selfing in high-fitness individuals, and an overabundance of apparent outcrossers and scarcity of apparent selfers when considering the full dataset.

Superficially, the subtlety of these effects contradicts ecological models of selfing, which assume an advantage and thus higher prevalence of selfing at low population density (43, 44). So far, empirical evidence for reproductive assurance in animals is mixed. While some support was found in androdioecious species such as the clam shrimp *Eulimnadia texana* (67) and the nematode *C. elegans* (68), studies of hermaphroditic snails struggled to find any. Selfing rates were not increased in low-density, disturbed, or temporary habitats (16, 19, 21, 31), neither the frequency nor duration of outcrossing depended on the length of a mating opportunity (45), and individuals did not adjust their waiting time to the perceived density of conspecifics (46). And yet, it is hard to imagine that the capacity for independent reproduction does not feature among the primary evolutionary drivers of selfing.

Our study offers a potential reconciliation between the reproductive assurance hypothesis and its limited empirical support in animals. First, reproductive assurance matters, but only in extreme situations. Consequently, there is an ascertainment bias: if a population is dense enough to be sampled, it might be too dense for selfing to be common. Second, it is important to understand that a population’s selfing rate depends on the distribution of individual selfing propensities among its members. As selfing propensities consist of an individual (*e.g.*, genotype) and environmental component (*e.g.*, population density), the relationship between density and population-level selfing rates is complicated. It depends on the relative frequency of individuals that can respond to density variation with increased or decreased selfing, *i.e.*, on the frequency of plastic individuals and selfers compared to outcrossers. Hence, population-level selfing rates can be low because the population mainly consists of outcrossers, or because density is high enough for outcrossing to occur even in plastic individuals and selfers. Interpreting high population-level selfing rates is perhaps easier, as they necessitate a sizeable proportion of self-fertile individuals. However, also in highly selfing populations density is difficult to predict, because the threshold density beneath which selfing occurs may be population- or species-specific.

These considerations might explain the apparent conflict between our field (near-zero selfing rate; 26) and laboratory results (widespread self-fertility). We think it likely that the high density of our field population prompts even selfers to reproduce through outcrossing. Consequently, the evident self-fertility of large parts of the population does not manifest under the current conditions. Interestingly, the commonness of selfing in the laboratory also among repeatedly paired snails suggests that, compared to their free-living conspecifics, these snails were still mate-limited. If so, then a future experiment with massively increased mate availability should reduce selfing to zero.

So, what maintains self-compatibility alleles in our study population, and why have they not gone to fixation? The cost of lacking self-compatibility is obvious – an occasional crash of population size will quickly wipe out any individuals without self-compatibility alleles. The benefits of lacking self-compatibility alleles, as demonstrated here, include a higher female LRS, resulting from increased fecundity and decreased embryo mortality. Also juvenile mortality was lower among outcrossed progenies. Hence, selfing did not only lead to inbreeding depression in embryo development, but also reduced juvenile survival. The presence of inbreeding depression in laboratory settings tallies with findings in other populations of *R. balthica* (69) and in a close relative (65, 66), and presumably represents a major obstacle for the spread of self-compatibility alleles. The reduced female lifetime fecundity of selfers (45% fewer eggs than outcrossers) is interesting as well. We previously found that selfed egg clutches were only half the size of (supposedly outcrossed) clutches laid after a female mating (49). Our study shows that potential compensation mechanisms of selfers, such as a faster clutch-laying rate, were – if at all present – clearly insufficient. Another benefit of lacking self-compatibility alleles might be an increased male LRS, particularly if selfers have sperm discounting (70) or reduced energy allocation to the male function (71). However, the strong sexual drive of selfers documented here (80% mated in the male role) suggests that reductions in siring success when selfing might be more modest. The ability to self while also acting as a paternal parent is a main requirement of another hypothesis for the evolution of selfing: the 50% transmission advantage of a selfer’s genes over those of an outcrosser (43, 72). If future work finds this ability to be common, then it provides a mechanism for maintaining self-compatibility alleles also at high population density.

Finally, it is worth mentioning the dangers of assuming that pairing individuals results in their offspring being outcrossed, as done in some early work (e.g., 65, 66). In our study, 44% of paired snails selfed at least partially, and 71% of snails that selfed post-isolation copulated as a female. Hence, outcrossing is neither guaranteed by pairing individuals once or repeatedly, nor by observing female copulations. Potential reasons for a failure to outcross include, *e.g.*, unsuccessful sperm transfer, cryptic female choice, sperm competition with autosperm, and genetic incompatibility between sperm and egg. Hence, to verify outcrossing events, genetic paternity analysis is unavoidable. Risks of failing to do so include underestimating the frequency and extent of selfing, and underestimating the strength of inbreeding in studies that compare the offspring of isolated parents (obligately selfed) with those of paired parents (purportedly outcrossed though potentially selfed; e.g., 48, 73, 74, 75). We here add to a growing number of studies that tested the assumption of (near-)exclusive outcrossing after pairing empirically, and found it occasionally confirmed (76), yet more often refuted (34, 54, 77, 78).

## Ethics

We performed our study in accordance with national laws.

## Data accessibility

Microsatellite markers: GenBank Accession nos. KX830983-KX830992. All data are available from Dryad X.

## Authors’ contributions

A.F. designed the study, performed experiments, carried out molecular work, and analysed the data. A.F. and A.S. drafted the manuscript, with input from J.J. All authors contributed to manuscript revision and gave final approval for publication.

## Supporting information

R Markdown Document

## Acknowledgements

We thank Hannele Penson, Pascal Reichlin, Oliver Subotic and David Tadres for help with laboratory experiments, and Natalie Sieber, Kirstin Kopp and Katri Seppälä for help with genotyping. DNA fragments were analysed for length polymorphisms using a 3730×l DNA Analyzer (Applied Biosystems) situated at the Genetic Diversity Centre (GDC), ETH Zurich.

