## Supplementary material for "Selfing-outcrossing as a gradient, not a dichotomy: propensity for selfing varies within a population of hermaphroditic animals": R Markdown Document

21 04 2023

### 1. Preparations

#### 1.1. Import data

```
# Paternity data of F2 snails
pat<-read.csv("Paternity_data.csv", header=T, sep=";")

# Survival data of F2 snails
sur<-read.csv("Survival_data_F2_snails.csv", header=T, sep=";")

# Data of all 274 F1 snails
sna<-read.csv("F1_snail_data.csv", header=T, sep=";")

# Design of mating trial F1 snails were subjected to
des<-read.csv("Design_of_mating_trial.csv", header=T, sep=";")
```

#### 1.2 Load required packages

```
library(beeswarm) # for scatter plots
library(car) # for Levene's Test
library(DHARMa) # for model diagnostics
library(glmmTMB) # for generalised linear mixed models
```

### 2. Figure 1

#### 2.1 Preparations for Figure 1a

```
a<-as.table(rbind(as.numeric(pat[2,2:ncol(pat)]), as.numeric(pat[3,2:ncol(pat
)])))
a2<-as.table(rbind(a, as.numeric(pat[4,2:ncol(pat)])))
a3<-as.table(rbind(a2, as.numeric(pat[5,2:ncol(pat)])))
a4<-as.table(rbind(a3, as.numeric(pat[6,2:ncol(pat)])))
a5<-as.table(rbind(a4, as.numeric(pat[7,2:ncol(pat)])))
a6<-as.table(rbind(a5, as.numeric(pat[8,2:ncol(pat)])))

dimnames(a6)<-list(response=c("Mother", "Father 1", "Father 2", "Father 3", "
Father 4", "Father 5", "Unassigned"))
```

#### 2.2 Figure 1a

```
#tiff(filename="Fig1a.tif", width=170, height=85, units="mm", res=300, points
ize=11)

par(mfrow=c(1,1))
par(mar=c(2,3,5.5,1), xpd=T)

barplot(a6,
    xlab="", ylab="", col=c("red", "lightskyblue1", "lightskyblue4", "green1"
,
                                                 "yellow1", "orange", "lightg
rey"), beside=FALSE,
```

```
    cex.axis=1, cex.lab=1, las=1, yaxt="n",
    names.arg=c(rep("\n", 56)),
    space=c(rep(0.2, 32), 4, rep(0.2, 23)))

title(ylab="# genotyped offspring", mgp=c(1.8,1,0), cex.lab=1)
axis(2, tick=T, at=seq(0,15,5), labels=seq(0,15,5), cex.axis=1, las=1, mgp=c(
2,0.75,0))

legend("topleft", legend=c("Mother (selfed)",rownames(a6)[2:3]), col=c("red",
"lightskyblue1", "lightskyblue4"),
fill=c("red", "lightskyblue1", "lightskyblue4"),
bty="n", cex=1, inset=c(0, -0.35), title="Sired by")

legend("top", legend=rownames(a6)[4:7], col=c("green1", "yellow1", "orange",
"lightgrey"),
fill=c("green1", "yellow1", "orange", "lightgrey"),
bty="n", cex=1, inset=c(0, -0.3))

axis(1, at=c(20, 58), labels=c("1 mating partner", "3-5 mating partners"), mg
p=c(2,0.5,0), cex.axis=1, lwd=0)

text(-7, 22, "a", cex=1.5, font=2)
```

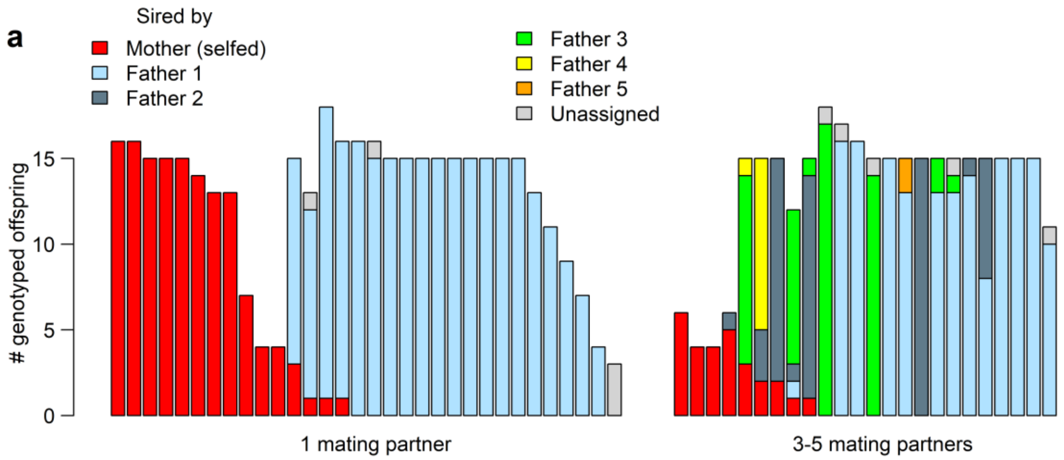

```
#dev.off()
```

#### 2.3 Preparations for Figure 1b-d

```
# transpose the dataframe
d<-as.data.frame(t(as.matrix(pat)))
colnames(d)<-d[1,]
d<-d[c(2:nrow(d)),]
rownames(d)<-c(1:nrow(d))

# ensure variables have the correct data type
d$mother<-as.factor(d$mother)
```

```
d$nr.juv.selfed<-as.numeric(d$nr.juv.selfed)
d$nr.juv.father1<-as.numeric(d$nr.juv.father1)
d$nr.juv.father2<-as.numeric(d$nr.juv.father2)
d$nr.juv.father3<-as.numeric(d$nr.juv.father3)
d$nr.juv.father4<-as.numeric(d$nr.juv.father4)
d$nr.juv.father5<-as.numeric(d$nr.juv.father5)
d$nr.juv.unassigned<-as.numeric(d$nr.juv.unassigned)
d$selfing.rate<-as.numeric(d$selfing.rate)
d$nr.paired<-as.numeric(d$nr.paired)
d$total.juv<-as.numeric(d$total.juv)
str(d)

## 'data.frame':    56 obs. of  11 variables:
##  $ mother           : Factor w/ 56 levels "11_4.1","14_3.7",..: 9 31 3 34
54 18 27 47 12 32 ...
##  $ nr.juv.selfed    : num  16 16 15 15 15 14 13 13 7 4 ...
##  $ nr.juv.father1   : num  0 0 0 0 0 0 0 0 0 0 ...
##  $ nr.juv.father2   : num  0 0 0 0 0 0 0 0 0 0 ...
##  $ nr.juv.father3   : num  0 0 0 0 0 0 0 0 0 0 ...
##  $ nr.juv.father4   : num  0 0 0 0 0 0 0 0 0 0 ...
##  $ nr.juv.father5   : num  0 0 0 0 0 0 0 0 0 0 ...
##  $ nr.juv.unassigned: num  0 0 0 0 0 0 0 0 0 0 ...
##  $ selfing.rate     : num  1 1 1 1 1 1 1 1 1 1 ...
##  $ nr.paired        : num  1 1 1 1 1 1 1 1 1 1 ...
##  $ total.juv        : num  16 16 15 15 15 14 13 13 7 4 ...

# add a column specifying the mating treatment
d$treat<-ifelse(d$nr.paired %in% 1, "once", "rep")
d$treat<-as.factor(d$treat)

# exclude the snail with unknown selfing rate
d<-d[is.na(d$selfing.rate)==F,]
nrow(d) # 55

## [1] 55

# compute proportions for Figs. 1c and 1d
d$nr.juv.outcr<-d$nr.juv.father1 + d$nr.juv.father2 + d$nr.juv.father3 +
d$nr.juv.father4 + d$nr.juv.father5
  # proportion of families with non-zero selfing rates
nrow(d[d$selfing.rate > 0.0 & d$treat %in% "once",]) # 15

## [1] 15

nrow(d[is.na(d$selfing.rate)==F & d$treat %in% "once",]) # 31

## [1] 31

propfamselfed.once<-15/31
propfamselfed.once # 0.48387

## [1] 0.483871
```

```
nrow(d[d$selfing.rate > 0.0 & d$treat %in% "rep",]) # 9

## [1] 9

nrow(d[is.na(d$selfing.rate)==F & d$treat %in% "rep",]) # 24

## [1] 24

propfamselfed.rep<-9/24
propfamselfed.rep # 0.375

## [1] 0.375

  # proportion of selfed juveniles
sum(d$nr.juv.selfed[d$treat %in% "once"]) # 138

## [1] 138

sum(d$nr.juv.outcr[d$treat %in% "once"])+sum(d$nr.juv.selfed[d$treat %in% "on
ce"]) # 403

## [1] 403

propselfed.once<-138/403
propselfed.once # 0.3424318

## [1] 0.3424318

sum(d$nr.juv.selfed[d$treat %in% "rep"]) # 28

## [1] 28

sum(d$nr.juv.outcr[d$treat %in% "rep"])+sum(d$nr.juv.selfed[d$treat %in% "rep
"]) # 314

## [1] 314

propselfed.rep<-28/314
propselfed.rep # 0.08917197

## [1] 0.08917197

# compute 95% confidence intervals for proportions shown in Figs. 1c and 1d
binom95CI<-function(x, n){
  df1_lower<-2*(n-x+1)
  df2_lower<-2*x
  df1_upper<-2*(x+1)
  df2_upper<-2*(n-x)
  F_lower<-qf(0.975, df1_lower, df2_lower)
  F_upper<-qf(0.975, df1_upper, df2_upper)
  CI_lower<-x/(x+(n-x+1)*F_lower)
  CI_upper<-((x+1)*F_upper)/(n-x+(x+1)*F_upper)
  return(c(CI_lower, CI_upper))
}
```

```
x<-c(15, 9)
n<-c(31, 24)
binom95CI(x, n)

## [1] 0.3015457 0.1879929 0.6693940 0.5940636

errorbars_fig1c<-binom95CI(x, n)

x<-c(138, 28)
n<-c(403, 314)
binom95CI(x, n)

## [1] 0.29616865 0.06007284 0.39102891 0.12629813

errorbars_fig1d<-binom95CI(x, n)
```

##### 2.4 Figures 1b-d

```
#tiff(filename="Fig1bcd.tif", width=170, height=85, units="mm", res=300,
#     pointsize=16)

par(mfrow=c(1,3))

# Fig. 1b
par(mar=c(3.5,3.5,2,0), xpd=T)
par(bty="n")
plot(d$selfing.rate~d$treat, las=1, xlab="", ylab="", col=c("lightskyblue1","
green1"),
     outlier.color=NA, cex.lab=1, cex.axis=1, xaxt="n", yaxt="n")
title(ylab="Selfing rate", mgp=c(2.2,1,0), cex.lab=1)
axis(1, tick=F, at=c(1, 2), labels=c("1\nmating\npartner", "3-5\nmating\npart
ners"),
     cex.axis=1, las=1, mgp=c(2,2.5,0))
axis(2, tick=T, at=seq(0,1,0.2), labels=c("0.0", "0.2", "0.4", "0.6", "0.8",
"1.0"),
     cex.axis=1, las=1, mgp=c(2,0.75,0))
beeswarm(d$selfing.rate~d$treat, pch=21, bg=c("lightskyblue1","green1"), cex=
1,
         spacing=0.3, add=T)
text(-0.15, 1.065, "b", cex=1.5, font=2)
points(tapply(d$selfing.rate, d$treat, mean, na.rm=T), pch=24, bg="white", ce
x=1.5)

# Fig. 1c
par(mar=c(3.5,3.5,2,1), xpd=T)
par(bty="n")
a<-barplot(height=c(propfamselfed.once, propfamselfed.rep),
        beside=TRUE, xlab="", col=c("lightskyblue1","green1"),
        ylab="", cex.axis=1, cex.lab=1, cex.names=1, las=1,
        mgp=c(3.5,1,0), space=c(0.3), xaxt="n", yaxt="n", ylim=c(0, 0.7))
```

```
title(ylab="% families with some selfing", mgp=c(2.2,1,0), cex.lab=1)
axis(1, tick=F, at=c(0.8, 2.1), labels=c("1\nmating\npartner", "3-5\nmating\n
partners"),
     cex.axis=1, las=1, mgp=c(2,2.5,0))
axis(2, tick=T, at=seq(0,0.6, 0.1), labels=c("0.0", "0.1", "0.2", "0.3", "0.4
", "0.5", "0.6"),
     cex.axis=1, las=1, mgp=c(2,0.75,0))
text(-0.55, 0.71, "c", cex=1.5, font=2)
segments(a, errorbars_fig1c[1:2], a, errorbars_fig1c[3:4], lwd=1)
arrows(a, errorbars_fig1c[1:2], a, errorbars_fig1c[3:4], lwd=1, angle=90, cod
e=3, length=0.05)

# Fig. 1d
par(mar=c(3.5,3.5,2,0), xpd=T)
par(bty="n")
a<-barplot(height=c(propselfed.once, propselfed.rep),
        beside=TRUE, xlab="", col=c("lightskyblue1","green1"),
        ylab="", cex.axis=1, cex.lab=1, cex.names=1, las=1,
        mgp=c(3.5,1,0), space=c(0.3), xaxt="n", yaxt="n", ylim=c(0, 0.4))

title(ylab="% selfed juveniles", mgp=c(2.2,1,0), cex.lab=1)
axis(1, tick=F, at=c(0.8, 2.1), labels=c("1\nmating\npartner", "3-5\nmating\n
partners"),
     cex.axis=1, las=1, mgp=c(2,2.5,0))
axis(2, tick=T, at=seq(0,0.4, 0.1), labels=c("0.0", "0.1", "0.2", "0.3", "0.4
"),
     cex.axis=1, las=1, mgp=c(2,0.75,0))
text(-0.55, 0.41, "d", cex=1.5, font=2)
segments(a, errorbars_fig1d[1:2], a, errorbars_fig1d[3:4], lwd=1)
arrows(a, errorbars_fig1d[1:2], a, errorbars_fig1d[3:4], lwd=1, angle=90, cod
e=3, length=0.05)
```

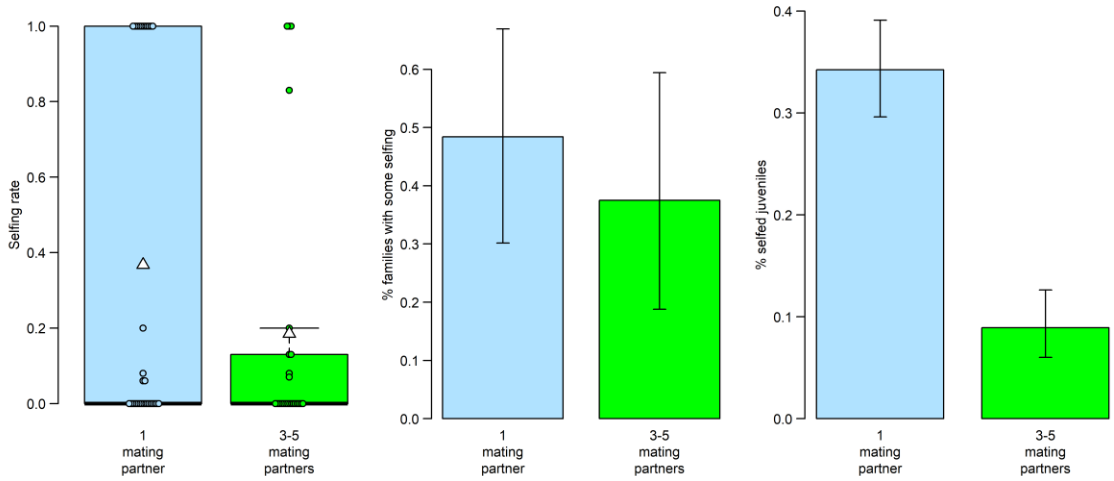

```
#dev.off()
```

### 3. Extracting numbers for Methods

#### 3.1 Calculating the number of P0 mothers and sibling groups among F1 snails

```
# for all 274 F1 snails subjected to mating trials
P0mothers.all<-sna$P0mother
length(unique(P0mothers.all)) # 38

## [1] 38

sibgroups.all<-sna$clutchID
length(unique(sibgroups.all)) # 108

## [1] 108

# for the subset of 56 snails with estimated selfing rates
P0mothers.sub<-sna$P0mother[sna$nr.genotyped>0 & sna$exclude %in% "no"]
length(unique(P0mothers.sub)) # 22

## [1] 22

sibgroups.sub<-sna$clutchID[sna$nr.genotyped>0 & sna$exclude %in% "no"]
length(unique(sibgroups.sub)) # 38

## [1] 38
```

#### 3.2 Calculating the number of F1 snails in both types of mating trials

```
# all F1 snails subjected to mating trials and paired also to unrelated snail
s
nrow(sna[sna$group %in% "paired.once" & sna$exclude %in% "no",]) # 131

## [1] 131

nrow(sna[sna$group %in% "paired.repeatedly" & sna$exclude %in% "no",]) # 137

## [1] 137

# all F1 snails with estimated selfing rates and paired also to unrelated sna
ils
nrow(sna[sna$group %in% "paired.once" & sna$nr.genotyped>0 & sna$exclude %in%
"no",]) # 32

## [1] 32

nrow(sna[sna$group %in% "paired.repeatedly" & sna$nr.genotyped>0 & sna$exclud
e %in% "no",]) # 24

## [1] 24
```

#### 3.3 Calculating the number of genotyped juveniles per family

```
# for snails with 15 or more genotyped juveniles
nrgen15ormore<-sna$nr.genotyped[sna$nr.genotyped>14 & sna$exclude %in% "no"]
length(nrgen15ormore) # 40 families
```

```
## [1] 40

mean(nrgen15ormore) # 15.975

## [1] 15.975

sd(nrgen15ormore) # 1.874081

## [1] 1.874081

min(nrgen15ormore) # 15

## [1] 15

max(nrgen15ormore) # 22

## [1] 22

# for snails with fewer than 15 genotyped juveniles
nrgen14orfewer<-sna$nr.genotyped[sna$nr.genotyped<15 & sna$nr.genotyped>0 & s
na$exclude %in% "no"]
length(nrgen14orfewer) # 16 families

## [1] 16

mean(nrgen14orfewer) # 8.0625

## [1] 8.0625

sd(nrgen14orfewer) # 3.623419

## [1] 3.623419

min(nrgen14orfewer) # 3

## [1] 3

max(nrgen14orfewer) # 14

## [1] 14

# total number of genotyped juveniles (without those from the snail only pair
ed with a sibling)
nrgen<-sna$nr.genotyped[sna$exclude %in% "no"]
sum(nrgen) # 768

## [1] 768
```

##### 3.4 Calculating the number of successfully genotyped juveniles per family

```
# number of successfully genotyped juveniles per family
nrsuccgen<-sna$nr.succ.genotyped[sna$nr.succ.genotyped>0 & sna$exclude %in% "
no"]
length(nrsuccgen) # 56 families
```

```
## [1] 56

mean(nrsuccgen) # 12.98214

## [1] 12.98214

sd(nrsuccgen) # 4.118654

## [1] 4.118654

min(nrsuccgen) # 3

## [1] 3

max(nrsuccgen) # 18

## [1] 18

# total number of successfully genotyped juveniles (without those from the sn
ail only paired # with a sibling)
sum(nrsuccgen) # 727

## [1] 727

# number and proportion of unsuccessfully genotyped juveniles
768-727 # 41

## [1] 41

41/768 # 0.05338542

## [1] 0.05338542
```

##### 3.5 Calculating the number of assigned juveniles per family

```
# number of juveniles per family that were assigned to a father
nrass<-sna$nr.assigned[sna$nr.assigned>0 & sna$exclude %in% "no"]
length(nrass) # 55 families

## [1] 55

mean(nrass) # 13.03636

## [1] 13.03636

sd(nrass) # 3.877588

## [1] 3.877588

min(nrass) # 4

## [1] 4

max(nrass) # 18

## [1] 18
```

```
# total number of assigned juveniles (without those from the snail only paire
d # with a sibling)
sum(nrass) # 717

## [1] 717

# number of unassigned and and proportion of assigned juveniles
727-717 # 10

## [1] 10

717/727 # 0.9862448

## [1] 0.9862448
```

### 4. Multiple paternity

#### 4.1 Estimating frequency and magnitude of multiple paternity

```
# Count number of fathers per family
for(i in 1:nrow(d)){
  x<-as.vector(d[i,c("nr.juv.selfed", "nr.juv.father1", "nr.juv.father2",
                     "nr.juv.father3", "nr.juv.father4", "nr.juv.father5")])
  d$nr.fathers[i]<-length(x[x!=0])
  }
d[c(50:55),c(1:7, 14)] # control - ok

##     mother nr.juv.selfed nr.juv.father1 nr.juv.father2 nr.juv.father3
## 51 40_3.16             0             14              1              0
## 52  45_2.3             0              8              7              0
## 53  51_3.2             0             15              0              0
## 54 52_4b.4             0             15              0              0
## 55 70_3.20             0             15              0              0
## 56  70_1.6             0             10              0              0
##    nr.juv.father4 nr.juv.father5 nr.fathers
## 51              0              0          2
## 52              0              0          2
## 53              0              0          1
## 54              0              0          1
## 55              0              0          1
## 56              0              0          1

# Calculate proportion of families with multiple paternity in treatment group
s
table(d$nr.fathers, d$treat)

##
##     once rep
##   1   27  13
##   2    4   7
```

```
##   3    0   3
##   4    0   1

4/31 # 0.1290323 for once-paired snails

## [1] 0.1290323

11/24 # 0.4583333 for repeatedly-paired snails

## [1] 0.4583333

# Chi-squared test for proportion of families with multiple paternity
# test with Yates' continuity correction as one cell frequency only five
row.names.mat1<-c("multpat", "nomultpat")
col.names.mat1<-c("once", "rep")
mat1<-matrix(data=c(4,27,11, 13), nrow=2, ncol=2, byrow=FALSE, dimnames=list(
row.names.mat1, col.names.mat1))
chisq.test(mat1, correct=T) # X-squared = 5.8285, df = 1, p-value = 0.01577

##
##  Pearson's Chi-squared test with Yates' continuity correction
##
## data:  mat1
## X-squared = 5.8285, df = 1, p-value = 0.01577

chisq.test(mat1, correct=T)$expected # expected counts

##                once       rep
## multpat    8.454545  6.545455
## nomultpat 22.545455 17.454545

# Calculate mean and maximum number of fathers
tapply(d$nr.fathers, d$treat, mean, na.rm=T) # 1.129032 (once), 1.666667 (rep
)

##     once      rep
## 1.129032 1.666667

tapply(d$nr.fathers, d$treat, sd, na.rm=T) # 0.3407771 (once), 0.8681147 (rep
)

##      once       rep
## 0.3407771 0.8681147

tapply(d$nr.fathers, d$treat, max, na.rm=T) # 2 (once), 4 (rep)

## once  rep
##    2    4
```

### 5. Effect of mate availability on selfing

#### 5.1 Effect on proportion of families with non-zero selfing rates

```
# How many families contained selfed offspring?
table(d$selfing.rate) # 2+1+2+2+2+1+14 = 24 families out of 55, i.e. 43.6%

##
##    0 0.06 0.07 0.08 0.13  0.2 0.83    1
##   31    2    1    2    2    2    1   14

# How many families contained few (max. 20%) vs. many (min 80%) selfed offspr
ing?
table(d$selfing.rate) # 9 few, 15 many

##
##    0 0.06 0.07 0.08 0.13  0.2 0.83    1
##   31    2    1    2    2    2    1   14

# How many families were selfed among once- and repeatedly paired snails?
table(d$selfing.rate, d$treat)

##
##         once rep
##   0       16  15
##   0.06     2   0
##   0.07     0   1
##   0.08     1   1
##   0.13     0   2
##   0.2      1   1
##   0.83     0   1
##   1       11   3

15/(15+16) # 0.483871 (once)

## [1] 0.483871

9/(9+15) # 0.375 (repeatedly)

## [1] 0.375

# Chi-squared test for proportion families with non-zero selfing rates
# test without Yates' continuity correction, as all cell frequencies above fi
ve
row.names.mat2<-c("selfing", "noselfing")
col.names.mat2<-c("once", "rep")
mat2<-matrix(data=c(15, 16, 9, 15), nrow=2, ncol=2, byrow=FALSE, dimnames=lis
t(row.names.mat2, col.names.mat2))
chisq.test(mat2, correct=F) # X-squared = 0.65191, df = 1, p-value = 0.4194

##
##  Pearson's Chi-squared test
##
```

```
## data:  mat2
## X-squared = 0.65191, df = 1, p-value = 0.4194
```

```
chisq.test(mat2)$expected # expected counts
```

```
##                 once      rep
## selfing   13.52727 10.47273
## noselfing 17.47273 13.52727
```

```
# Test whether doubling the sample size turns the treatment effect significan
t
mat2double<-matrix(data=c(30, 32, 18, 30), nrow=2, ncol=2, byrow=FALSE, dimna
mes=list(row.names.mat2, col.names.mat2))
chisq.test(mat2double, correct=F) # X-squared = 1.3038, df = 1, p-value = 0.2
535
```

```
##
##  Pearson's Chi-squared test
##
## data:  mat2double
## X-squared = 1.3038, df = 1, p-value = 0.2535
```

##### 5.2 Effect on mean selfing rates and their variance

```
# Mean selfing rate for once- and repeatedly paired snails
tapply(d$selfing.rate, d$treat, mean, na.rm=T)
```

```
##      once       rep
## 0.3677419 0.1850000
```

```
#      once       rep
# 0.3677419 0.1850000
tapply(d$selfing.rate, d$treat, sd, na.rm=T)
```

```
##      once       rep
## 0.4782378 0.3582931
```

```
#      once       rep
# 0.4782378 0.3582931

wilcox.test(d$selfing.rate[d$treat %in% "once"],
       d$selfing.rate[d$treat %in% "rep"])
```

```
## Warning in wilcox.test.default(d$selfing.rate[d$treat %in% "once"],
## d$selfing.rate[d$treat %in% : cannot compute exact p-value with ties
```

```
##
##  Wilcoxon rank sum test with continuity correction
##
## data:  d$selfing.rate[d$treat %in% "once"] and d$selfing.rate[d$treat %in%
"rep"]
## W = 433.5, p-value = 0.2484
## alternative hypothesis: true location shift is not equal to 0
```

```
# W = 433.5, p-value = 0.2484

# Test whether doubling the sample size turns the treatment effect significan
t
d.double<-rbind(d, d)
wilcox.test(d.double$selfing.rate[d.double$treat %in% "once"],
       d.double$selfing.rate[d.double$treat %in% "rep"])

##
##  Wilcoxon rank sum test with continuity correction
##
## data:  d.double$selfing.rate[d.double$treat %in% "once"] and d.double$self
ing.rate[d.double$treat %in% "rep"]
## W = 1734, p-value = 0.09897
## alternative hypothesis: true location shift is not equal to 0

# W = 1734, p-value = 0.09897

# Variance in selfing rate within once- and repeatedly paired snails
tapply(d$selfing.rate, d$treat, var, na.rm=T) # 0.2287114 (once), 0.1283739 (
rep)

##      once       rep
## 0.2287114 0.1283739

# Testing whether this difference is statistically significant
fligner.test(d$selfing.rate~d$treat) # Fligner-Killeen:med chi-squared = 1.74
51, df = 1, p-value = 0.1865

##
##  Fligner-Killeen test of homogeneity of variances
##
## data:  d$selfing.rate by d$treat
## Fligner-Killeen:med chi-squared = 1.7451, df = 1, p-value = 0.1865

leveneTest(d$selfing.rate~d$treat) # df = 1, 53, F-value = 2.4396, p-value =
0.1243

## Levene's Test for Homogeneity of Variance (center = median)
##       Df F value Pr(>F)
## group  1  2.4396 0.1243
##       53
```

##### 5.3 Effect on number of selfed juveniles

```
# How many juveniles were selfed?
tapply(d$nr.juv.selfed, d$treat, sum) # 138 (once), 28 (rep)

## once  rep
##  138   28

tapply(d$nr.juv.outcr, d$treat, sum) # 265 (once), 286 (rep)
```

```
## once  rep
##  265  286

propselfed.once<-138/(138+265) # 0.3424318
propselfed.once

## [1] 0.3424318

propselfed.rep<-28/(28+286) # 0.08917197
propselfed.rep

## [1] 0.08917197

propselfed<-(138+28)/(138+265+28+286) # 0.2315202, i.e., 166/717 (all)
propselfed

## [1] 0.2315202

# Chi-squared test for proportion of selfed juveniles
# test without Yates' continuity correction, as all cell frequencies above fi
ve
row.names.mat3<-c("selfed", "outcrossed")
col.names.mat3<-c("once", "rep")
mat3<-matrix(data=c(138, 265, 28, 286), nrow=2, ncol=2, byrow=FALSE, dimnames
=list(row.names.mat3, col.names.mat3))
chisq.test(mat3, correct=F) # X-squared = 63.625, df = 1, p-value = 1.505e-15

##
##  Pearson's Chi-squared test
##
## data:  mat3
## X-squared = 63.625, df = 1, p-value = 1.505e-15

chisq.test(mat3, correct=F)$expected # expected counts

##                 once       rep
## selfed      93.30265  72.69735
## outcrossed 309.69735 241.30265
```

### 6. Model 1: What influences the probability of selfing post-isolation?

#### 6.1 Preparations

```
# merge paternity and F1 snail data
s0<-sna[sna$snailID %in% d$mother,]
nrow(s0) # 55 control - ok

## [1] 55

colnames(d)[1]<-"snailID"
d1<-merge(d, s0, by="snailID")
nrow(d1) # 55 control - ok
```

```
## [1] 55

# merge paternity and F2 survival data
j1<-sur[sur$Mother %in% d$snailID,]
nrow(j1) # 55 control - ok

## [1] 55

colnames(j1)[1]<-"snailID"
d2<-merge(d1, j1, by="snailID")
nrow(d2) # 55 control - ok

## [1] 55

# add pair identity on mating opportunity 1 (from design of mating trial data
file)
partID<-des[,c("snailID", "pairID")]
d2<-merge(d2, partID, by="snailID", all.x=T)
nrow(d2) # 55 control - ok

## [1] 55

# rename columns with total number of surviving pre- and postmating F2 juveni
les
colnames(d2)[colnames(d2)=="nr.premating.snails"]<-"total.surv.juv.pre"
colnames(d2)[colnames(d2)=="nr.postmating.snails"]<-"total.surv.juv.post"

# add column with selfing in isolation yes/no
d2$self.isol<-ifelse(d2$total.nr.dev.embryos.pre > 0, "yes", "no")
table(d2$self.isol, d2$total.nr.dev.embryos.pre)

##
##        0 1 2 3 4 5 9 20 21 27 31 32 35 41 44 49 51 52 69 71 73 75 80 82 87
93 95
##    no  5 0 0 0 0 0 0  0  0  0  0  0  0  0  0  0  0  0  0  0  0  0  0  0  0
0  0
##    yes 0 4 2 1 1 1 1  1  1  1  1  1  1  1  1  2  1  1  1  2  1  1  1  1  1
1  1
##
##        96 107 111 117 126 132 138 141 142 143 153 156 171 182 210 329
##    no   0   0   0   0   0   0   0   0   0   0   0   0   0   0   0   0
##    yes  2   1   1   1   1   1   2   1   1   1   1   1   1   1   1   1

# add column with selfing post-isolation yes/no
d2$selfing<-ifelse(d2$nr.juv.selfed > 0, "yes", "no")
table(d2$selfing, d2$nr.juv.selfed) # control - ok

##
##         0  1  2  3  4  5  6  7 13 14 15 16
##    no  31  0  0  0  0  0  0  0  0  0  0  0
##    yes  0  5  2  2  4  1  1  1  2  1  3  2
```

```
d2$selfing<-as.factor(d2$selfing)

# make columns "P0mother" and "pairID" a factor
d2$P0mother<-as.factor(d2$P0mother)
d2$pairID<-as.factor(d2$pairID)

# add column with nr of male mating partners
d2$nr.male.partners<-d2$fem.matings.with.x.partners

# add column with nr of female mating partners
d2$nr.fem.partners<-d2$male.matings.with.x.partners
```

##### 6.2 Model 1: Probability of selfing post-isolation (Table 1)

```
m1<-glmmTMB(selfing~treat + total.nr.dev.embryos.pre + nr.male.partners
            + nr.fem.partners + (1|P0mother), data=d2, family=binomial)
summary(m1)
```

```
##  Family: binomial  ( logit )
## Formula:
## selfing ~ treat + total.nr.dev.embryos.pre + nr.male.partners +
##     nr.fem.partners + (1 | P0mother)
## Data: d2
##
##      AIC      BIC   logLik deviance df.resid
##     76.6     88.6    -32.3     64.6       49
##
## Random effects:
##
## Conditional model:
##  Groups   Name        Variance  Std.Dev.
##  P0mother (Intercept) 1.403e-09 3.745e-05
## Number of obs: 55, groups:  P0mother, 22
##
## Conditional model:
##                           Estimate Std. Error z value Pr(>|z|)
## (Intercept)              -0.216077   0.817550  -0.264   0.7915
## treatrep                  1.753554   1.766236   0.993   0.3208
## total.nr.dev.embryos.pre  0.011870   0.005298   2.240   0.0251 *
## nr.male.partners         -0.823713   0.558230  -1.476   0.1401
## nr.fem.partners          -0.189071   0.409130  -0.462   0.6440
## ---
## Signif. codes:  0 '***' 0.001 '**' 0.01 '*' 0.05 '.' 0.1 ' ' 1
```

```
summary(m1)$coef
```

```
## $cond
##                             Estimate  Std. Error    z value   Pr(>|z|)
## (Intercept)              -0.21607749 0.817549815 -0.2642989 0.79154963
## treatrep                  1.75355437 1.766236159  0.9928199 0.32079774
## total.nr.dev.embryos.pre  0.01186999 0.005298273  2.2403510 0.02506814
```

```
## nr.male.partners          -0.82371258 0.558229596 -1.4755803 0.14005661
## nr.fem.partners           -0.18907069 0.409129829 -0.4621288 0.64398893
##
## $zi
## NULL
##
## $disp
## NULL
```

```
# Diagnostic plot
res_m1<-resid(m1)
fitted_m1<-fitted(m1)
plot(res_m1~fitted_m1, las=1, xlab="Fitted values", ylab="Residuals",
     cex.lab=1.6, cex.axis=1.6, cex=1.6)
```

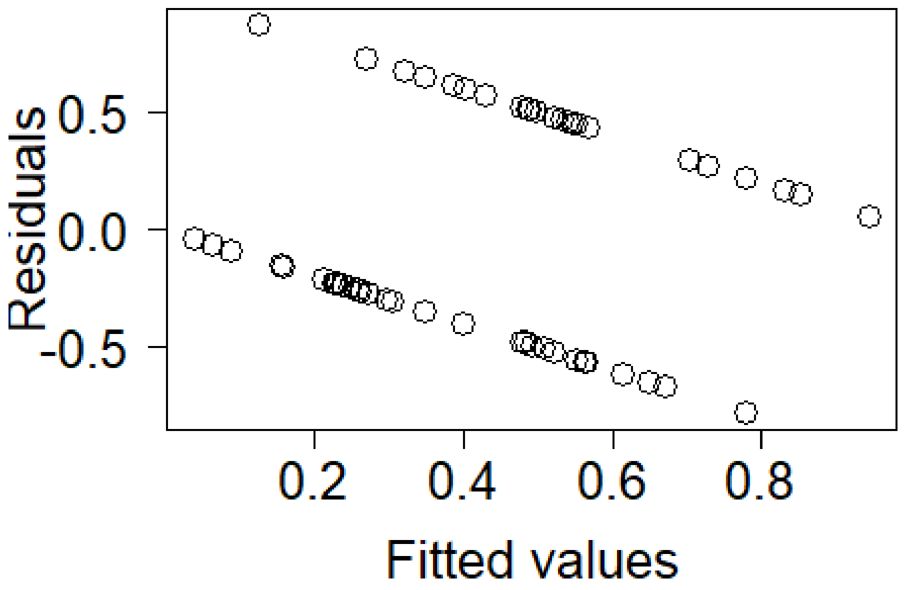

```
# Test the model fit using package DHARMa
  # simulate residuals
simres_m1<-simulateResiduals(fittedModel=m1, plot=F)
simres_m1all<-residuals(simres_m1)
  # make the plots
plot(simres_m1)
```

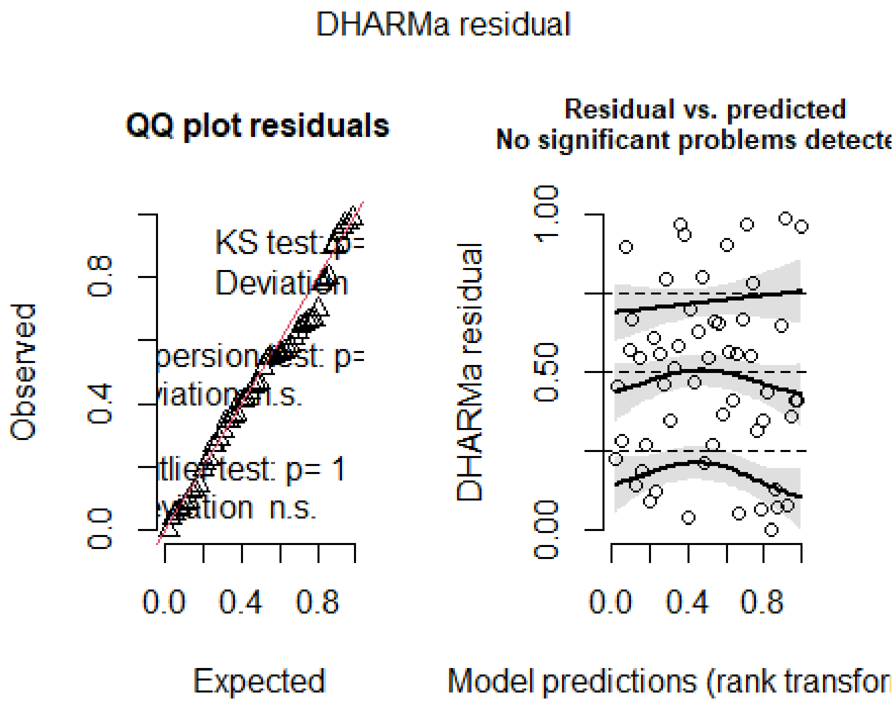

```
# Significance of random effects: P0mother
m1b<-glmmTMB(selfing~treat + total.nr.dev.embryos.pre + nr.male.partners
            + nr.fem.partners, data=d2, family=binomial)
anova(m1, m1b)

## Data: d2
## Models:
## m1b: selfing ~ treat + total.nr.dev.embryos.pre + nr.male.partners + , zi=
~0, disp=~1
## m1b:     nr.fem.partners, zi=~0, disp=~1
## m1: selfing ~ treat + total.nr.dev.embryos.pre + nr.male.partners + , zi=~
0, disp=~1
## m1:     nr.fem.partners + (1 | P0mother), zi=~0, disp=~1
##      Df    AIC    BIC  logLik deviance Chisq Chi Df Pr(>Chisq)
## m1b   5 74.563 84.600 -32.281   64.563
## m1    6 76.563 88.607 -32.281   64.563     0      1          1

# Model without the outlier that produced >300 developed embryos in isolation
d2nooutlier<-d2[d2$total.nr.dev.embryos.pre < 300,]
nrow(d2nooutlier) # 54 - ok

## [1] 54

m1c<-glmmTMB(selfing~treat + total.nr.dev.embryos.pre + nr.male.partners
            + nr.fem.partners + (1|P0mother), data=d2nooutlier, family=binomi
al)
summary(m1c) # still significant effect of selfing in isolation
```

```
##  Family: binomial  ( logit )
## Formula:
## selfing ~ treat + total.nr.dev.embryos.pre + nr.male.partners +
##     nr.fem.partners + (1 | P0mother)
## Data: d2nooutlier
##
##      AIC      BIC   logLik deviance df.resid
##     76.4     88.4    -32.2     64.4       48
##
## Random effects:
##
## Conditional model:
##  Groups   Name        Variance  Std.Dev.
##  P0mother (Intercept) 1.638e-09 4.047e-05
## Number of obs: 54, groups:  P0mother, 22
##
## Conditional model:
##                          Estimate Std. Error z value Pr(>|z|)
## (Intercept)             -0.169267   0.835334  -0.203   0.8394
## treatrep                 1.777846   1.764822   1.007   0.3138
## total.nr.dev.embryos.pre  0.011381   0.005592   2.035   0.0418 *
## nr.male.partners        -0.817808   0.556783  -1.469   0.1419
## nr.fem.partners         -0.208806   0.415626  -0.502   0.6154
## ---
## Signif. codes:  0 '***' 0.001 '**' 0.01 '*' 0.05 '.' 0.1 ' ' 1
```

```
# Test whether doubling the sample size turns the treatment effect significan
t
d2double<-rbind(d2, d2)
m1double<-glmmTMB(selfing~treat + total.nr.dev.embryos.pre + nr.male.partners
            + nr.fem.partners + (1|P0mother), data=d2double, family=binomial)
summary(m1double) # still non-significant treatment effect
```

```
##  Family: binomial  ( logit )
## Formula:
## selfing ~ treat + total.nr.dev.embryos.pre + nr.male.partners +
##     nr.fem.partners + (1 | P0mother)
## Data: d2double
##
##      AIC      BIC   logLik deviance df.resid
##    136.1    152.3    -62.1    124.1      104
##
## Random effects:
##
## Conditional model:
##  Groups   Name        Variance Std.Dev.
##  P0mother (Intercept) 4.989    2.234
## Number of obs: 110, groups:  P0mother, 22
##
## Conditional model:
```

```
##                              Estimate Std. Error z value Pr(>|z|)
## (Intercept)                  0.088908   1.116608   0.080   0.9365
## treatrep                     2.256252   1.766752   1.277   0.2016
## total.nr.dev.embryos.pre     0.016256   0.006669   2.438   0.0148 *
## nr.male.partners            -1.272402   0.683536  -1.861   0.0627 .
## nr.fem.partners             -0.405180   0.432073  -0.938   0.3484
## ---
## Signif. codes:  0 '***' 0.001 '**' 0.01 '*' 0.05 '.' 0.1 ' ' 1
```

##### 6.3 Mortality rate prior to genotyping (for Methods)

```
d2$mort.rate.post<-((d2$total.nr.dev.embryos.post.reared-d2$total.surv.juv.po
st)/d2$total.nr.dev.embryos.post.reared)*100
mean(d2$mort.rate.post) # 53.36508

## [1] 53.36508

sd(d2$mort.rate.post) # 18.13457

## [1] 18.13457
```

#### 7. Figure 2

##### 7.1 Ascertaining selfing propensity for snails with estimated selfing rate

```
# Add a column specifying snails' propensity for selfing
d2$category<-ifelse(d2$selfing.rate == 0 & d2$total.nr.dev.embryos.pre == 0,
"Outcrosser", NA)
d2$category<-ifelse(d2$selfing.rate == 0 & d2$total.nr.dev.embryos.pre > 0, "
Plastic switcher", d2$category)
d2$category<-ifelse(d2$selfing.rate > 0 & d2$selfing.rate < 0.3 & d2$total.nr
.dev.embryos.pre > 0, "Plastic mixer", d2$category)
d2$category<-ifelse(d2$selfing.rate > 0.7, "Selfer", d2$category)
table(d2$selfing.rate, d2$category) # control - ok

##       
##         Outcrosser Plastic mixer Plastic switcher Selfer
##   0              5             0               26      0
##   0.06           0             2                0      0
##   0.07           0             1                0      0
##   0.08           0             2                0      0
##   0.13           0             2                0      0
##   0.2            0             2                0      0
##   0.83           0             0                0      1
##   1              0             0                0     14

table(d2$total.nr.dev.embryos.pre, d2$category) # control - ok

##      
##        Outcrosser Plastic mixer Plastic switcher Selfer
##   0             5             0                0      0
##   1             0             0                4      0
```

```
##   2            0             0                2      0
##   3            0             0                1      0
##   4            0             0                1      0
##   5            0             0                1      0
##   9            0             0                1      0
##   20           0             1                0      0
##   21           0             0                1      0
##   27           0             0                0      1
##   31           0             0                0      1
##   32           0             0                1      0
##   35           0             0                0      1
##   41           0             0                0      1
##   44           0             0                1      0
##   49           0             1                0      1
##   51           0             0                1      0
##   52           0             0                0      1
##   69           0             0                1      0
##   71           0             0                1      1
##   73           0             0                0      1
##   75           0             0                1      0
##   80           0             0                0      1
##   82           0             0                1      0
##   87           0             0                0      1
##   93           0             0                0      1
##   95           0             1                0      0
##   96           0             1                1      0
##   107          0             0                0      1
##   111          0             0                1      0
##   117          0             0                1      0
##   126          0             0                1      0
##   132          0             0                1      0
##   138          0             2                0      0
##   141          0             0                0      1
##   142          0             1                0      0
##   143          0             0                1      0
##   153          0             0                0      1
##   156          0             0                1      0
##   171          0             1                0      0
##   182          0             0                0      1
##   210          0             0                1      0
##   329          0             1                0      0
```

```
d2$category<-as.factor(d2$category)
d2$category<-factor(d2$category, levels=c("Selfer", "Plastic mixer",
                                          "Plastic switcher", "Outcrosser"))
# Frequency of selfing propensities
table(d2$category)/55
```

```
##
##           Selfer    Plastic mixer Plastic switcher       Outcrosser
##       0.27272727       0.16363636       0.47272727       0.09090909

table(d2$treat, d2$category)

##
##         Selfer Plastic mixer Plastic switcher Outcrosser
##   once     11             4               15          1
##   rep       4             5               11          4

# Chi-squared test for frequency of selfing propensities
# among once- and repeatedly paired snails
categories<-table(d2$category, d2$treat)
categories

##
##                    once rep
##   Selfer             11   4
##   Plastic mixer       4   5
##   Plastic switcher   15  11
##   Outcrosser          1   4

chisq.test(categories) # X-squared = 4.983, df = 3, p-value = 0.173

## Warning in chisq.test(categories): Chi-squared approximation may be incorr
ect

##
##  Pearson's Chi-squared test
##
## data:  categories
## X-squared = 4.983, df = 3, p-value = 0.173

chisq.test(categories)$expected

## Warning in chisq.test(categories): Chi-squared approximation may be incorr
ect

##
##                         once       rep
##   Selfer            8.454545  6.545455
##   Plastic mixer     5.072727  3.927273
##   Plastic switcher 14.654545 11.345455
##   Outcrosser        2.818182  2.181818

chisq.test(categories, simulate.p.value=TRUE) # X-squared = 4.983, df = NA, p
-value = 0.1959; still non-significant

##
##  Pearson's Chi-squared test with simulated p-value (based on 2000
##  replicates)
```

```
##
## data:  categories
## X-squared = 4.983, df = NA, p-value = 0.1934
```

```
fisher.test(categories) # p-value = 0.1986, i.e., also non-significant
```

```
##
##  Fisher's Exact Test for Count Data
##
## data:  categories
## p-value = 0.1986
## alternative hypothesis: two.sided
```

#### 7.2 Ascertaining selfing propensity for all female fertile snails

```
# Mark snails from the subsample and add their genotyping-based selfing prope
nsity to s
focals<-d2[, c("snailID", "category")]
s1<-merge(sna, focals, by="snailID", all=T)
nrow(s1) # 274 - ok
```

```
## [1] 274
```

```
s1$focal<-ifelse(is.na(s1$category)==F, "red", "lightskyblue4")
table(s1$focal, s1$category) # ok
```

```
##
##                  Selfer Plastic mixer Plastic switcher Outcrosser
##   lightskyblue4      0             0                0          0
##   red               15             9               26          5
```

```
# Restrict whole dataset to female fertile snails and exclude snails with mis
sing data and
# those only mated to a sibling
s2<-s1[s1$total.nr.dev.embryos>0,]
s2<-s2[s2$exclude %in% "no",]
nrow(s2) # 124
```

```
## [1] 124
```

```
# Specify the category of non-focal snails
s2$cat<-ifelse(s2$focal %in% "lightskyblue4" & s2$total.nr.fem.matings > 0 &
s2$total.nr.dev.embryos.pre ==0, "outc", NA)
s2$cat<-ifelse(s2$focal %in% "lightskyblue4" & s2$total.nr.fem.matings > 0 &
s2$total.nr.dev.embryos.pre > 0 & s2$total.nr.dev.embryos.post > 0, "plas", s
2$cat)
s2$cat<-ifelse(s2$focal %in% "lightskyblue4" & s2$total.nr.fem.matings == 0,
"self", s2$cat)
s2$cat<-ifelse(s2$focal %in% "lightskyblue4" & s2$total.nr.dev.embryos.pre >
0 & s2$total.nr.dev.embryos.post == 0, "self", s2$cat)

s2[s2$focal %in% "red", c("focal", "category", "cat")] # ok
```

```
##     focal          category  cat
## 1     red Plastic switcher <NA>
## 5     red Plastic switcher <NA>
## 6     red           Selfer <NA>
## 20    red    Plastic mixer <NA>
## 21    red    Plastic mixer <NA>
## 22    red Plastic switcher <NA>
## 23    red    Plastic mixer <NA>
## 45    red Plastic switcher <NA>
## 46    red           Selfer <NA>
## 53    red Plastic switcher <NA>
## 54    red Plastic switcher <NA>
## 57    red           Selfer <NA>
## 58    red Plastic switcher <NA>
## 61    red Plastic switcher <NA>
## 63    red    Plastic mixer <NA>
## 65    red Plastic switcher <NA>
## 69    red Plastic switcher <NA>
## 73    red           Selfer <NA>
## 83    red       Outcrosser <NA>
## 94    red Plastic switcher <NA>
## 100   red    Plastic mixer <NA>
## 101   red Plastic switcher <NA>
## 102   red       Outcrosser <NA>
## 103   red    Plastic mixer <NA>
## 106   red Plastic switcher <NA>
## 108   red       Outcrosser <NA>
## 109   red           Selfer <NA>
## 110   red Plastic switcher <NA>
## 115   red Plastic switcher <NA>
## 116   red Plastic switcher <NA>
## 122   red           Selfer <NA>
## 123   red           Selfer <NA>
## 126   red Plastic switcher <NA>
## 130   red           Selfer <NA>
## 132   red           Selfer <NA>
## 136   red Plastic switcher <NA>
## 144   red Plastic switcher <NA>
## 150   red    Plastic mixer <NA>
## 164   red           Selfer <NA>
## 165   red           Selfer <NA>
## 168   red Plastic switcher <NA>
## 181   red Plastic switcher <NA>
## 182   red    Plastic mixer <NA>
## 183   red Plastic switcher <NA>
## 188   red       Outcrosser <NA>
## 189   red           Selfer <NA>
## 190   red Plastic switcher <NA>
## 192   red    Plastic mixer <NA>
## 211   red       Outcrosser <NA>
```

```
## 220   red Plastic switcher <NA>
## 225   red Plastic switcher <NA>
## 226   red             Selfer <NA>
## 233   red             Selfer <NA>
## 269   red Plastic switcher <NA>
## 273   red             Selfer <NA>
```

```
s2[s2$focal %in% "lightskyblue4", c("focal", "category", "cat")] # ok
```

```
##             focal category  cat
## 2   lightskyblue4     <NA> outc
## 25  lightskyblue4     <NA> self
## 38  lightskyblue4     <NA> outc
## 40  lightskyblue4     <NA> outc
## 41  lightskyblue4     <NA> outc
## 48  lightskyblue4     <NA> self
## 52  lightskyblue4     <NA> outc
## 55  lightskyblue4     <NA> outc
## 56  lightskyblue4     <NA> self
## 59  lightskyblue4     <NA> outc
## 60  lightskyblue4     <NA> outc
## 64  lightskyblue4     <NA> self
## 67  lightskyblue4     <NA> plas
## 70  lightskyblue4     <NA> outc
## 75  lightskyblue4     <NA> outc
## 81  lightskyblue4     <NA> self
## 82  lightskyblue4     <NA> self
## 85  lightskyblue4     <NA> outc
## 98  lightskyblue4     <NA> outc
## 99  lightskyblue4     <NA> outc
## 114 lightskyblue4     <NA> self
## 117 lightskyblue4     <NA> self
## 124 lightskyblue4     <NA> outc
## 125 lightskyblue4     <NA> self
## 133 lightskyblue4     <NA> outc
## 135 lightskyblue4     <NA> self
## 139 lightskyblue4     <NA> outc
## 141 lightskyblue4     <NA> self
## 147 lightskyblue4     <NA> plas
## 149 lightskyblue4     <NA> self
## 152 lightskyblue4     <NA> plas
## 153 lightskyblue4     <NA> outc
## 157 lightskyblue4     <NA> self
## 158 lightskyblue4     <NA> self
## 159 lightskyblue4     <NA> self
## 163 lightskyblue4     <NA> outc
## 166 lightskyblue4     <NA> plas
## 169 lightskyblue4     <NA> outc
## 172 lightskyblue4     <NA> outc
## 177 lightskyblue4     <NA> outc
```

```
## 186 lightskyblue4     <NA> plas
## 191 lightskyblue4     <NA> self
## 193 lightskyblue4     <NA> plas
## 194 lightskyblue4     <NA> outc
## 195 lightskyblue4     <NA> plas
## 196 lightskyblue4     <NA> plas
## 198 lightskyblue4     <NA> outc
## 200 lightskyblue4     <NA> self
## 201 lightskyblue4     <NA> outc
## 202 lightskyblue4     <NA> outc
## 204 lightskyblue4     <NA> outc
## 206 lightskyblue4     <NA> outc
## 207 lightskyblue4     <NA> self
## 212 lightskyblue4     <NA> self
## 213 lightskyblue4     <NA> outc
## 214 lightskyblue4     <NA> self
## 215 lightskyblue4     <NA> plas
## 216 lightskyblue4     <NA> self
## 219 lightskyblue4     <NA> outc
## 221 lightskyblue4     <NA> outc
## 224 lightskyblue4     <NA> self
## 230 lightskyblue4     <NA> self
## 258 lightskyblue4     <NA> outc
## 261 lightskyblue4     <NA> outc
## 263 lightskyblue4     <NA> outc
## 264 lightskyblue4     <NA> plas
## 268 lightskyblue4     <NA> self
## 272 lightskyblue4     <NA> plas
## 274 lightskyblue4     <NA> self

table(s2$cat, s2$total.nr.fem.matings) # ok

## 
##         0  1  2  3  4  5  6  7  8 11
##   outc  0 14  8  5  1  0  2  1  1  1
##   plas  0  7  3  0  0  1  0  0  0  0
##   self 18  4  1  2  0  0  0  0  0  0

table(s2$cat, s2$total.nr.dev.embryos.pre) # ok

## 
##         0  1  2  3  4  5  7  8  9 10 11 18 20 21 23 27 31 32 34 35 41 42 4
4 49
##   outc 33  0  0  0  0  0  0  0  0  0  0  0  0  0  0  0  0  0  0  0  0  0
0  0
##   plas  0  2  0  0  0  0  0  0  0  0  0  1  0  0  1  0  0  0  1  1  0  1
0  0
##   self  7  6  1  0  2  1  1  1  0  1  1  0  0  1  0  0  0  0  0  0  0  0
0  0
## 
##        51 52 55 69 71 73 75 77 79 80 82 87 90 93 94 95 96 107 111 117 126
```

```
132
##   outc  0  0  0  0  0  0  0  0  0  0  0  0  0  0  0  0  0   0   0   0   0
0
##   plas  0  0  0  0  0  0  0  1  0  0  0  0  0  0  1  0  0   0   0   0   0
0
##   self  0  0  1  0  0  0  0  0  1  0  0  0  1  0  0  0  0   0   0   0   0
0
##
##        138 140 141 142 143 153 156 171 182 193 210 329
##   outc   0   0   0   0   0   0   0   0   0   0   0   0
##   plas   0   1   0   0   0   0   0   0   0   1   0   0
##   self   0   0   0   0   0   0   0   0   0   0   0   0
```

```
table(s2$cat, s2$total.nr.dev.embryos.post) # ok
```

```
##
##          0  2  4  5  6  7  8  9 10 11 12 13 14 16 18 26 30 31 33 34 36 51 5
3 58
##   outc  0  0  0  1  0  0  0  1  0  1  1  0  1  0  0  0  0  0  1  0  1  0
0  1
##   plas  0  0  1  0  1  1  1  0  2  0  0  0  0  0  1  0  0  1  0  0  0  0
0  0
##   self 12  1  0  1  0  0  1  0  0  0  0  1  1  1  1  1  0  0  0  1  0  0
0  0
##
##        59 64 65 68 70 73 77 82 88 89 90 95 98 102 105 107 141 149 158 167
169
##   outc  0  1  0  0  0  0  0  0  0  0  1  0  0   1   1   0   1   0   1   0
0
##   plas  0  0  0  0  0  1  0  1  0  0  0  0  0   0   0   0   0   1   0   0
0
##   self  1  0  0  0  1  0  0  0  0  0  0  0  0   0   0   1   0   0   0   0
0
##
##        170 173 178 181 189 191 192 193 194 199 201 211 216 225 232 236 249
253
##   outc   0   0   1   0   0   1   0   0   0   1   0   1   1   0   0   1   0
1
##   plas   0   0   0   0   0   0   0   0   0   0   0   0   0   0   0   0   0
0
##   self   0   0   0   0   1   0   0   0   0   0   0   0   0   0   0   0   0
0
##
##        265 269 271 274 279 280 307 311 316 320 335 337 365 366 370 372 375
381
##   outc   1   0   1   0   0   0   0   0   1   1   0   0   0   1   1   1   0
0
##   plas   0   0   0   0   0   0   0   0   0   0   0   0   0   0   0   0   0
0
##   self   0   0   0   0   0   0   0   0   0   0   0   0   0   0   0   0   0
```

```
0
##
##        387 390 391 397 420 421 438 439 446 448 457 462 467 478 482 545 571
641
##   outc   0   1   1   0   1   1   0   0   0   0   0   0   0   0   0   0   0
1
##   plas   0   0   0   0   0   0   0   0   0   0   0   0   0   0   0   0   0
0
##   self   0   0   0   0   0   0   0   0   0   0   0   0   0   0   0   0   0
0
##
##        698
##   outc   0
##   plas   0
##   self   0

# Combine the category assignments of focal and non-focal snails in one colum
n
# thereby merging the "Plastic switcher" and "Plastic mixer" groups of focals
s2$cat[s2$category %in% c("Plastic switcher", "Plastic mixer")]<-"plas"
s2$cat[s2$category %in% c("Selfer")]<-"self"
s2$cat[s2$category %in% c("Outcrosser")]<-"outc"

table(s2$category, s2$cat) # ok

##
##                      outc plas self
##   Selfer                0    0   15
##   Plastic mixer         0    9    0
##   Plastic switcher      0   26    0
##   Outcrosser            5    0    0

table(s2$cat) # ok

##
## outc plas self
##   38   46   40

s2$cat<-as.factor(s2$cat)
levels(s2$cat)

## [1] "outc" "plas" "self"

s2$cat <-factor(s2$cat, levels=c("self", "plas", "outc"))

# Frequency of selfing propensities
table(s2$cat)/124

##
##      self      plas      outc
## 0.3225806 0.3709677 0.3064516
```

```
# Chi-squared test for frequency of selfing propensities
# among once- and repeatedly paired snails
categories<-table(s2$cat, s2$group)
categories

##
##        paired.once paired.repeatedly
##   self          29                11
##   plas          26                20
##   outc          11                27

chisq.test(categories) # X-squared = 15.166, df = 2, p-value = 0.0005089

##
##  Pearson's Chi-squared test
##
## data:  categories
## X-squared = 15.166, df = 2, p-value = 0.0005089

chisq.test(categories)$expected

##
##        paired.once paired.repeatedly
##   self    21.29032          18.70968
##   plas    24.48387          21.51613
##   outc    20.22581          17.77419

chisq.test(categories, simulate.p.value=TRUE) # X-squared = 15.166, df = NA,
p-value = 0.0009995

##
##  Pearson's Chi-squared test with simulated p-value (based on 2000
##  replicates)
##
## data:  categories
## X-squared = 15.166, df = NA, p-value = 0.0009995

fisher.test(categories) # p-value = 0.0005025

##
##  Fisher's Exact Test for Count Data
##
## data:  categories
## p-value = 0.0005025
## alternative hypothesis: two.sided

# Frequency of selfing propensities within treatment groups
table(s2$cat[s2$group %in% "paired.once"])/66

##
##      self      plas      outc
## 0.4393939 0.3939394 0.1666667
```

```
table(s2$cat[s2$group %in% "paired.repeatedly"])/58

##
##      self      plas      outc
## 0.1896552 0.3448276 0.4655172
```

#### 7.3 Figure 2

```
# Frequencies for Fig. 2a

table(d2$selfing.rate, d2$treat)

##
##        once rep
##   0      16  15
##   0.06    2   0
##   0.07    0   1
##   0.08    1   1
##   0.13    0   2
##   0.2     1   1
##   0.83    0   1
##   1      11   3

# Frequencies for Fig. 2b

table(d2$total.nr.dev.embryos.pre[d2$selfing.rate %in% 0])

##
##   0   1   2   3   4   5   9  21  32  44  51  69  71  75  82  96 111 117 12
6 132
##   5   4   2   1   1   1   1   1   1   1   1   1   1   1   1   1   1   1
1   1
## 143 156 210
##   1   1   1

table(d2$total.nr.dev.embryos.pre[d2$selfing.rate > 0   & d2$selfing.rate < 0
.11])

##
##  20  49 138 171 329
##   1   1   1   1   1

table(d2$total.nr.dev.embryos.pre[d2$selfing.rate > 0.1 & d2$selfing.rate < 0
.21])

##
##  95  96 138 142
##   1   1   1   1

table(d2$total.nr.dev.embryos.pre[d2$selfing.rate > 0.2 & d2$selfing.rate < 0
.31])

## < table of extent 0 >
```

```
table(d2$total.nr.dev.embryos.pre[d2$selfing.rate > 0.3 & d2$selfing.rate < 0
.41])

## < table of extent 0 >

table(d2$total.nr.dev.embryos.pre[d2$selfing.rate > 0.4 & d2$selfing.rate < 0
.51])

## < table of extent 0 >

table(d2$total.nr.dev.embryos.pre[d2$selfing.rate > 0.5 & d2$selfing.rate < 0
.61])

## < table of extent 0 >

table(d2$total.nr.dev.embryos.pre[d2$selfing.rate > 0.6 & d2$selfing.rate < 0
.71])

## < table of extent 0 >

table(d2$total.nr.dev.embryos.pre[d2$selfing.rate > 0.7 & d2$selfing.rate < 0
.81])

## < table of extent 0 >

table(d2$total.nr.dev.embryos.pre[d2$selfing.rate > 0.8 & d2$selfing.rate < 0
.91])

##
## 73
##  1

table(d2$total.nr.dev.embryos.pre[d2$selfing.rate > 0.9])

##
##  27  31  35  41  49  52  71  80  87  93 107 141 153 182
##   1   1   1   1   1   1   1   1   1   1   1   1   1   1

###################

# Figure 2

#tiff(filename="Fig2.tif", width=170, height=85, units="mm",
#      res=600, pointsize=16)

par(mfrow=c(1,3))
par(mar=c(3.5,3.5,2,0), xpd=T)

# Fig. 2a

barplot(height=cbind("0" = c(16,15),
                    "0.01-0.1" = c(3,2),
```

```
                    "0.11-0.2" = c(1,3),
                    "0.21-0.3" = c(0,0),
                    "0.31-0.4" = c(0,0),
                    "0.41-0.5" = c(0,0),
                    "0.51-0.6" = c(0,0),
                    "0.61-0.7" = c(0,0),
                    "0.71-0.8" = c(0,0),
                    "0.81-0.9" = c(0,1),
                    "0.91-1.0" = c(11,3)),
beside=TRUE, col=c("lightskyblue1","green1"), xlab="",
ylab="", cex.axis=1, cex.lab=1, cex.names=1, las=1, xaxt="n", yaxt="n",
space=c(0, 0.6), ylim=c(0, 16))

title(ylab="# genotyped F1 snails", mgp=c(1.7,1,0), cex.lab=1)
title(xlab="Selfing rate post-isolation", mgp=c(2,1,0), cex.lab=1)
axis(1, tick=T, at=c(1.6, 14.5, 27.7), labels=c("0", "0.5", "1"), cex.axis=1,
las=1,
     mgp=c(2,0.5,0))
axis(2, tick=T, at=c(0,5,10,15), labels=c(0,5,10,15), cex.axis=1, las=1, mgp=
c(2,0.75,0))
text(-7.6, 16.5, "a", cex=1.5, font=2)

legend("top", legend=c("1 mating\npartner", "3-5 mating\npartners"),
fill=c("lightskyblue1","green1"), col=c("lightskyblue1","green1"), bty="n",
cex=1, x.intersp=0.75, y.intersp=1.75, inset=c(0.25,-0.05))

# Fig. 2b

barplot(height=cbind(       "0" = c(7,6,2,11,5),
                    "0.01-0.1" = c(3,0,1,1,0),
                    "0.11-0.2" = c(2,2,0,0,0),
                    "0.21-0.3" = c(0,0,0,0,0),
                    "0.31-0.4" = c(0,0,0,0,0),
                    "0.41-0.5" = c(0,0,0,0,0),
                    "0.51-0.6" = c(0,0,0,0,0),
                    "0.61-0.7" = c(0,0,0,0,0),
                    "0.71-0.8" = c(0,0,0,0,0),
                    "0.81-0.9" = c(0,1,0,0,0),
                    "0.91-1.0" = c(4,5,5,0,0)),
beside=FALSE, col=c("red", "lightskyblue4", "lightskyblue1", "yellow1", "gree
n1"), xlab="",
ylab="", cex.axis=1, cex.lab=1, cex.names=1, las=1, xaxt="n", yaxt="n")

title(ylab="# genotyped F1 snails", mgp=c(2,1,0), cex.lab=1)
title(xlab="Selfing rate post-isolation", mgp=c(2,1,0), cex.lab=1)
axis(1, tick=T, at=c(0.7, 6.7, 12.7), labels=c("0", "0.5", "1"), cex.axis=1,
las=1,
     mgp=c(2,0.5,0))
axis(2, tick=T, at=c(0,5,10,15,20,25,30), labels=c(0,5,10,15,20,25,30), cex.a
```

```
xis=1, las=1, mgp=c(2,0.75,0))

legend("top", title="# dev. embryos\nin isolation",
       legend=c("0", "1-25", "26-50","51-100", ">100"),
fill=c("green1", "yellow1", "lightskyblue1", "lightskyblue4","red"),
col=c("green1", "yellow1", "lightskyblue1", "lightskyblue4","red"), bty="n",
cex=1, inset=c(0, 0.07))

text(-4, 32, "b", cex=1.5, font=2)

# Fig. 2c

barplot(height=cbind(      "0" = c(0,0,26,5),
                    "0.01-0.1" = c(0,5,0,0),
                    "0.11-0.2" = c(0,4,0,0),
                    "0.21-0.3" = c(0,0,0,0),
                    "0.31-0.4" = c(0,0,0,0),
                    "0.41-0.5" = c(0,0,0,0),
                    "0.51-0.6" = c(0,0,0,0),
                    "0.61-0.7" = c(0,0,0,0),
                    "0.71-0.8" = c(0,0,0,0),
                    "0.81-0.9" = c(1,0,0,0),
                    "0.91-1.0" = c(14,0,0,0)),
beside=FALSE, col=c("black", "dimgray", "lightgrey", "white"), xlab="",
ylab="", cex.axis=1, cex.lab=1, cex.names=1, las=1, xaxt="n", yaxt="n")

title(ylab="# genotyped F1 snails", mgp=c(2,1,0), cex.lab=1)
title(xlab="Selfing rate post-isolation", mgp=c(2,1,0), cex.lab=1)
axis(1, tick=T, at=c(0.7, 6.7, 12.7), labels=c("0", "0.5", "1"), cex.axis=1,
las=1,
     mgp=c(2,0.5,0))
axis(2, tick=T, at=c(0,5,10,15,20,25,30), labels=c(0,5,10,15,20,25,30), cex.a
xis=1, las=1, mgp=c(2,0.75,0))

legend("top", legend=c("Outcrosser", "Plastic switcher", "Plastic mixer","Sel
fer"),
fill=c("white", "lightgrey", "dimgray","black"),
col=c("white", "lightgrey", "dimgray","black"), bty="n",
cex=1, inset=c(0.02, -0.03))

text(-4, 32, "c", cex=1.5, font=2)
```

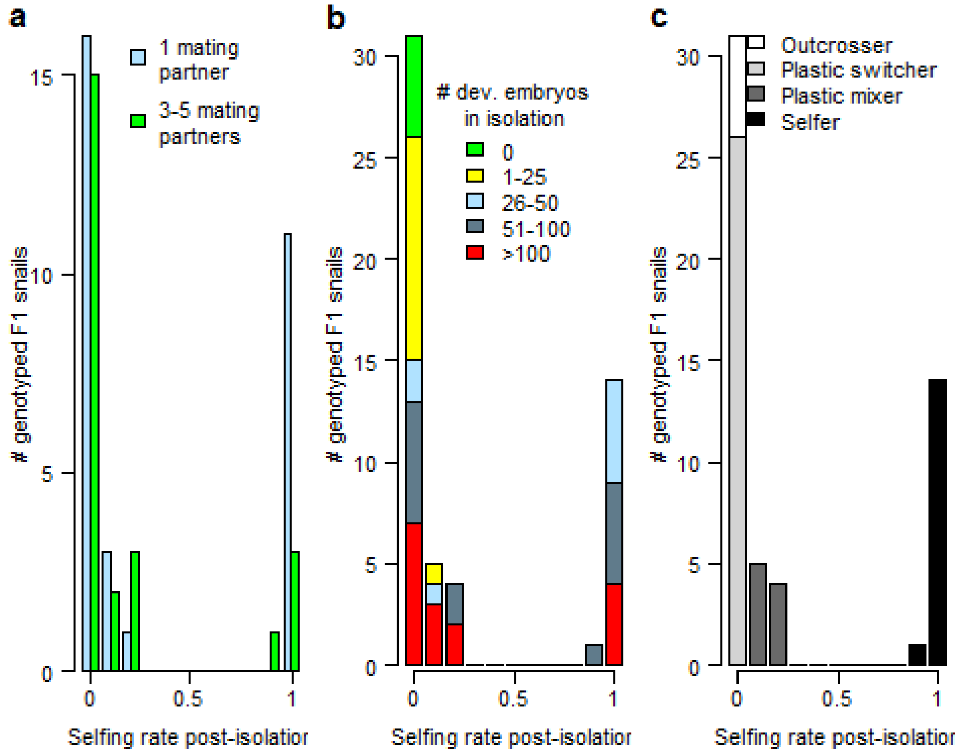

```
#dev.off()
```

#### 7.4 Figure S1

```
# Create a dataset including all snails except those excluded because of miss
ing data
# or because they were only paired with a sibling

s3<-sna[sna$exclude %in% "no",]
nrow(s3) # 268 ok

## [1] 268

# Compute the number of undeveloped embryos produced in isolation in s3 (268
snails)
# and in d2 (55 snails with selfing rate estimates)

s3$total.nr.undev.embryos.pre<-s3$total.nr.eggs.pre-s3$total.nr.dev.embryos.p
re
d2$total.nr.undev.embryos.pre<-d2$total.nr.eggs.pre-d2$total.nr.dev.embryos.p
re

# Frequencies for Figure S1b (undeveloped embryos produced in isolation)

table(d2$total.nr.undev.embryos.pre[d2$selfing.rate %in% 0])
```

```
##
##   0   1   2   7  11  13  15  17  22  27  40  60  64  68  72 128 143 154
##   6   2   6   1   2   2   1   1   1   1   1   1   1   1   1   1   1   1

table(d2$total.nr.undev.embryos.pre[d2$selfing.rate > 0   & d2$selfing.rate <
0.11])

##
##  11  12  23  66 115
##   1   1   1   1   1

table(d2$total.nr.undev.embryos.pre[d2$selfing.rate > 0.1 & d2$selfing.rate <
0.21])

##
## 34 49 62 70
##  1  1  1  1

table(d2$total.nr.undev.embryos.pre[d2$selfing.rate > 0.8 & d2$selfing.rate <
0.91])

##
## 26
##  1

table(d2$total.nr.undev.embryos.pre[d2$selfing.rate > 0.9])

##
##  1  3  4 10 22 32 33 39 40 45 61 97
##  1  2  1  1  1  1  1  2  1  1  1  1

# Frequencies for Figure S1c (all eggs produced in isolation)

table(d2$total.nr.eggs.pre[d2$selfing.rate %in% 0])

##
##   0   1   2   3   4  11  13  16  18  32  46  58  59  63  91 109 134 135 14
1 145
##   3   2   1   3   1   1   1   1   1   1   1   1   1   1   1   1   1   1
1   1
## 157 183 210 224 229 353
##   1   1   1   1   1   1

table(d2$total.nr.eggs.pre[d2$selfing.rate > 0   & d2$selfing.rate < 0.11])

##
##  32  72 237 253 340
##   1   1   1   1   1

table(d2$total.nr.eggs.pre[d2$selfing.rate > 0.1 & d2$selfing.rate < 0.21])
```

```
##
## 130 157 191 208
##   1   1   1   1
```

```
table(d2$total.nr.eggs.pre[d2$selfing.rate > 0.8 & d2$selfing.rate < 0.91])
```

```
##
## 99
##  1
```

```
table(d2$total.nr.eggs.pre[d2$selfing.rate > 0.9])
```

```
##
##  38  53  59  63  70  94  97 104 120 144 148 163 204 221
##   1   1   1   1   1   1   1   1   1   1   1   1   1   1
```

```
###################

# Figure S1

#tiff(filename="FigS1.tif", width=170, height=85, units="mm",
#     res=600, pointsize=16)

par(mfrow=c(1,3))
par(mar=c(3.5,3.1,2,0.6), xpd=T)

# Fig. S1a

barplot(height=cbind(      "0" = c(3,4,2,16,6),
                    "0.01-0.1" = c(1,1,0,3,0),
                    "0.11-0.2" = c(0,2,2,0,0),
                    "0.21-0.3" = c(0,0,0,0,0),
                    "0.31-0.4" = c(0,0,0,0,0),
                    "0.41-0.5" = c(0,0,0,0,0),
                    "0.51-0.6" = c(0,0,0,0,0),
                    "0.61-0.7" = c(0,0,0,0,0),
                    "0.71-0.8" = c(0,0,0,0,0),
                    "0.81-0.9" = c(0,0,1,0,0),
                    "0.91-1.0" = c(0,2,6,6,0)),
beside=FALSE, col=c("red", "lightskyblue4", "lightskyblue1", "yellow1", "gree
n1"), xlab="",
ylab="", cex.axis=1, cex.lab=1, cex.names=1, las=1, xaxt="n", yaxt="n")

title(ylab="# genotyped F1 snails", mgp=c(2,1,0), cex.lab=1)
title(xlab="Selfing rate post-isolation", mgp=c(2,1,0), cex.lab=1)
axis(1, tick=T, at=c(0.7, 6.7, 12.7), labels=c("0", "0.5", "1"), cex.axis=1,
las=1,
     mgp=c(2,0.5,0))
axis(2, tick=T, at=c(0,5,10,15,20,25,30), labels=c(0,5,10,15,20,25,30), cex.a
xis=1, las=1, mgp=c(2,0.75,0))
```

```
legend("top", title="# undev. embryos\nin isolation",
       legend=c("0", "1-25", "26-50","51-100", ">100"),
fill=c("green1", "yellow1", "lightskyblue1", "lightskyblue4","red"),
col=c("green1", "yellow1", "lightskyblue1", "lightskyblue4","red"), bty="n",
cex=1, inset=c(0, 0.07))

text(-4, 32, "a", cex=1.5, font=2)

# Fig. S1b

barplot(height=cbind(       "0" = c(11,4,2,11,3),
                    "0.01-0.1" = c(3,1,1,0,0),
                    "0.11-0.2" = c(4,0,0,0,0),
                    "0.21-0.3" = c(0,0,0,0,0),
                    "0.31-0.4" = c(0,0,0,0,0),
                    "0.41-0.5" = c(0,0,0,0,0),
                    "0.51-0.6" = c(0,0,0,0,0),
                    "0.61-0.7" = c(0,0,0,0,0),
                    "0.71-0.8" = c(0,0,0,0,0),
                    "0.81-0.9" = c(0,1,0,0,0),
                    "0.91-1.0" = c(7,6,1,0,0)),
beside=FALSE, col=c("red", "lightskyblue4", "lightskyblue1", "yellow1", "gree
n1"), xlab="",
ylab="", cex.axis=1, cex.lab=1, cex.names=1, las=1, xaxt="n", yaxt="n")

title(ylab="# genotyped F1 snails", mgp=c(2,1,0), cex.lab=1)
title(xlab="Selfing rate post-isolation", mgp=c(2,1,0), cex.lab=1)
axis(1, tick=T, at=c(0.7, 6.7, 12.7), labels=c("0", "0.5", "1"), cex.axis=1,
las=1,
     mgp=c(2,0.5,0))
axis(2, tick=T, at=c(0,5,10,15,20,25,30), labels=c(0,5,10,15,20,25,30), cex.a
xis=1, las=1, mgp=c(2,0.75,0))

legend("top", title="# eggs\nin isolation",
       legend=c("0", "1-25", "26-50","51-100", ">100"),
fill=c("green1", "yellow1", "lightskyblue1", "lightskyblue4","red"),
col=c("green1", "yellow1", "lightskyblue1", "lightskyblue4","red"), bty="n",
cex=1, inset=c(0, 0.07))

text(-4, 32, "b", cex=1.5, font=2)

# Fig. S1c

plot(s3$total.nr.undev.embryos.pre~s3$total.nr.dev.embryos.pre, col="white",
     las=1, xlab="", ylab="", xaxt="n", yaxt="n", ylim=c(0, 200))

title(ylab="# undev. embryos in isolation", mgp=c(2.2,1,0), cex.lab=1)
title(xlab="# dev. embryos in isolation", mgp=c(2,1,0), cex.lab=1)
```

```
axis(1, tick=T, at=c(0, 100, 200, 300), labels=c("0", "100", "200", "300"),
     cex.axis=1, las=1, mgp=c(2,0.5,0))
axis(2, tick=T, at=c(0, 50, 100, 150, 200),
     labels=c("0", "50", "100", "150", "200"), cex.axis=1,
     las=1, mgp=c(2,0.75,0))

points(s3$total.nr.undev.embryos.pre[s3$nr.assigned>0]~s3$total.nr.dev.embryo
s.pre[s3$nr.assigned>0], col="red")
points(s3$total.nr.undev.embryos.pre[s3$nr.assigned==0]~s3$total.nr.dev.embry
os.pre[s3$nr.assigned==0], col="lightskyblue4")

legend("topright", title="Offspring\ngenotyped", legend=c("Yes", "No"),
       col=c("red", "lightskyblue4"), pch=c(1,1), bty="n",
       cex=1, inset=c(0, 0.07))

text(-115, 213, "c", cex=1.5, font=2)
```

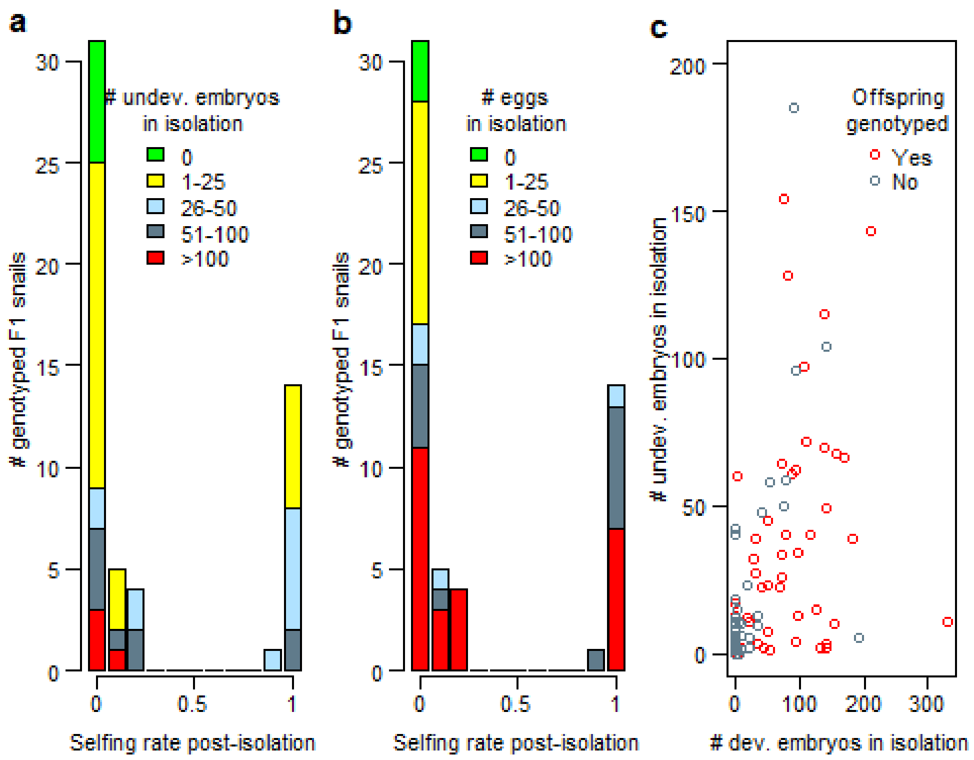

```
#dev.off()
```

#### 7.5 Relationship between number of developed vs undeveloped embryos in isolation

```
# How often did snails without developed embryos also lack undeveloped embryo
s?

nrow(s3[s3$total.nr.dev.embryos.pre==0,]) # 189
```

```
## [1] 189

nrow(s3[s3$total.nr.dev.embryos.pre==0 & s3$total.nr.undev.embryos.pre==0,])
# 164

## [1] 164

164/189 # 0.8677249 proportion of snails lacking both developed and undevelop
ed embryos

## [1] 0.8677249

# How many undeveloped embryos did the remaining 13.2% of snails produce?

nrow(s3[s3$total.nr.dev.embryos.pre==0 & s3$total.nr.undev.embryos.pre>0,]) #
25 snails

## [1] 25

mean(s3$total.nr.undev.embryos.pre[s3$total.nr.dev.embryos.pre==0 & s3$total.
nr.undev.embryos.pre>0])

## [1] 8.6

# 8.6
sd  (s3$total.nr.undev.embryos.pre[s3$total.nr.dev.embryos.pre==0 & s3$total.
nr.undev.embryos.pre>0])

## [1] 10.87811

# 10.87811
max (s3$total.nr.undev.embryos.pre[s3$total.nr.dev.embryos.pre==0 & s3$total.
nr.undev.embryos.pre>0])

## [1] 42

# 42

# What about snails with max. 10 developed embryos?

nrow(s3[s3$total.nr.dev.embryos.pre %in% c(1:10) & s3$total.nr.undev.embryos.
pre>0,]) # 17 snails

## [1] 17

mean(s3$total.nr.undev.embryos.pre[s3$total.nr.dev.embryos.pre %in% c(1:10) &
s3$total.nr.undev.embryos.pre>0])

## [1] 8.411765

# 8.411765
sd  (s3$total.nr.undev.embryos.pre[s3$total.nr.dev.embryos.pre %in% c(1:10) &
s3$total.nr.undev.embryos.pre>0])
```

```
## [1] 14.24368

# 14.24368
max (s3$total.nr.undev.embryos.pre[s3$total.nr.dev.embryos.pre %in% c(1:10) &
s3$total.nr.undev.embryos.pre>0])

## [1] 60

# 60

# Correlation between number of developed and undeveloped embryos in isolatio
n

cor.test(s3$total.nr.dev.embryos.pre, s3$total.nr.undev.embryos.pre)

##
##  Pearson's product-moment correlation
##
## data:  s3$total.nr.dev.embryos.pre and s3$total.nr.undev.embryos.pre
## t = 12.531, df = 266, p-value < 2.2e-16
## alternative hypothesis: true correlation is not equal to 0
## 95 percent confidence interval:
##  0.5279780 0.6794742
## sample estimates:
##       cor
## 0.6092559

# r = 0.6092559, t = 12.531, df = 266, p-value < 2.2e-16
```

#### 7.6 Figure S2

```
#tiff(filename="FigS2.tif", width=85, height=85, units="mm",
#     res=600, pointsize=11)

par(mfrow=c(1,1))
par(mar=c(3.5,3.5,1,0), xpd=T)

plot(s2$total.nr.dev.embryos.post~s2$total.nr.dev.embryos.pre, col="white",
     las=1, xlab="", ylab="", xaxt="n", yaxt="n", xlim=c(-10, 330))

title(ylab="# dev. embryos post-isol.", mgp=c(2.2,1,0), cex.lab=1)
title(xlab="# dev. embryos in isolation", mgp=c(2,1,0), cex.lab=1)
axis(1, tick=T, at=c(0, 100, 200, 300), labels=c("0", "100", "200", "300"),
     cex.axis=1, las=1, mgp=c(2,0.5,0))
axis(2, tick=T, at=c(0, 100, 200, 300, 400, 500, 600, 700),
     labels=c("0", "100", "200", "300", "400", "500", "600", "700"), cex.axis
=1,
     las=1, mgp=c(2,0.5,0))
segments(x0=0, y0=0, x1=0, y1=700, lwd=8, col="lightgrey")
segments(x0=0, y0=0, x1=330, y1=0, lwd=8, col="lightgrey")
```

```
points(s2$total.nr.dev.embryos.post~s2$total.nr.dev.embryos.pre, col=s2$focal
)

legend("topright", title="Offspring\ngenotyped", legend=c("Yes", "No"),
       col=c("red", "lightskyblue4"), pch=c(1,1), bty="n",
       cex=1, inset=c(0, 0.07))
```

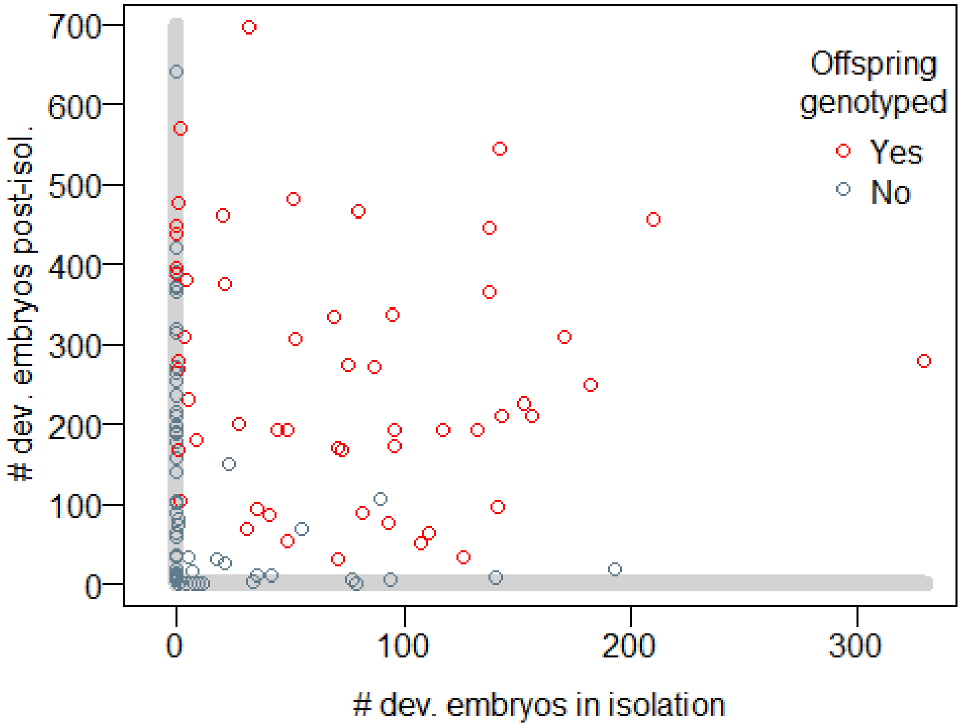

```
#dev.off()
```

#### 8. Figure 3

##### 8.1 Preparations

```
# Compute proportion of undeveloped embryos (for Figure 3)
s2$total.nr.undev.embryos.pre<-s2$total.nr.eggs.pre-s2$total.nr.dev.embryos.p
re
s2$total.nr.undev.embryos.post<-s2$total.nr.eggs.post-s2$total.nr.dev.embryos
.post
s2$total.nr.undev.embryos<-s2$total.nr.undev.embryos.pre+s2$total.nr.undev.em
bryos.post
s2$prop.undev.embryos<-(s2$total.nr.undev.embryos/s2$total.nr.eggs)*100

# Compute proportion of undeveloped embryos (for Figure S3)
d2$total.nr.undev.embryos.post<-d2$total.nr.eggs.post-d2$total.nr.dev.embryos
.post
d2$total.nr.undev.embryos<-d2$total.nr.undev.embryos.pre+d2$total.nr.undev.em
```

```
bryos.post
d2$prop.undev.embryos<-(d2$total.nr.undev.embryos/d2$total.nr.eggs)*100

# Compute proportion of non-surviving juveniles (for Figure 3)
colnames(sur)[1]<-"snailID"
s2<-merge(s2, sur, by="snailID", all.x=T)
nrow(s2) # 124 ok
```

```
## [1] 124
```

```
s2$nr.premating.snails<-ifelse(is.na(s2$nr.premating.snails)==T, 0, s2$nr.pre
mating.snails)
s2$nr.postmating.snails<-ifelse(is.na(s2$nr.postmating.snails)==T, 0, s2$nr.p
ostmating.snails)
s2$total.surv.juv<-s2$nr.premating.snails+s2$nr.postmating.snails
s2$total.nr.dev.embryos.reared<-s2$total.nr.dev.embryos.pre.reared+s2$total.n
r.dev.embryos.post.reared
s2$mort.rate<-((s2$total.nr.dev.embryos.reared-s2$total.surv.juv)/
  s2$total.nr.dev.embryos.reared)*100
s2[,c("snailID", "total.nr.dev.embryos.reared", "total.surv.juv", "mort.rate"
)]
```

```
##     snailID total.nr.dev.embryos.reared total.surv.juv  mort.rate
## 1    11_4.1                         330             37  88.787879
## 2    14_3.1                         309            131  57.605178
## 3    14_3.7                         475            188  60.421053
## 4   17_1.10                         356            163  54.213483
## 5    22_2.3                         448            133  70.312500
## 6    22_2.6                         438            116  73.515982
## 7    22_4.1                          93             51  45.161290
## 8    22_5.2                         444            150  66.216216
## 9   25_1.10                           0              0        NaN
## 10   25_4.1                         163             77  52.760736
## 11   25_7.5                         275            117  57.454545
## 12   25_7.9                         177            117  33.898305
## 13   31_2.1                         451            163  63.858093
## 14   31_2.2                         344             54  84.302326
## 15   31_2.4                         112             28  75.000000
## 16   31_3.1                         105             16  84.761905
## 17   31_3.3                         227            115  49.339207
## 18   31_3.4                         142             77  45.774648
## 19   31_3.5                         174             74  57.471264
## 20   31_3.6                          23             12  47.826087
## 21   31_3.7                         166             46  72.289157
## 22   31_4.2                         223             89  60.089686
## 23   31_4.5                         294            131  55.442177
## 24   31_4.6                         102             66  35.294118
## 25   31_4.8                         160             64  60.000000
## 26   34_2.6                         186            117  37.096774
## 27   34_2.7                           0              0        NaN
```

```
## 28   34_2.8                                   453            140  69.094923
## 29   36_2.7                                    83              1  98.795181
## 30   38_1.1                                   209            146  30.143541
## 31  38_10.2                                   237            100  57.805907
## 32   38_2.1                                    91             39  57.142857
## 33   38_3.2                                   260            124  52.307692
## 34   38_5.3                                     1              0 100.000000
## 35   38_5.6                                    11              6  45.454545
## 36   38_6.1                                   250            166  33.600000
## 37   38_9.7                                   156             40  74.358974
## 38  40_1.25                                   220            128  41.818182
## 39   40_1.4                                   472            111  76.483051
## 40   40_1.5                                   131             46  64.885496
## 41   40_1.6                                   335            141  57.910448
## 42   40_1.7                                   320            143  55.312500
## 43   40_1.8                                   371            102  72.506739
## 44   40_1.9                                   380            127  66.578947
## 45  40_3.10                                   361            192  46.814404
## 46  40_3.16                                   239            148  38.075314
## 47   40_3.7                                   114             46  59.649123
## 48   40_4.2                                   329            212  35.562310
## 49   43_5.3                                   157             31  80.254777
## 50   43_5.4                                   176             37  78.977273
## 51   43_5.5                                   206            124  39.805825
## 52  44_3b.1                                    21              2  90.476190
## 53   45_1.2                                   281             46  83.629893
## 54   45_1.3                                    99             22  77.777778
## 55   45_1.5                                    57             24  57.894737
## 56   45_1.8                                     1              1   0.000000
## 57   45_2.3                                   262            164  37.404580
## 58   45_9.1                                   324             55  83.024691
## 59   46_2.4                                   206             43  79.126214
## 60   46_2.5                                     0              0        NaN
## 61   48_7.1                                     0              0        NaN
## 62   51_3.2                                   168            124  26.190476
## 63  52_10.1                                   211             18  91.469194
## 64   52_3.1                                     0              0        NaN
## 65  52_4b.4                                   287             89  68.989547
## 66   52_8.4                                   172             21  87.790698
## 67   54_2.1                                    38             22  42.105263
## 68  54_3a.1                                   369            136  63.143631
## 69  54_3b.1                                    64             20  68.750000
## 70   54_5.2                                    58             28  51.724138
## 71   55_8.1                                    79              1  98.734177
## 72   58_1.1                                     0              0        NaN
## 73   58_1.3                                     1              1   0.000000
## 74   61_1.3                                    33              7  78.787879
## 75   61_3.1                                   151             28  81.456954
## 76   61_7.2                                   100             42  58.000000
## 77   61_7.4                                    49             11  77.551020
```

```
## 78   63_1.7                           231          121  47.619048
## 79   63_1.8                           173          129  25.433526
## 80  64_1.12                             5            2  60.000000
## 81   64_3.4                             9            9   0.000000
## 82   67_2.7                           274           46  83.211679
## 83   70_1.5                           221           69  68.778281
## 84   70_1.6                            87           30  65.517241
## 85  70_3.12                           101           17  83.168317
## 86  70_3.14                           268          152  43.283582
## 87  70_3.18                           128           44  65.625000
## 88  70_3.20                           192           91  52.604167
## 89  70_3.22                             0            0        NaN
## 90  70_3.26                           415           94  77.349398
## 91  70_3.28                            45           12  73.333333
## 92  70_3.29                            30           25  16.666667
## 93   70_3.3                            28            0 100.000000
## 94  70_3.31                            44           27  38.636364
## 95   70_3.4                           270           95  64.814815
## 96  70_5.13                             1            0 100.000000
## 97   70_5.2                           133           68  48.872180
## 98   70_5.5                            60            9  85.000000
## 99   70_5.8                           231           84  63.636364
## 100  70_8.2                             0            0        NaN
## 101  70_8.4                             1            1   0.000000
## 102  70_9.5                           285          154  45.964912
## 103  71_3.4                           110           38  65.454545
## 104  71_3.5                           211          124  41.232227
## 105  75_3.2                             1            0 100.000000
## 106  75_5.1                           211           63  70.142180
## 107  75_6.2                            12            6  50.000000
## 108  75_6.6                           349          116  66.762178
## 109  75_7.1                           265           91  65.660377
## 110  76_2.3                           234          107  54.273504
## 111  76_3.4                             2            0 100.000000
## 112  79_2.1                           168          104  38.095238
## 113  79_4.4                           100           44  56.000000
## 114  83_3.1                             4            1  75.000000
## 115  83_3.4                           416          229  44.951923
## 116   9_2.7                             0            0        NaN
## 117  9_3.12                            11           11   0.000000
## 118  9_3.15                            14           13   7.142857
## 119  9_3.16                            51           20  60.784314
## 120   9_3.5                             4            2  50.000000
## 121   9_4.2                           199          134  32.663317
## 122   9_4.7                           148           40  72.972973
## 123   9_5.1                           169           26  84.615385
## 124   9_5.4                             0            0        NaN
```

```
# Compute proportion of non-surviving juveniles (for Figure S3)
d2$total.surv.juv<-d2$total.surv.juv.pre+d2$total.surv.juv.post
```

```
d2$total.nr.dev.embryos.reared<-d2$total.nr.dev.embryos.pre.reared+d2$total.n
r.dev.embryos.post.reared
d2$mort.rate<-((d2$total.nr.dev.embryos.reared-d2$total.surv.juv)/
  d2$total.nr.dev.embryos.reared)*100

# new column with colours for data points (black for focals, colours for non-
focals)
s2$focal2<-ifelse(s2$cat %in% "self", "red", NA)
s2$focal2<-ifelse(s2$cat %in% "plas", "lightskyblue3", s2$focal2)
s2$focal2<-ifelse(s2$cat %in% "outc", "green1", s2$focal2)
s2$focal2<-ifelse(s2$focal %in% "red", "black", s2$focal2)
table(s2$focal2, s2$focal) # ok

##
##                  lightskyblue4 red
##   black                      0  55
##   green1                    33   0
##   lightskyblue3             11   0
##   red                       25   0

table(s2$focal2, s2$cat) # ok

##
##                  self plas outc
##   black            15   35    5
##   green1            0    0   33
##   lightskyblue3     0   11    0
##   red              25    0    0
```

#### 8.2 Figure 3

```
#tiff(filename="Fig3.tif", width=190, height=85, units="mm",
#     res=600, pointsize=16)

par(mfrow=c(1,3))

# Fig. 3a

par(mar=c(4.5,3,0,0), xpd=T)

plot(s2$total.nr.dev.embryos~s2$cat, las=1, xlab="", xaxt="n", yaxt="n",
     ylab="", col=c("red", "lightskyblue3", "green1"),
     outlier.color=NA, cex.lab=1, cex.axis=1, ylim=c(0, 750))
title(ylab="Female LRS [# dev. embryos]", mgp=c(2.2, 1, 0), cex.lab=1)
title(xlab="Propensity for selfing", mgp=c(3.5, 1, 0), cex.lab=1)
axis(1, tick=F, at=c(1, 2, 3), labels=c("App.\nselfer\n",
                                        "App.\nplastic\nsnail",
                                        "App.\nout-\ncrosser"),
     cex.axis=1, las=1, mgp=c(2, 2, 0))

axis(2, tick=T, at=c(0,200,400,600), labels=c(0,200,400,600), cex.axis=1, las
```

```
=1, mgp=c(2,0.75,0))
beeswarm(s2$total.nr.dev.embryos~s2$cat, pch=21,
         pwbg=s2$focal2, cex=1, spacing=0.7, add=T, ylim=c(0, 750))
points(tapply(s2$total.nr.dev.embryos, s2$cat, mean, na.rm=T), pch=24, bg="wh
ite", cex=1.5)
text(0.55, 725, "a", cex=1.5, font=2)

# Fig. 3b

plot(s2$prop.undev.embryos~s2$cat, las=1, xlab="", xaxt="n", yaxt="n",
     ylab="", col=c("red", "lightskyblue3", "green1"),
     outlier.color=NA, cex.lab=1, cex.axis=1, ylim=c(0, 100))
title(ylab="% undeveloped embryos", mgp=c(1.9, 1, 0), cex.lab=1)
title(xlab="Propensity for selfing", mgp=c(3.5, 1, 0), cex.lab=1)
axis(1, tick=F, at=c(1, 2, 3), labels=c("App.\nselfer\n",
                                        "App.\nplastic\nsnail",
                                        "App.\nout-\ncrosser"),
     cex.axis=1, las=1, mgp=c(2, 2, 0))

axis(2, tick=T, at=c(0,20,40,60,80,100), labels=c(0,20,40,60,80,100), cex.axi
s=1, las=1, mgp=c(2,0.75,0))
beeswarm(s2$prop.undev.embryos~s2$cat, pch=21,
         pwbg=s2$focal2, cex=1, spacing=0.8, add=T, ylim=c(0, 100))
points(tapply(s2$prop.undev.embryos, s2$cat, mean, na.rm=T), pch=24, bg="whit
e", cex=1.5)
text(0.55, 97, "b", cex=1.5, font=2)

# Fig. 3c

plot(s2$mort.rate~s2$cat, las=1, xlab="", xaxt="n", yaxt="n",
     ylab="", col=c("red", "lightskyblue3", "green1"),
     outlier.color=NA, cex.lab=1, cex.axis=1, ylim=c(0, 100))
title(ylab="% dead juveniles", mgp=c(1.9, 1, 0), cex.lab=1)
title(xlab="Propensity for selfing", mgp=c(3.5, 1, 0), cex.lab=1)
axis(1, tick=F, at=c(1, 2, 3), labels=c("App.\nselfer\n",
                                        "App.\nplastic\nsnail",
                                        "App.\nout-\ncrosser"),
     cex.axis=1, las=1, mgp=c(2, 2, 0))

axis(2, tick=T, at=c(0,20,40,60,80,100), labels=c(0,20,40,60,80,100), cex.axi
s=1, las=1, mgp=c(2,0.75,0))
beeswarm(s2$mort.rate~s2$cat, pch=21,
         pwbg=s2$focal2, cex=1, spacing=0.8, add=T, ylim=c(0, 100))
points(tapply(s2$mort.rate, s2$cat, mean, na.rm=T), pch=24, bg="white", cex=1
.5)
text(0.55, 97, "c", cex=1.5, font=2)
```

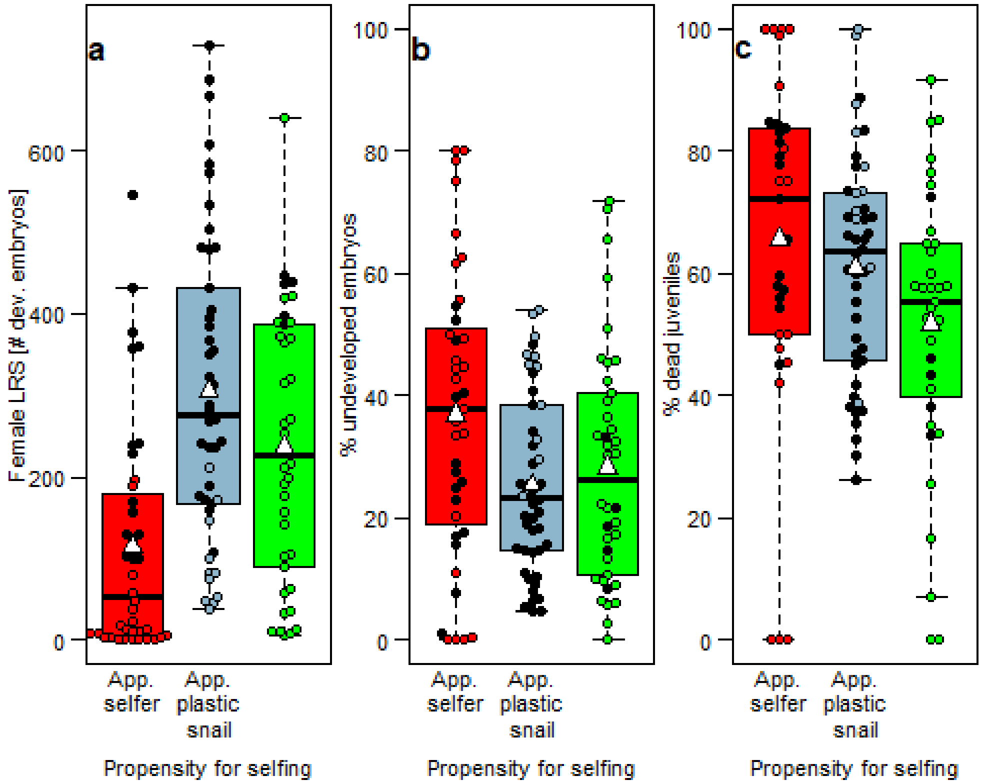

```
#dev.off()
```

#### 8.3 Figure S3

```
#tiff(filename="FigS3.tif", width=170, height=170, units="mm",
#      res=600, pointsize=13)

par(mfrow=c(2,2))

# Fig. S3a

par(mar=c(4.5,3,1,0), xpd=T)

plot(d2$total.nr.dev.embryos~d2$category, las=1, xlab="", xaxt="n", yaxt="n",
     ylab="", col=c("red", "lightskyblue4", "lightskyblue1", "green1"),
     outlier.color=NA, cex.lab=1, cex.axis=1, ylim=c(0, 750))
title(ylab="Female LRS [# dev. embryos]", mgp=c(2.2,1,0), cex.lab=1)
title(xlab="Propensity for selfing", mgp=c(3.5, 1, 0), cex.lab=1)
axis(1, tick=F, at=c(1, 2, 3, 4), labels=c("Selfer\n\n",
                                          "Plastic\nmixer\n",
                                          "Plastic\nswit-\ncher",
                                          "Out-\ncros-\nser"),
     cex.axis=1, las=1, mgp=c(2,2,0))
axis(2, tick=T, at=c(0,200,400,600), labels=c(0,200,400,600), cex.axis=1, las
=1, mgp=c(2,0.75,0))
beeswarm(d2$total.nr.dev.embryos~d2$category, pch=21,
         bg=c("red", "lightskyblue4", "lightskyblue1", "green1"), cex=1, spac
```

```
ing=0.8, add=T, ylim=c(0, 750))
points(tapply(d2$total.nr.dev.embryos, d2$category, mean, na.rm=T), pch=24, b
g="white", cex=1.5)
text(0.55, 725, "a", cex=1.5, font=2)

# Fig. S3b

plot(d2$prop.undev.embryos~d2$category, las=1, xlab="", xaxt="n", yaxt="n",
     ylab="", col=c("red", "lightskyblue4", "lightskyblue1", "green1"),
     outlier.color=NA, cex.lab=1, cex.axis=1, ylim=c(0, 100))
title(ylab="% undeveloped embryos", mgp=c(1.9, 1, 0), cex.lab=1)
title(xlab="Propensity for selfing", mgp=c(3.5, 1, 0), cex.lab=1)
axis(1, tick=F, at=c(1, 2, 3, 4), labels=c("Selfer\n\n",
                                        "Plastic\nmixer\n",
                                        "Plastic\nswit-\ncher",
                                        "Out-\ncros-\nser"),
     cex.axis=1, las=1, mgp=c(2,2,0))
axis(2, tick=T, at=c(0,20,40,60,80,100), labels=c(0,20,40,60,80,100), cex.axi
s=1, las=1, mgp=c(2,0.75,0))
beeswarm(d2$prop.undev.embryos~d2$category, pch=21,
         bg=c("red", "lightskyblue4", "lightskyblue1", "green1"), cex=1, spac
ing=0.8, add=T, ylim=c(0, 100))
points(tapply(d2$prop.undev.embryos, d2$category, mean, na.rm=T), pch=24, bg=
"white", cex=1.5)
text(0.55, 97, "b", cex=1.5, font=2)

# Fig. S3c

plot(d2$mort.rate~d2$category, las=1, xlab="", xaxt="n", yaxt="n",
     ylab="", col=c("red", "lightskyblue4", "lightskyblue1", "green1"),
     outlier.color=NA, cex.lab=1, cex.axis=1, ylim=c(0, 100))
title(ylab="% dead juveniles", mgp=c(1.9, 1, 0), cex.lab=1)
title(xlab="Propensity for selfing", mgp=c(3.5, 1, 0), cex.lab=1)
axis(1, tick=F, at=c(1, 2, 3, 4), labels=c("Selfer\n\n",
                                        "Plastic\nmixer\n",
                                        "Plastic\nswit-\ncher",
                                        "Out-\ncros-\nser"),
     cex.axis=1, las=1, mgp=c(2,2,0))

axis(2, tick=T, at=c(0,20,40,60,80,100), labels=c(0,20,40,60,80,100), cex.axi
s=1, las=1, mgp=c(2,0.75,0))
beeswarm(d2$mort.rate~d2$category, pch=21,
         bg=c("red", "lightskyblue4", "lightskyblue1", "green1"), cex=1, spac
ing=0.8, add=T, ylim=c(0, 100))
points(tapply(d2$mort.rate, d2$category, mean, na.rm=T), pch=24, bg="white",
cex=1.5)
text(0.55, 97, "c", cex=1.5, font=2)

#dev.off()
```

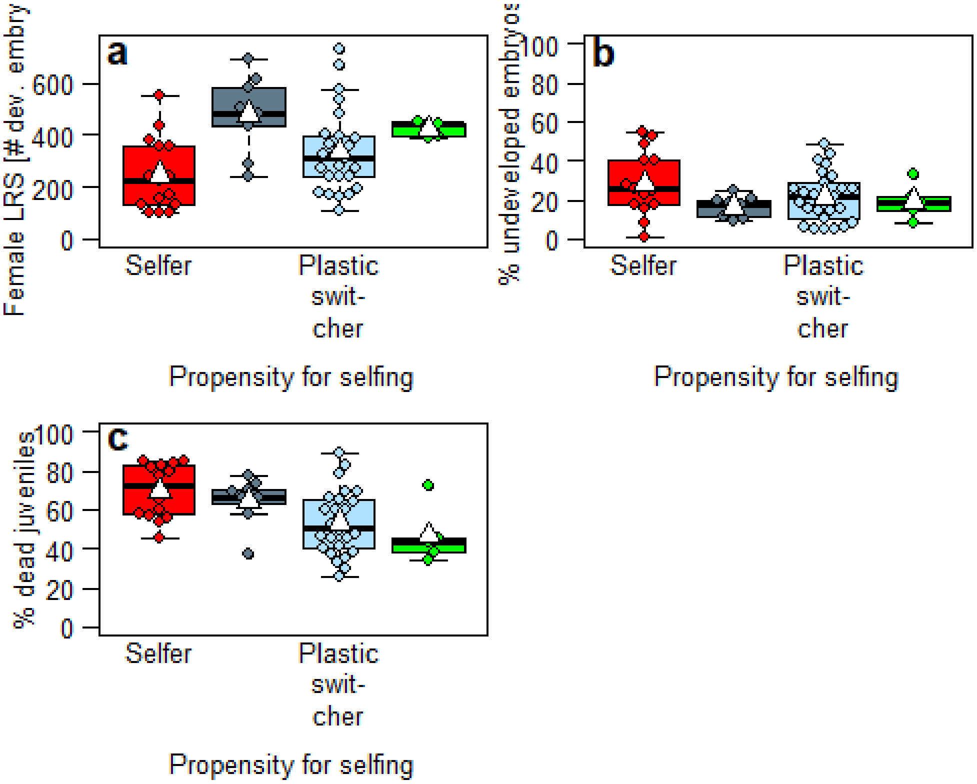

#### 9. Models 2-7: Do snails with different selfing propensity differ in female LRS?

##### 9.1 Preparations

```
# exclude outcrossers because of low sample size (n = 5)
d3<-d2[d2$category %in% c("Selfer", "Plastic mixer", "Plastic switcher"),]
nrow(d3) # 50 ok

## [1] 50
```

##### 9.2 Model 2: Number of developed embryos in snails with selfing rate estimate (Table S3)

```
m2<-glmmTMB(total.nr.dev.embryos ~ category + shell.size + (1|P0mother) + (1|
pairID), data=d3, family=nbinom1)
summary(m2)

##  Family: nbinom1  ( log )
## Formula:
## total.nr.dev.embryos ~ category + shell.size + (1 | P0mother) +
##     (1 | pairID)
## Data: d3
##
##      AIC      BIC   logLik deviance df.resid
##    640.2    653.6   -313.1    626.2       43
```

```
##
## Random effects:
##
## Conditional model:
##  Groups   Name        Variance  Std.Dev.
##  P0mother (Intercept) 2.721e-11 5.216e-06
##  pairID   (Intercept) 2.450e-09 4.950e-05
## Number of obs: 50, groups:  P0mother, 22; pairID, 42
##
## Dispersion parameter for nbinom1 family (): 56.1
##
## Conditional model:
##                           Estimate Std. Error z value Pr(>|z|)
## (Intercept)                4.14231    0.56962   7.272 3.54e-13 ***
## categoryPlastic mixer      0.67075    0.16316   4.111 3.94e-05 ***
## categoryPlastic switcher   0.36459    0.14421   2.528   0.0115 *
## shell.size                 0.09211    0.03740   2.463   0.0138 *
## ---
## Signif. codes:  0 '***' 0.001 '**' 0.01 '*' 0.05 '.' 0.1 ' ' 1
```

```
summary(m2)$coef
```

```
## $cond
##                             Estimate Std. Error  z value     Pr(>|z|)
## (Intercept)               4.14231303 0.56962382 7.272015 3.541639e-13
## categoryPlastic mixer     0.67074509 0.16315520 4.111086 3.938021e-05
## categoryPlastic switcher 0.36459012 0.14420694 2.528243 1.146351e-02
## shell.size                0.09211241 0.03739976 2.462915 1.378127e-02
##
## $zi
## NULL
##
## $disp
## NULL
```

```
# Diagnostic plot
res_m2<-resid(m2)
fitted_m2<-fitted(m2)
plot(res_m2~fitted_m2, las=1, xlab="Fitted values", ylab="Residuals",
     cex.lab=1.6, cex.axis=1.6, cex=1.6)
```

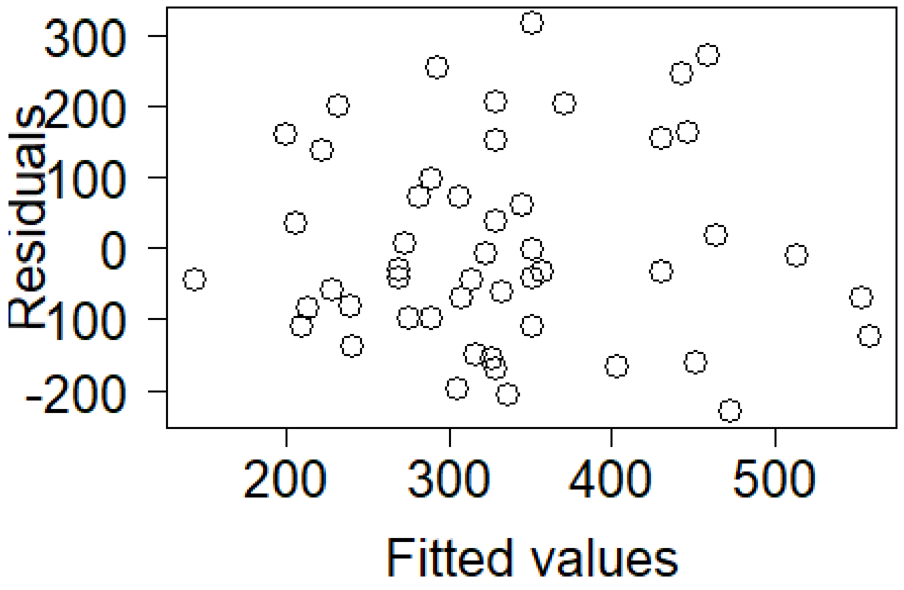

```
# Test the model fit using package DHARMa
  # simulate residuals
simres_m2<-simulateResiduals(fittedModel=m2, plot=F)
simres_m2all<-residuals(simres_m2)
  # make the plots
plot(simres_m2)
```

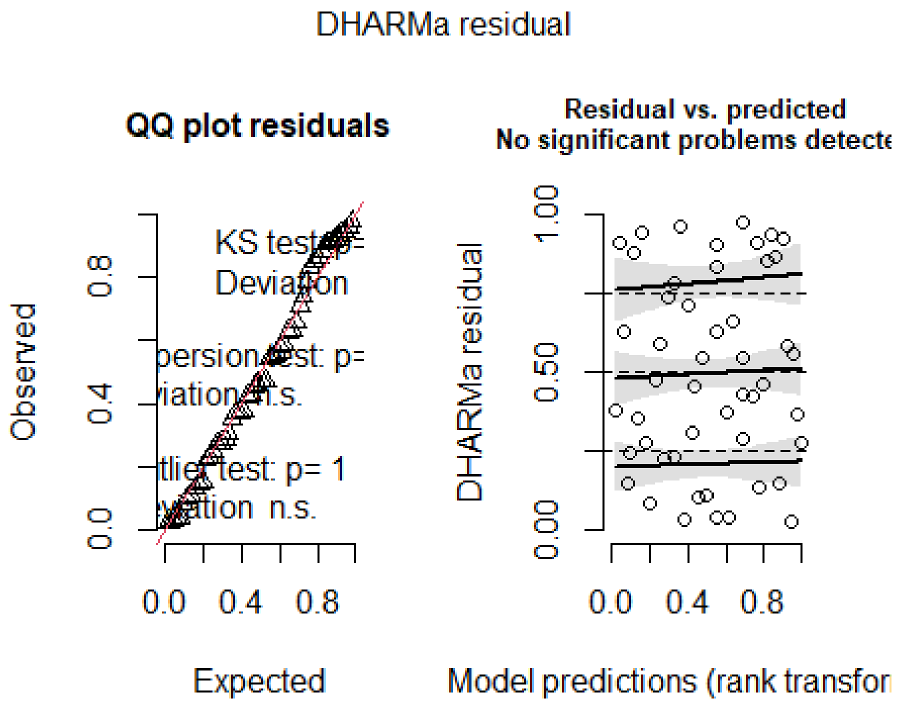

```
par(mfrow=c(2,2), mar=c(5,5,5,4))

# Significance of random effects: P0mother
m2b<-glmmTMB(total.nr.dev.embryos~category + shell.size + (1|pairID), data=d3
, family=nbinom1)
anova(m2, m2b)

## Data: d3
## Models:
## m2b: total.nr.dev.embryos ~ category + shell.size + (1 | pairID), zi=~0, d
isp=~1
## m2: total.nr.dev.embryos ~ category + shell.size + (1 | P0mother) + , zi=~
0, disp=~1
## m2:     (1 | pairID), zi=~0, disp=~1
##     Df    AIC    BIC  logLik deviance Chisq Chi Df Pr(>Chisq)
## m2b  6 638.21 649.68 -313.11   626.21
## m2   7 640.21 653.59 -313.11   626.21     0      1          1

# Significance of random effects: pairID
m2c<-glmmTMB(total.nr.dev.embryos~category + shell.size + (1|P0mother), data=
d3, family=nbinom1)
anova(m2, m2c)

## Data: d3
## Models:
## m2c: total.nr.dev.embryos ~ category + shell.size + (1 | P0mother), zi=~0,
disp=~1
```

```
## m2: total.nr.dev.embryos ~ category + shell.size + (1 | P0mother) + , zi=~
0, disp=~1
## m2:     (1 | pairID), zi=~0, disp=~1
##     Df    AIC    BIC  logLik deviance Chisq Chi Df Pr(>Chisq)
## m2c  6 638.21 649.68 -313.11   626.21
## m2   7 640.21 653.59 -313.11   626.21     0      1          1

# Percentage difference in means
tapply(d3$total.nr.dev.embryos, d3$category, mean)

##           Selfer    Plastic mixer Plastic switcher       Outcrosser
##         244.7333         479.0000         334.2308               NA

((479.0000-244.7333)/479.0000)*100 # selfers produce 48.9% fewer dev. embryos
than plastic mixers

## [1] 48.90745

((334.2308-244.7333)/334.2308)*100 # selfers produce 26.8% fewer dev. embryos
than plastic switchers

## [1] 26.77716
```

#### 9.3 Model 3: Number of eggs in snails with selfing rate estimate (Table S3)

```
m3<-glmmTMB(total.nr.eggs~category + shell.size + (1|P0mother), data=d3, fami
ly=nbinom1)
summary(m3)

##  Family: nbinom1  ( log )
## Formula:          total.nr.eggs ~ category + shell.size + (1 | P0mother)
## Data: d3
##
##      AIC      BIC   logLik deviance df.resid
##    652.9    664.4   -320.5    640.9       44
##
## Random effects:
##
## Conditional model:
##  Groups   Name        Variance  Std.Dev.
##  P0mother (Intercept) 6.122e-10 2.474e-05
## Number of obs: 50, groups:  P0mother, 22
##
## Dispersion parameter for nbinom1 family (): 56.9
##
## Conditional model:
##                           Estimate Std. Error z value Pr(>|z|)
## (Intercept)                4.30234    0.50912   8.450  < 2e-16 ***
## categoryPlastic mixer      0.50635    0.14628   3.462 0.000537 ***
## categoryPlastic switcher   0.27694    0.12640   2.191 0.028453 *
## shell.size                 0.10371    0.03348   3.098 0.001948 **
```

```
## ---
## Signif. codes:  0 '***' 0.001 '**' 0.01 '*' 0.05 '.' 0.1 ' ' 1
```

```
summary(m3)$coef
```

```
## $cond
##                              Estimate Std. Error  z value     Pr(>|z|)
## (Intercept)                 4.3023401 0.50912439 8.450470 2.901335e-17
## categoryPlastic mixer       0.5063489 0.14627531 3.461616 5.369432e-04
## categoryPlastic switcher 0.2769390 0.12639969 2.190979 2.845333e-02
## shell.size                  0.1037089 0.03347504 3.098097 1.947675e-03
##
## $zi
## NULL
##
## $disp
## NULL
```

```
# Diagnostic plot
res_m3<-resid(m3)
fitted_m3<-fitted(m3)
plot(res_m3~fitted_m3, las=1, xlab="Fitted values", ylab="Residuals",
     cex.lab=1.6, cex.axis=1.6, cex=1.6)
```

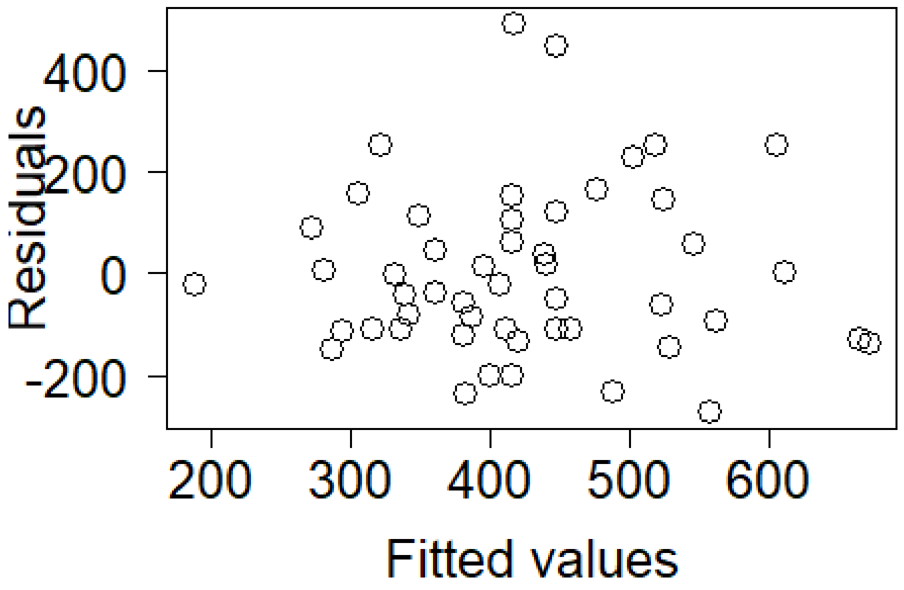

```
# Test the model fit using package DHARMa
  # simulate residuals
simres_m3<-simulateResiduals(fittedModel=m3, plot=F)
```

```
simres_m3all<-residuals(simres_m3)
  # make the plots
plot(simres_m3)
```

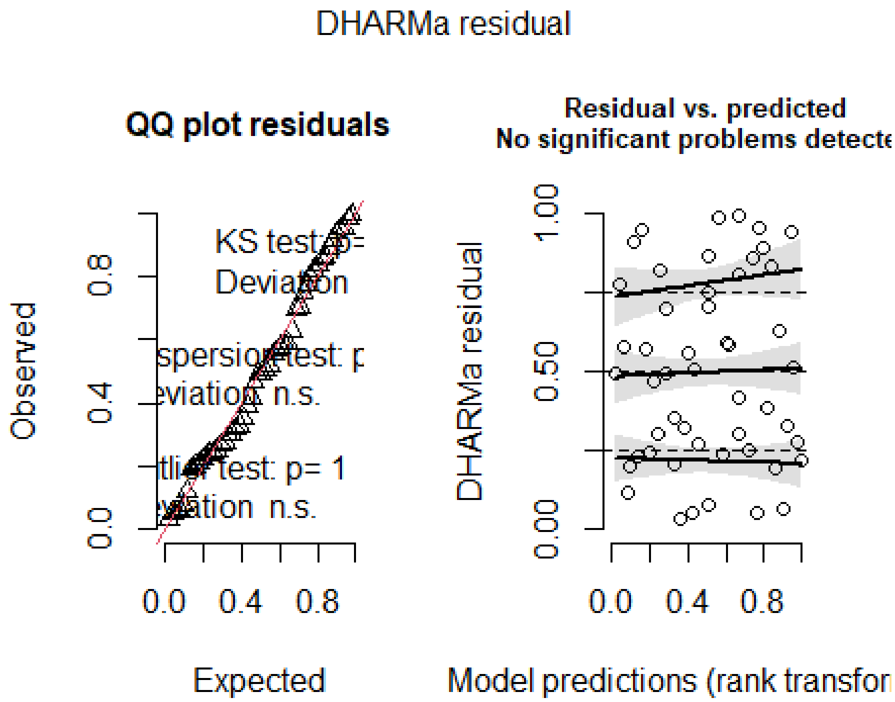

```
par(mfrow=c(2,2), mar=c(5,5,5,4))

# Significance of random effects: P0mother
m3b<-glmmTMB(total.nr.eggs~category + shell.size, data=d3, family=nbinom1)
anova(m3, m3b)

## Data: d3
## Models:
## m3b: total.nr.eggs ~ category + shell.size, zi=~0, disp=~1
## m3: total.nr.eggs ~ category + shell.size + (1 | P0mother), zi=~0, disp=~1
##     Df    AIC    BIC  logLik deviance Chisq Chi Df Pr(>Chisq)
## m3b  5 650.94 660.50 -320.47   640.94
## m3   6 652.94 664.41 -320.47   640.94     0      1          1

# Percentage difference in means
tapply(d3$total.nr.eggs, d3$category, mean)

##           Selfer    Plastic mixer Plastic switcher       Outcrosser
##         341.8667         569.2222         422.4231               NA

((569.2222-341.8667)/569.2222)*100 # selfers produce 39.9% fewer eggs than pl
astic mixers

## [1] 39.94143
```

```
((422.4231-341.8667)/422.4231)*100 # selfers produce 19.1% fewer eggs than pl
astic switchers

## [1] 19.07007
```

#### 9.4 Model 4: Proportion of undeveloped embryos in snails with selfing rate estimate (Table S3)

```
# Assess distribution of dependent variable
ks.test(d3$prop.undev.embryos, pnorm, mean(d3$prop.undev.embryos, na.rm=T), s
d(d3$prop.undev.embryos, na.rm=T)) # D = 0.12566, p-value = 0.3773

##
##  Exact one-sample Kolmogorov-Smirnov test
##
## data:  d3$prop.undev.embryos
## D = 0.12566, p-value = 0.3773
## alternative hypothesis: two-sided

  # no significant deviation from normality
par(mfrow=c(1,2))
hist(d3$prop.undev.embryos, breaks=50, main="", xlab="% undeveloped embryos",
las=1)
qqnorm(d3$prop.undev.embryos, pch = 1, frame = FALSE, main="", las=1)
qqline(d3$prop.undev.embryos, col = "steelblue", lwd = 2)
```

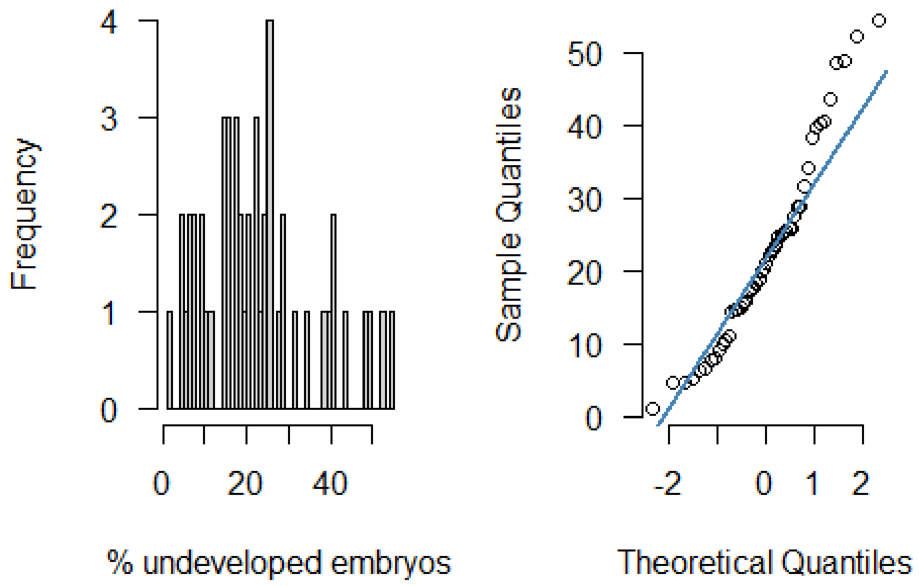

```
# Model
m4<-glmmTMB(prop.undev.embryos~category + shell.size + (1|P0mother) + (1|pair
```

```
ID), data=d3, family=gaussian)
summary(m4)
```

```
##  Family: gaussian  ( identity )
## Formula:
## prop.undev.embryos ~ category + shell.size + (1 | P0mother) +      (1 | pa
irID)
## Data: d3
##
##      AIC      BIC   logLik deviance df.resid
##    407.3    420.7   -196.7    393.3       43
##
## Random effects:
##
## Conditional model:
##  Groups   Name        Variance  Std.Dev.
##  P0mother (Intercept) 2.518e+01  5.017618
##  pairID   (Intercept) 4.807e-06  0.002193
##  Residual             1.318e+02 11.480339
## Number of obs: 50, groups:  P0mother, 22; pairID, 42
##
## Dispersion estimate for gaussian family (sigma^2):  132
##
## Conditional model:
##                            Estimate Std. Error z value Pr(>|z|)
## (Intercept)                 17.5456    16.8911   1.039   0.2989
## categoryPlastic mixer      -12.5874     5.3737  -2.342   0.0192 *
## categoryPlastic switcher    -5.3470     4.2537  -1.257   0.2088
## shell.size                   0.7171     1.1437   0.627   0.5307
## ---
## Signif. codes:  0 '***' 0.001 '**' 0.01 '*' 0.05 '.' 0.1 ' ' 1
```

```
summary(m4)$coef
```

```
## $cond
##                             Estimate Std. Error    z value   Pr(>|z|)
## (Intercept)                17.545645  16.891141  1.0387483 0.29892180
## categoryPlastic mixer     -12.587360   5.373729 -2.3423882 0.01916077
## categoryPlastic switcher   -5.346980   4.253738 -1.2570072 0.20875102
## shell.size                  0.717125   1.143720  0.6270111 0.53065198
##
## $zi
## NULL
##
## $disp
## NULL
```

```
# Diagnostic plot
res_m4<-resid(m4)
fitted_m4<-fitted(m4)
plot(res_m4~fitted_m4, las=1, xlab="Fitted values", ylab="Residuals",
```

```
     cex.lab=1.6, cex.axis=1.6, cex=1.6)

# Test the model fit using package DHARMa
  # simulate residuals
simres_m4<-simulateResiduals(fittedModel=m4, plot=F)
simres_m4all<-residuals(simres_m4)
  # make the plots
plot(simres_m4)
```

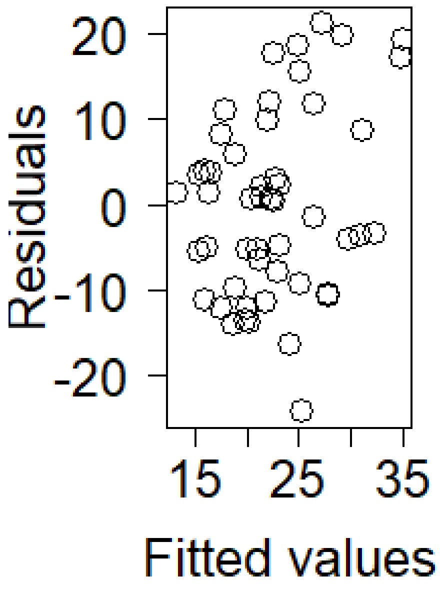

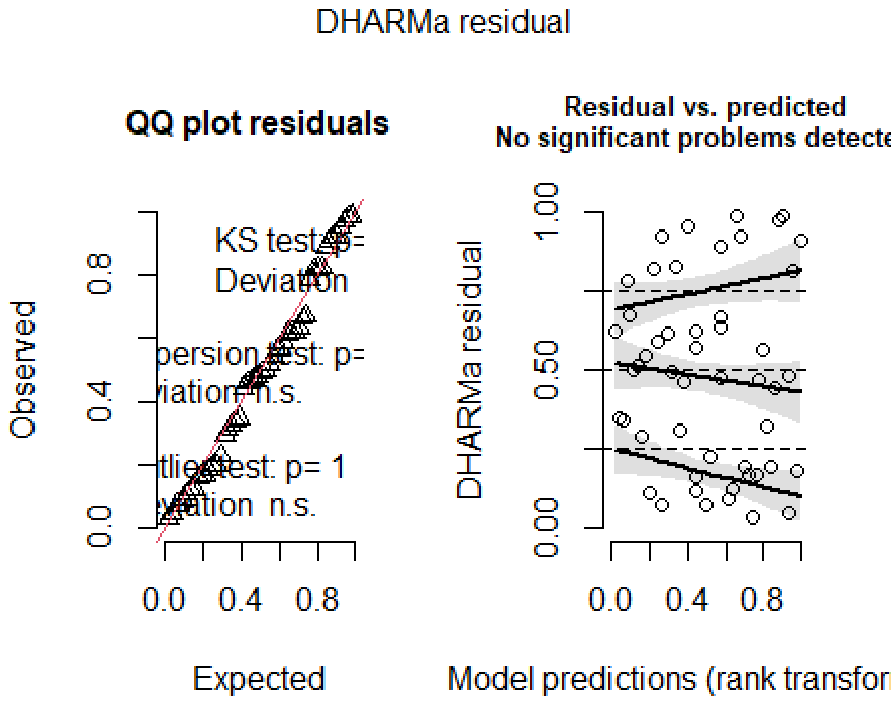

```
par(mfrow=c(2,2), mar=c(5,5,5,4))

# Significance of random effects: P0mother
m4b<-glmmTMB(prop.undev.embryos~category + shell.size + (1|pairID), data=d3,
```

```
family=gaussian)
anova(m4, m4b)

## Data: d3
## Models:
## m4b: prop.undev.embryos ~ category + shell.size + (1 | pairID), zi=~0, dis
p=~1
## m4: prop.undev.embryos ~ category + shell.size + (1 | P0mother) + , zi=~0,
disp=~1
## m4:     (1 | pairID), zi=~0, disp=~1
##      Df    AIC    BIC  logLik deviance  Chisq Chi Df Pr(>Chisq)
## m4b  6 406.27 417.74 -197.13   394.27
## m4   7 407.33 420.71 -196.66   393.33 0.9424      1     0.3317

# Significance of random effects: pairID
m4c<-glmmTMB(prop.undev.embryos~category + shell.size + (1|P0mother), data=d3
, family=gaussian)
anova(m4, m4c)

## Data: d3
## Models:
## m4c: prop.undev.embryos ~ category + shell.size + (1 | P0mother), zi=~0, d
isp=~1
## m4: prop.undev.embryos ~ category + shell.size + (1 | P0mother) + , zi=~0,
disp=~1
## m4:     (1 | pairID), zi=~0, disp=~1
##      Df    AIC    BIC  logLik deviance Chisq Chi Df Pr(>Chisq)
## m4c  6 405.33 416.80 -196.66   393.33
## m4   7 407.33 420.71 -196.66   393.33     0      1          1

# Means of different groups
tapply(d3$prop.undev.embryos, d3$category, mean)

##            Selfer    Plastic mixer Plastic switcher       Outcrosser
##          28.32341         16.22737         21.41214               NA

# Selfer: 28.32, Plastic mixer: 16.23, Plastic switcher: 21.41
```

#### 9.5 Model 5: Number of developed embryos in all female fertile snails (Table 2)

```
# Preparations
s2$P0mother<-as.factor(s2$P0mother)
s2<-merge(s2, partID, by="snailID", all.x=T)
nrow(s2) # 124 control - ok

## [1] 124

s2$pairID<-as.factor(s2$pairID)

# Model
m5<-glmmTMB(total.nr.dev.embryos~cat + shell.size + (1|P0mother) + (1|pairID)
```

```
, data=s2, family=nbinom1)
summary(m5)
```

```
##  Family: nbinom1  ( log )
## Formula:
## total.nr.dev.embryos ~ cat + shell.size + (1 | P0mother) + (1 |      pairI
D)
## Data: s2
##
##      AIC      BIC   logLik deviance df.resid
##   1530.0   1549.7   -758.0   1516.0      116
##
## Random effects:
##
## Conditional model:
##  Groups   Name        Variance  Std.Dev.
##  P0mother (Intercept) 1.195e-01 3.457e-01
##  pairID   (Intercept) 3.458e-09 5.881e-05
## Number of obs: 123, groups:  P0mother, 31; pairID, 88
##
## Dispersion parameter for nbinom1 family ():  128
##
## Conditional model:
##             Estimate Std. Error z value Pr(>|z|)
## (Intercept)  2.59109    0.63042   4.110 3.95e-05 ***
## catplas      1.08012    0.17619   6.130 8.76e-10 ***
## catoutc      0.89174    0.18557   4.805 1.54e-06 ***
## shell.size   0.14136    0.04327   3.267  0.00109 **
## ---
## Signif. codes:  0 '***' 0.001 '**' 0.01 '*' 0.05 '.' 0.1 ' ' 1
```

```
summary(m5)$coef
```

```
## $cond
##              Estimate Std. Error  z value     Pr(>|z|)
## (Intercept) 2.5910877 0.63041682 4.110118 3.954564e-05
## catplas     1.0801240 0.17619054 6.130432 8.764097e-10
## catoutc     0.8917428 0.18557058 4.805411 1.544342e-06
## shell.size  0.1413603 0.04326687 3.267172 1.086277e-03
##
## $zi
## NULL
##
## $disp
## NULL
```

```
# Diagnostic plot
res_m5<-resid(m5)
fitted_m5<-fitted(m5)
plot(res_m5~fitted_m5, las=1, xlab="Fitted values", ylab="Residuals",
     cex.lab=1.6, cex.axis=1.6, cex=1.6)
```

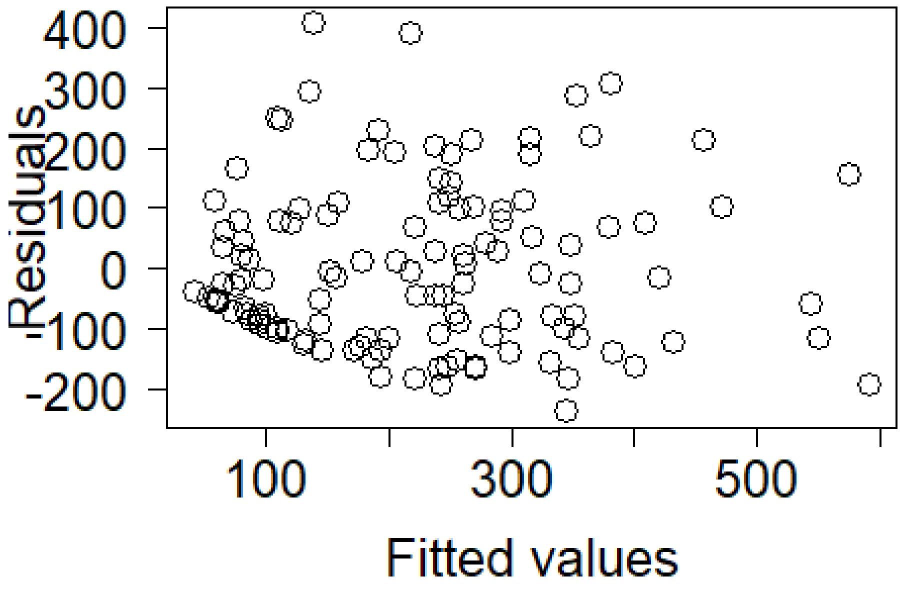

```
# Test the model fit using package DHARMa
  # simulate residuals
simres_m5<-simulateResiduals(fittedModel=m5, plot=F)
simres_m5all<-residuals(simres_m5)
  # make the plots
plot(simres_m5)
```

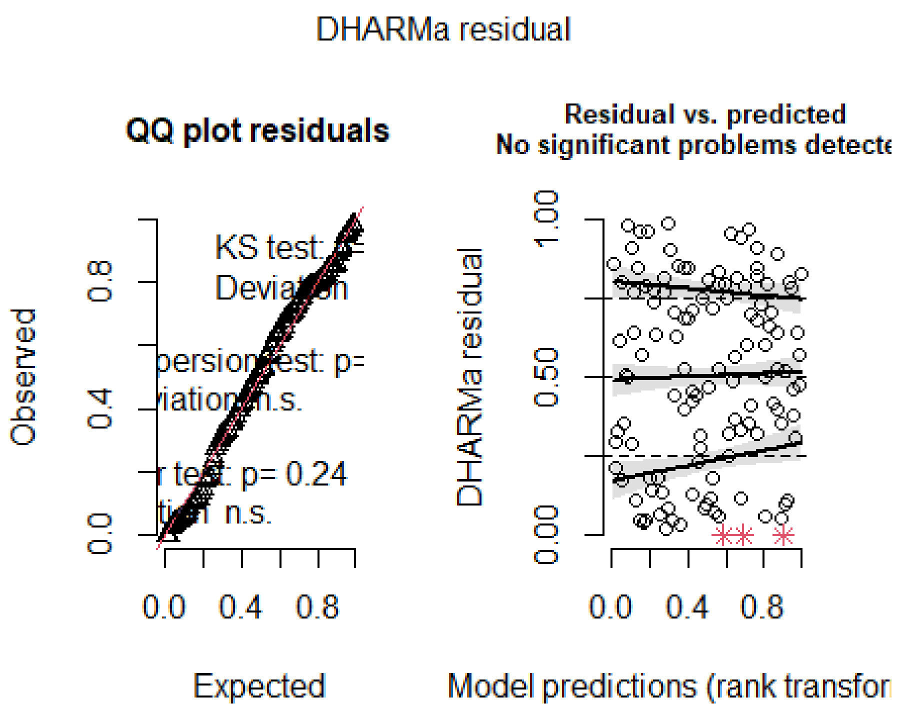

```
par(mfrow=c(2,2), mar=c(5,5,5,4))

# Significance of random effects: P0mother
m5b<-glmmTMB(total.nr.dev.embryos~cat + shell.size + (1|pairID), data=s2, fam
ily=nbinom1)
anova(m5, m5b)

## Data: s2
## Models:
## m5b: total.nr.dev.embryos ~ cat + shell.size + (1 | pairID), zi=~0, disp=~
1
## m5: total.nr.dev.embryos ~ cat + shell.size + (1 | P0mother) + (1 | , zi=~
0, disp=~1
## m5:     pairID), zi=~0, disp=~1
##     Df  AIC    BIC  logLik deviance  Chisq Chi Df Pr(>Chisq)
## m5b  6 1538 1554.9 -763.00     1526
## m5   7 1530 1549.7 -758.01     1516 9.9837      1   0.001579 **
## ---
## Signif. codes:  0 '***' 0.001 '**' 0.01 '*' 0.05 '.' 0.1 ' ' 1

# Significance of random effects: pairID
m5c<-glmmTMB(total.nr.dev.embryos~cat + shell.size + (1|P0mother), data=s2, f
amily=nbinom1)
anova(m5, m5c)

## Data: s2
## Models:
```

```
## m5c: total.nr.dev.embryos ~ cat + shell.size + (1 | P0mother), zi=~0, disp
=~1
## m5: total.nr.dev.embryos ~ cat + shell.size + (1 | P0mother) + (1 | , zi=~
0, disp=~1
## m5:     pairID), zi=~0, disp=~1
##     Df  AIC    BIC  logLik deviance Chisq Chi Df Pr(>Chisq)
## m5c  6 1528 1544.9 -758.01     1516
## m5   7 1530 1549.7 -758.01     1516     0      1          1

# Percentage difference in means
tapply(s2$total.nr.dev.embryos, s2$cat, mean)

##     self     plas     outc
## 113.3500 305.5870 236.1579

((305.5870-113.3500)/305.5870)*100 # selfers produce 62.9% fewer developed em
bryos than plastic snails

## [1] 62.90745

((236.1579-113.3500)/236.1579)*100 # selfers produce 52.0% fewer developed em
bryos than outcrossers

## [1] 52.00245
```

#### 9.6 Model 6: Number of eggs in all female fertile snails (Table 2)

```
m6<-glmmTMB(total.nr.eggs~cat + shell.size + (1|P0mother), data=s2, family=nb
inom1)
summary(m6)

##  Family: nbinom1  ( log )
## Formula:          total.nr.eggs ~ cat + shell.size + (1 | P0mother)
## Data: s2
##
##      AIC      BIC   logLik deviance df.resid
##   1596.8   1613.7   -792.4   1584.8      117
##
## Random effects:
##
## Conditional model:
##  Groups   Name        Variance Std.Dev.
##  P0mother (Intercept) 0.07495  0.2738
## Number of obs: 123, groups:  P0mother, 31
##
## Dispersion parameter for nbinom1 family ():  146
##
## Conditional model:
##             Estimate Std. Error z value Pr(>|z|)
## (Intercept)  2.75243    0.59332   4.639 3.50e-06 ***
## catplas      1.01571    0.16632   6.107 1.02e-09 ***
## catoutc      0.82776    0.17269   4.793 1.64e-06 ***
```

```
## shell.size   0.15454    0.04061   3.805 0.000142 ***
## ---
## Signif. codes:  0 '***' 0.001 '**' 0.01 '*' 0.05 '.' 0.1 ' ' 1
```

```
summary(m6)$coef
```

```
## $cond
##                Estimate Std. Error  z value     Pr(>|z|)
## (Intercept) 2.7524268 0.59331618 4.639056 3.500048e-06
## catplas     1.0157144 0.16631890 6.107029 1.015026e-09
## catoutc     0.8277587 0.17269372 4.793218 1.641269e-06
## shell.size  0.1545441 0.04061359 3.805232 1.416712e-04
##
## $zi
## NULL
##
## $disp
## NULL
```

```
# Diagnostic plot
res_m6<-resid(m6)
fitted_m6<-fitted(m6)
plot(res_m6~fitted_m6, las=1, xlab="Fitted values", ylab="Residuals",
     cex.lab=1.6, cex.axis=1.6, cex=1.6)
```

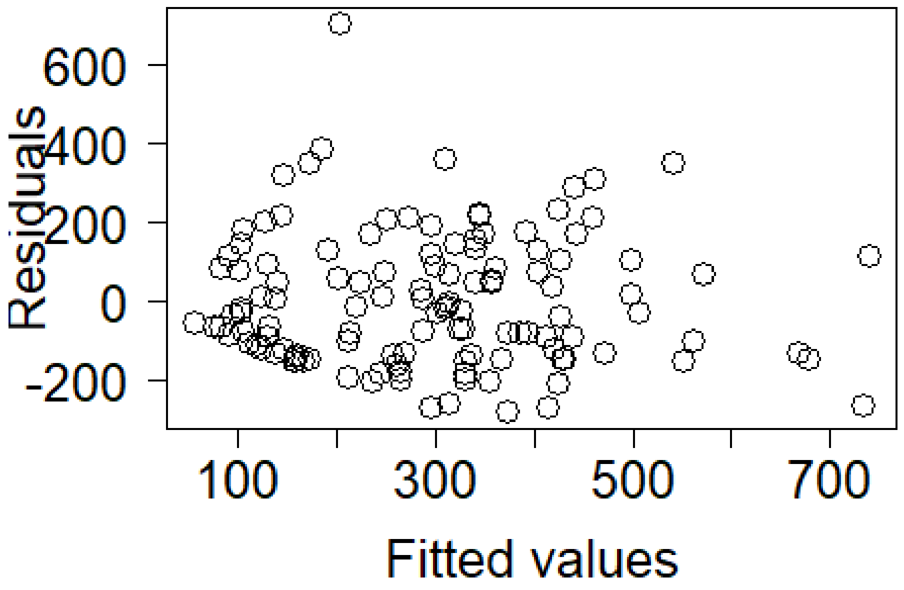

```
# Test the model fit using package DHARMa
  # simulate residuals
```

```
simres_m6<-simulateResiduals(fittedModel=m6, plot=F)
simres_m6all<-residuals(simres_m6)
  # make the plots
plot(simres_m6)
```

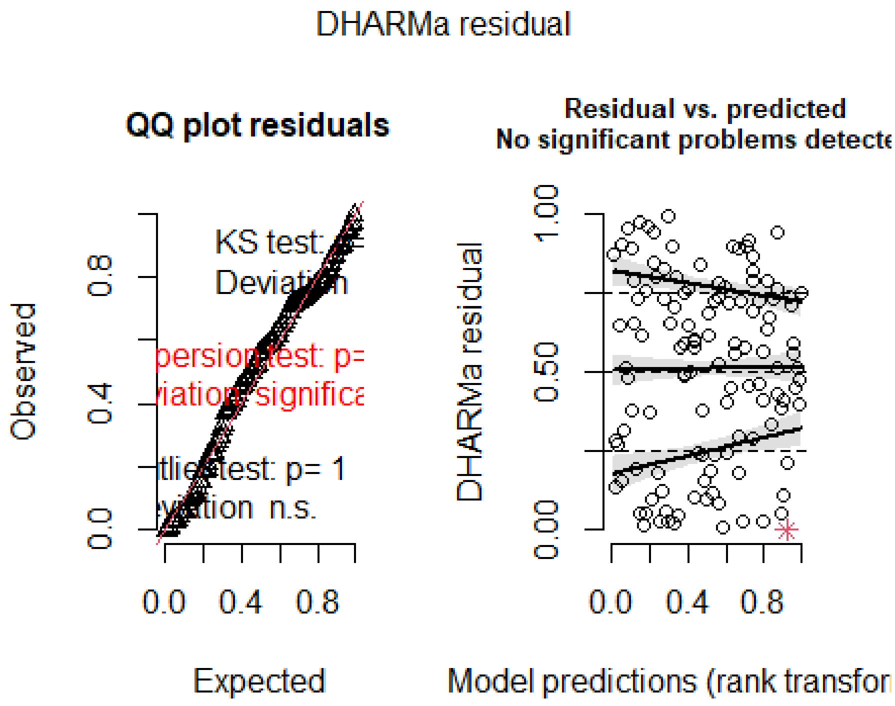

```
par(mfrow=c(2,2), mar=c(5,5,5,4))

# Significance of random effects: P0mother
m6b<-glmmTMB(total.nr.eggs~cat + shell.size, data=s2, family=nbinom1)
anova(m6, m6b)

## Data: s2
## Models:
## m6b: total.nr.eggs ~ cat + shell.size, zi=~0, disp=~1
## m6: total.nr.eggs ~ cat + shell.size + (1 | P0mother), zi=~0, disp=~1
##     Df    AIC    BIC  logLik deviance  Chisq Chi Df Pr(>Chisq)
## m6b  5 1600.4 1614.5 -795.21   1590.4
## m6   6 1596.8 1613.7 -792.42   1584.8 5.5881      1    0.01808 *
## ---
## Signif. codes:  0 '***' 0.001 '**' 0.01 '*' 0.05 '.' 0.1 ' ' 1

# Percentage difference in means
tapply(s2$total.nr.eggs, s2$cat, mean)

##     self     plas     outc
## 168.2000 390.8043 305.1316
```

```
((390.8043-168.2000)/390.8043)*100 # selfers produce 57.0% fewer eggs than pl
astic snails

## [1] 56.96056

((305.1316-168.2000)/305.1316)*100 # selfers produce 44.9% fewer eggs than ou
tcrossers

## [1] 44.87624
```

#### 9.7 Model 7: Proportion of undeveloped embryos in all female fertile snails (Table 2)

```
# Assess distribution of dependent variable
ks.test(s2$prop.undev.embryos, pnorm, mean(s2$prop.undev.embryos, na.rm=T), s
d(s2$prop.undev.embryos, na.rm=T)) # D = 0.084847, p-value = 0.3339

## Warning in ks.test.default(s2$prop.undev.embryos, pnorm,
## mean(s2$prop.undev.embryos, : ties should not be present for the
## Kolmogorov-Smirnov test

##
##  Asymptotic one-sample Kolmogorov-Smirnov test
##
## data:  s2$prop.undev.embryos
## D = 0.084847, p-value = 0.3339
## alternative hypothesis: two-sided

  # no significant deviation from normality
par(mfrow=c(1,2))
hist(s2$prop.undev.embryos, breaks=50, main="", xlab="% undeveloped embryos",
las=1)
qqnorm(s2$prop.undev.embryos, pch = 1, frame = FALSE, main="", las=1)
qqline(s2$prop.undev.embryos, col = "steelblue", lwd = 2)
```

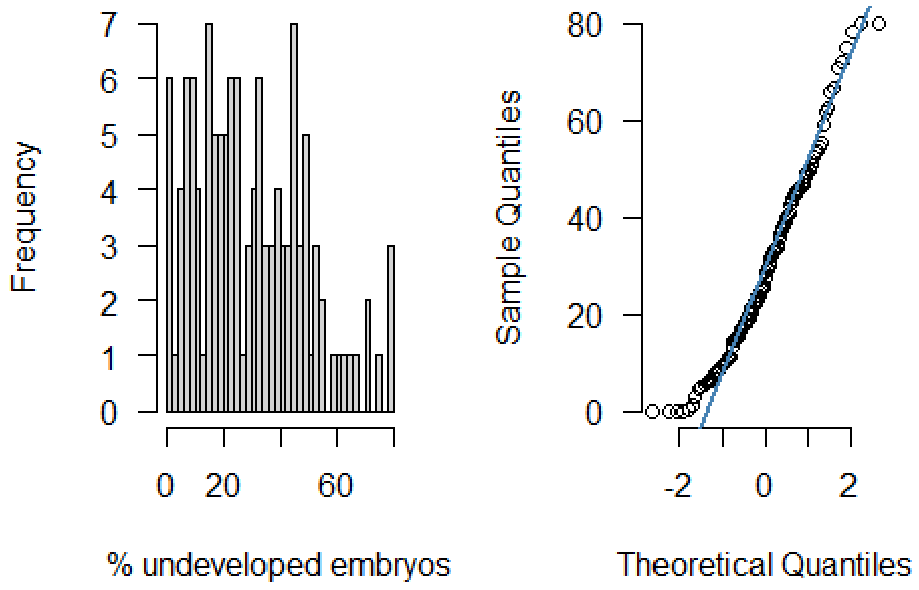

```
# Model
m7<-glmmTMB(prop.undev.embryos~cat + shell.size + (1|P0mother) + (1|pairID),
data=s2, family=gaussian)
summary(m7)
```

```
##  Family: gaussian  ( identity )
## Formula:
## prop.undev.embryos ~ cat + shell.size + (1 | P0mother) + (1 |      pairID)
## Data: s2
##
##      AIC      BIC   logLik deviance df.resid
##   1072.2   1091.9   -529.1   1058.2      116
##
## Random effects:
##
## Conditional model:
##  Groups   Name        Variance Std.Dev.
##  P0mother (Intercept) 103.36   10.167
##  pairID   (Intercept)  12.95    3.599
##  Residual             247.38   15.728
## Number of obs: 123, groups:  P0mother, 31; pairID, 88
##
## Dispersion estimate for gaussian family (sigma^2):  247
##
## Conditional model:
##             Estimate Std. Error z value Pr(>|z|)
## (Intercept)  30.7793    14.8665   2.070 0.038417 *
```

```
## catplas     -12.7675     3.8045  -3.356 0.000791 ***
## catoutc     -11.1430     3.9235  -2.840 0.004510 **
## shell.size    0.4736     1.0665   0.444 0.656973
## ---
## Signif. codes:  0 '***' 0.001 '**' 0.01 '*' 0.05 '.' 0.1 ' ' 1
```

```
summary(m7)$coef
```

```
## $cond
##                Estimate Std. Error    z value     Pr(>|z|)
## (Intercept)  30.7793474  14.866537  2.0703777 0.0384169847
## catplas     -12.7675181   3.804544 -3.3558602 0.0007911855
## catoutc     -11.1430457   3.923456 -2.8401097 0.0045098028
## shell.size    0.4736428   1.066532  0.4440961 0.6569730758
##
## $zi
## NULL
##
## $disp
## NULL
```

```
# Diagnostic plot
res_m7<-resid(m7)
fitted_m7<-fitted(m7)
plot(res_m7~fitted_m7, las=1, xlab="Fitted values", ylab="Residuals",
     cex.lab=1.6, cex.axis=1.6, cex=1.6)

# Test the model fit using package DHARMa
  # simulate residuals
simres_m7<-simulateResiduals(fittedModel=m7, plot=F)
simres_m7all<-residuals(simres_m7)
  # make the plots
plot(simres_m7)
```

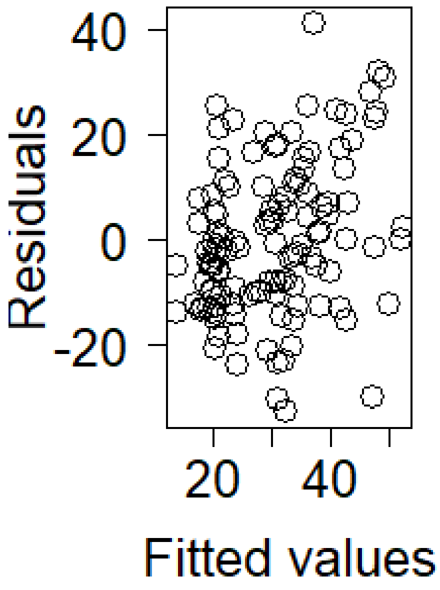

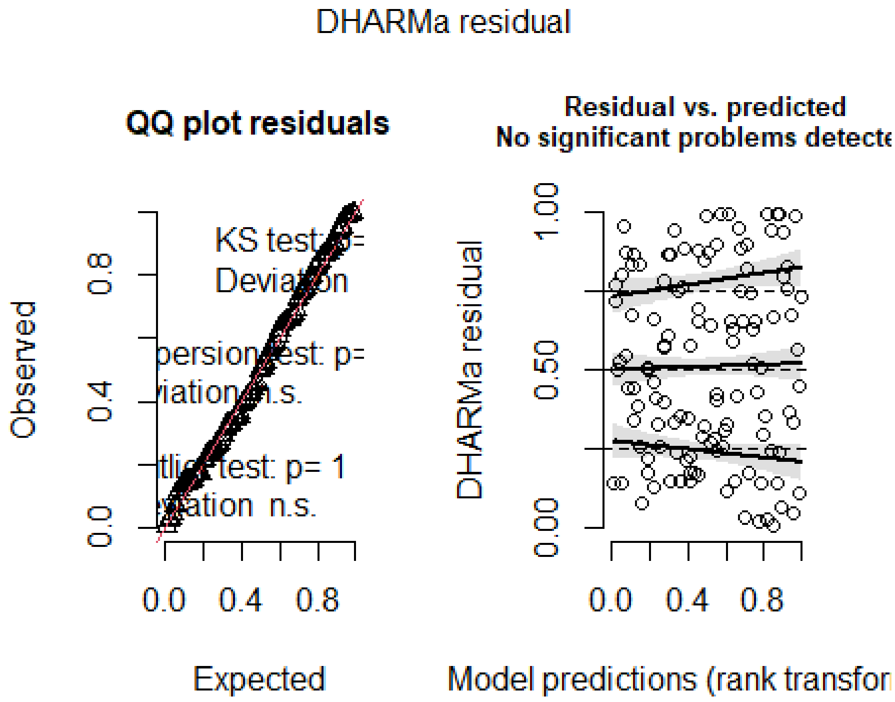

```
par(mfrow=c(2,2), mar=c(5,5,5,4))

# Significance of random effects: P0mother
m7b<-glmmTMB(prop.undev.embryos~cat + shell.size + (1|pairID), data=s2, famil
```

```
y=gaussian)
anova(m7, m7b)

## Data: s2
## Models:
## m7b: prop.undev.embryos ~ cat + shell.size + (1 | pairID), zi=~0, disp=~1
## m7: prop.undev.embryos ~ cat + shell.size + (1 | P0mother) + (1 | , zi=~0,
disp=~1
## m7:     pairID), zi=~0, disp=~1
##     Df    AIC    BIC  logLik deviance  Chisq Chi Df Pr(>Chisq)
## m7b  6 1079.6 1096.5 -533.79   1067.6
## m7   7 1072.2 1091.9 -529.10   1058.2 9.3799      1   0.002194 **
## ---
## Signif. codes:  0 '***' 0.001 '**' 0.01 '*' 0.05 '.' 0.1 ' ' 1

# Significance of random effects: pairID
m7c<-glmmTMB(prop.undev.embryos~cat + shell.size + (1|P0mother), data=s2, fam
ily=gaussian)
anova(m7, m7c)

## Data: s2
## Models:
## m7c: prop.undev.embryos ~ cat + shell.size + (1 | P0mother), zi=~0, disp=~
1
## m7: prop.undev.embryos ~ cat + shell.size + (1 | P0mother) + (1 | , zi=~0,
disp=~1
## m7:     pairID), zi=~0, disp=~1
##     Df    AIC    BIC  logLik deviance Chisq Chi Df Pr(>Chisq)
## m7c  6 1070.3 1087.1 -529.13   1058.3
## m7   7 1072.2 1091.9 -529.10   1058.2  0.06      1     0.8066

# Means of different groups
tapply(s2$prop.undev.embryos, s2$cat, mean)

##     self     plas     outc
## 36.79047 25.36906 28.24343

# self: 36.79047, plas: 25.36906, outc: 28.24343
```

### 10. Models 8-9: Do snails with different selfing propensity differ in their offsprings' juvenile survival?

#### 10.1 Model 8: Proportion of juveniles that died in snails with selfing rate estimate (Table S3)

```
# Assess distribution of dependent variable
ks.test(d3$mort.rate, pnorm, mean(d3$mort.rate, na.rm=T), sd(d3$mort.rate, na
.rm=T)) # D = 0.071501, p-value = 0.9442
```

```
##
##  Exact one-sample Kolmogorov-Smirnov test
##
## data:  d3$mort.rate
## D = 0.071501, p-value = 0.9442
## alternative hypothesis: two-sided
```

```
  # no significant deviation from normality
par(mfrow=c(1,2))
hist(d3$mort.rate, breaks=30, main="", xlab="% dead juveniles", las=1)
qqnorm(d3$mort.rate, pch = 1, frame = FALSE, main="", las=1)
qqline(d3$mort.rate, col = "steelblue", lwd = 2)
```

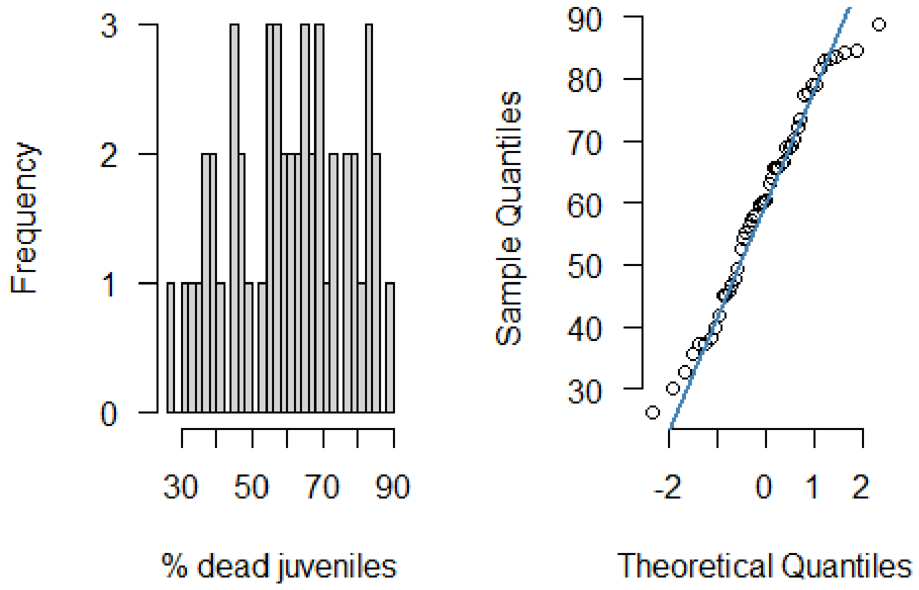

```
# Model
m8<-glmmTMB(mort.rate~category + shell.size + (1|P0mother) + (1|pairID), data
=d3, family=gaussian)
summary(m8)
```

```
##  Family: gaussian  ( identity )
## Formula:
## mort.rate ~ category + shell.size + (1 | P0mother) + (1 | pairID)
## Data: d3
##
##      AIC      BIC   logLik deviance df.resid
##    414.0    427.4   -200.0    400.0       43
##
## Random effects:
```

```
##
## Conditional model:
##  Groups   Name        Variance  Std.Dev.
##  P0mother (Intercept) 1.060e-06  0.001029
##  pairID   (Intercept) 6.682e+01  8.174449
##  Residual             1.121e+02 10.588078
## Number of obs: 50, groups:  P0mother, 22; pairID, 42
##
## Dispersion estimate for gaussian family (sigma^2):  112
##
## Conditional model:
##                           Estimate Std. Error z value Pr(>|z|)
## (Intercept)                116.783     18.039   6.474 9.55e-11 ***
## categoryPlastic mixer       -3.403      5.526  -0.616  0.53795
## categoryPlastic switcher   -17.756      4.490  -3.955 7.66e-05 ***
## shell.size                  -3.301      1.225  -2.695  0.00704 **
## ---
## Signif. codes:  0 '***' 0.001 '**' 0.01 '*' 0.05 '.' 0.1 ' ' 1
```

```
summary(m8)$coef
```

```
## $cond
##                             Estimate Std. Error    z value     Pr(>|z|)
## (Intercept)               116.782699  18.039037  6.4738878 9.551283e-11
## categoryPlastic mixer      -3.403459   5.525878 -0.6159129 5.379520e-01
## categoryPlastic switcher -17.755542   4.489784 -3.9546539 7.664553e-05
## shell.size                 -3.300534   1.224779 -2.6948011 7.043066e-03
##
## $zi
## NULL
##
## $disp
## NULL
```

```
# Diagnostic plot
res_m8<-resid(m8)
fitted_m8<-fitted(m8)
plot(res_m8~fitted_m8, las=1, xlab="Fitted values", ylab="Residuals",
     cex.lab=1.6, cex.axis=1.6, cex=1.6)

# Test the model fit using package DHARMa
  # simulate residuals
simres_m8<-simulateResiduals(fittedModel=m8, plot=F)
simres_m8all<-residuals(simres_m8)
  # make the plots
plot(simres_m8)
```

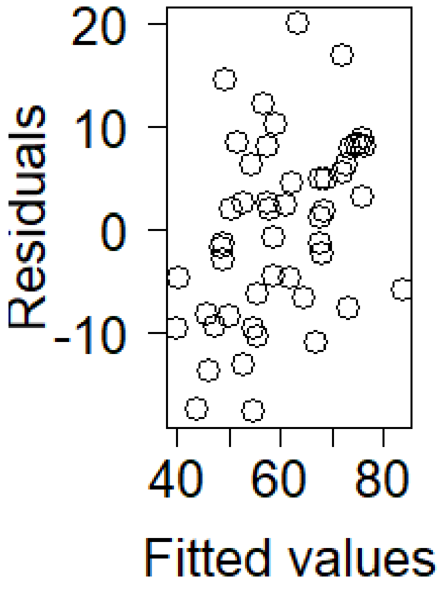

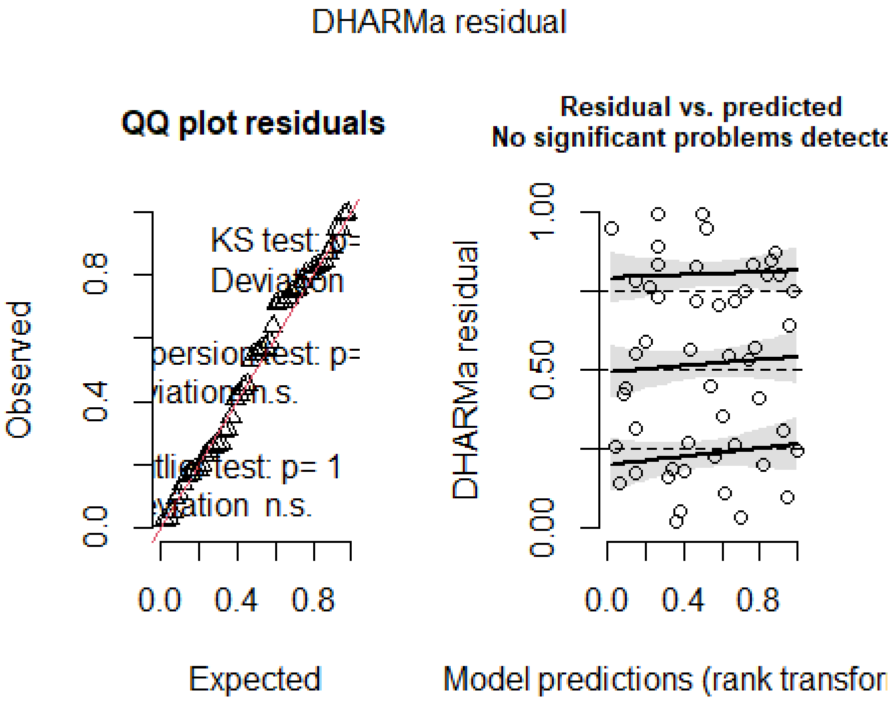

```
par(mfrow=c(2,2), mar=c(5,5,5,4))

# Significance of random effects: P0mother
m8b<-glmmTMB(mort.rate~category + shell.size + (1|pairID), data=d3, family=ga
```

```
ussian)
anova(m8, m8b)

## Data: d3
## Models:
## m8b: mort.rate ~ category + shell.size + (1 | pairID), zi=~0, disp=~1
## m8: mort.rate ~ category + shell.size + (1 | P0mother) + (1 | pairID), zi=
~0, disp=~1
##     Df    AIC    BIC  logLik deviance Chisq Chi Df Pr(>Chisq)
## m8b  6 412.04 423.51 -200.02   400.04
## m8   7 414.04 427.43 -200.02   400.04     0      1          1

# Significance of random effects: pairID
m8c<-glmmTMB(mort.rate~category + shell.size + (1|P0mother), data=d3, family=
gaussian)
anova(m8, m8c)

## Data: d3
## Models:
## m8c: mort.rate ~ category + shell.size + (1 | P0mother), zi=~0, disp=~1
## m8: mort.rate ~ category + shell.size + (1 | P0mother) + (1 | pairID), zi=
~0, disp=~1
##     Df    AIC    BIC  logLik deviance Chisq Chi Df Pr(>Chisq)
## m8c  6 413.21 424.68 -200.60   401.21
## m8   7 414.04 427.43 -200.02   400.04 1.165      1     0.2804

# Means of different groups
tapply(d3$mort.rate, d3$category, mean)

##            Selfer    Plastic mixer Plastic switcher       Outcrosser
##          69.45365         64.54469         53.41986               NA

# Selfer: 69.45365 , Plastic mixer: 64.54469, Plastic switcher: 53.41986
```

#### 10.2 Model 9: Proportion of juveniles that died in all female fertile snails (Table 2)

```
# Assess distribution of dependent variable
ks.test(s2$mort.rate, pnorm, mean(s2$mort.rate, na.rm=T), sd(s2$mort.rate, na
.rm=T)) # D = 0.057596, p-value = 0.8439

## Warning in ks.test.default(s2$mort.rate, pnorm, mean(s2$mort.rate, na.rm =
T),
## : ties should not be present for the Kolmogorov-Smirnov test

##
##  Asymptotic one-sample Kolmogorov-Smirnov test
##
## data:  s2$mort.rate
## D = 0.057596, p-value = 0.8439
## alternative hypothesis: two-sided
```

```
  # no significant deviation from normality
par(mfrow=c(1,2))
hist(s2$mort.rate, breaks=30, main="", xlab="% dead juveniles", las=1)
qqnorm(s2$mort.rate, pch = 1, frame = FALSE, main="", las=1)
qqline(s2$mort.rate, col = "steelblue", lwd = 2)
```

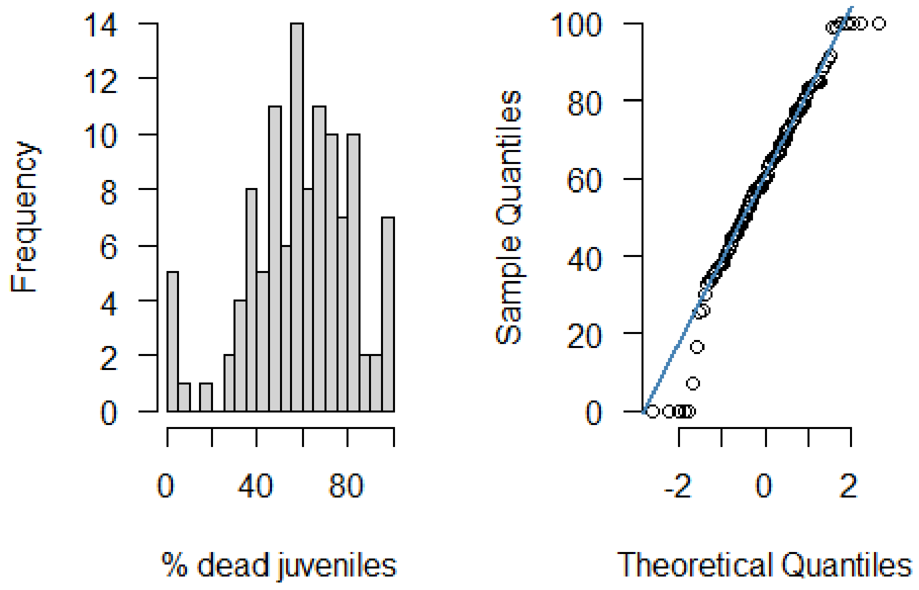

```
# Model
m9<-glmmTMB(mort.rate~cat + shell.size + (1|P0mother) + (1|pairID), data=s2,
family=gaussian)
summary(m9)
```

```
##  Family: gaussian  ( identity )
## Formula:          mort.rate ~ cat + shell.size + (1 | P0mother) + (1 | pai
rID)
## Data: s2
##
##      AIC      BIC   logLik deviance df.resid
##   1022.2   1041.3   -504.1   1008.2      106
##
## Random effects:
##
## Conditional model:
##  Groups   Name        Variance  Std.Dev.
##  P0mother (Intercept) 9.570e-10 3.094e-05
##  pairID   (Intercept) 1.590e+02 1.261e+01
##  Residual             2.968e+02 1.723e+01
## Number of obs: 113, groups:  P0mother, 30; pairID, 80
```

```
##
## Dispersion estimate for gaussian family (sigma^2):  297
##
## Conditional model:
##              Estimate Std. Error z value Pr(>|z|)
## (Intercept)  106.086     19.508   5.438 5.38e-08 ***
## catplas       -7.800      4.942  -1.578  0.11451
## catoutc      -16.693      5.120  -3.261  0.00111 **
## shell.size    -2.710      1.380  -1.964  0.04956 *
## ---
## Signif. codes:  0 '***' 0.001 '**' 0.01 '*' 0.05 '.' 0.1 ' ' 1
```

```
summary(m9)$coef
```

```
## $cond
##                Estimate Std. Error   z value     Pr(>|z|)
## (Intercept) 106.085895   19.507873  5.438107 5.384975e-08
## catplas      -7.799799    4.942117 -1.578230 1.145127e-01
## catoutc     -16.692881    5.119560 -3.260609 1.111734e-03
## shell.size   -2.709560    1.379791 -1.963747 4.955946e-02
##
## $zi
## NULL
##
## $disp
## NULL
```

```
# Diagnostic plot
res_m9<-resid(m9)
fitted_m9<-fitted(m9)
plot(res_m9~fitted_m9, las=1, xlab="Fitted values", ylab="Residuals",
     cex.lab=1.6, cex.axis=1.6, cex=1.6)

# Test the model fit using package DHARMa
  # simulate residuals
simres_m9<-simulateResiduals(fittedModel=m9, plot=F)
simres_m9all<-residuals(simres_m9)
  # make the plots
plot(simres_m9)
```

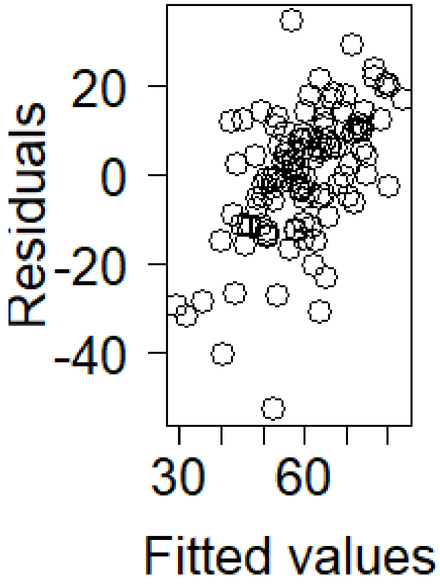

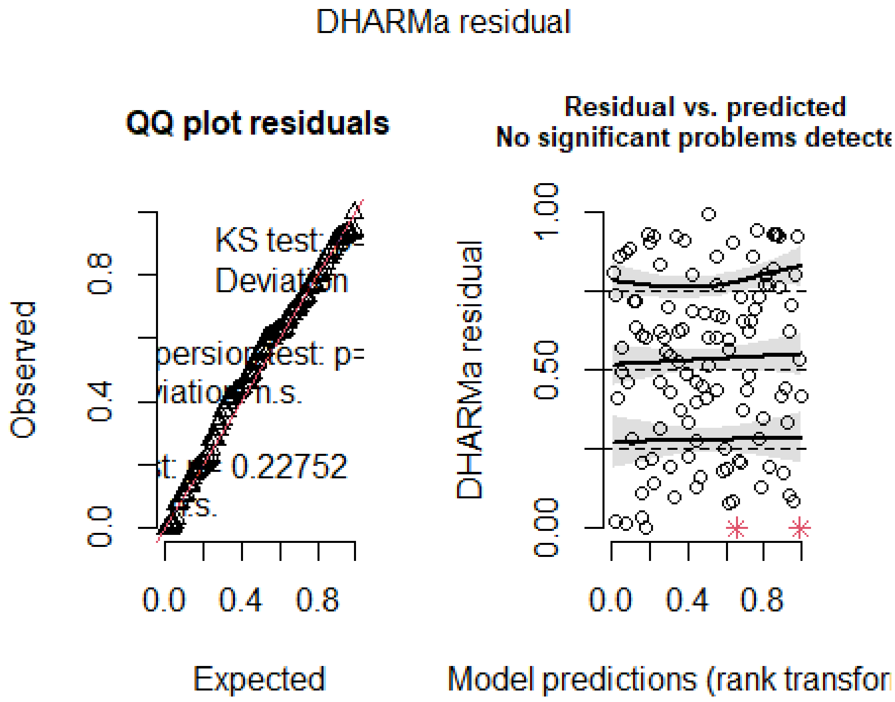

```
par(mfrow=c(2,2), mar=c(5,5,5,4))

# Significance of random effects: P0mother
m9b<-glmmTMB(mort.rate~cat + shell.size + (1|pairID), data=s2, family=gaussia
```

```
n)
anova(m9, m9b)

## Data: s2
## Models:
## m9b: mort.rate ~ cat + shell.size + (1 | pairID), zi=~0, disp=~1
## m9: mort.rate ~ cat + shell.size + (1 | P0mother) + (1 | pairID), zi=~0, d
isp=~1
##     Df    AIC    BIC  logLik deviance Chisq Chi Df Pr(>Chisq)
## m9b  6 1020.2 1036.5 -504.09   1008.2
## m9   7 1022.2 1041.3 -504.09   1008.2     0      1          1

# Significance of random effects: pairID
m9c<-glmmTMB(mort.rate~cat + shell.size + (1|P0mother), data=s2, family=gauss
ian)
anova(m9, m9c)

## Data: s2
## Models:
## m9c: mort.rate ~ cat + shell.size + (1 | P0mother), zi=~0, disp=~1
## m9: mort.rate ~ cat + shell.size + (1 | P0mother) + (1 | pairID), zi=~0, d
isp=~1
##     Df    AIC    BIC  logLik deviance  Chisq Chi Df Pr(>Chisq)
## m9c  6 1022.7 1039.1 -505.35   1010.7
## m9   7 1022.2 1041.3 -504.09   1008.2 2.5094      1     0.1132

# Means of different groups
tapply(s2$mort.rate, s2$cat, mean, na.rm=T)

##     self     plas     outc
## 65.51850 60.90746 51.64772

# apparent selfer: 65.51850 , apparent plastic snail: 60.90746, apparent outc
rosser: 51.64772
```

### 11. Figure 4

#### 11.1 Preparations

```
# Make column specifying whether snails mated in both sexual roles
d2$cop.both<-ifelse(d2$cop.type %in% "both", 1, 0)
table(d2$cop.both, d2$cop.type)

##
##     both never only fem only male
##   0    0     1        7         7
##   1   40     0        0         0

table(d2$cop.both, d2$category)
```

```
##
##     Selfer Plastic mixer Plastic switcher Outcrosser
##   0      9             3                2          1
##   1      6             6               24          4

d3$cop.both<-ifelse(d3$cop.type %in% "both", 1, 0)
table(d3$cop.both, d3$cop.type)

##
##     both never only fem only male
##   0    0     1        6         7
##   1   36     0        0         0

table(d3$cop.both, d3$category)

##
##     Selfer Plastic mixer Plastic switcher Outcrosser
##   0      9             3                2          0
##   1      6             6               24          0

# Compute 95% confidence intervals for proportions for Fig. 4
x<-c(6, 6, 24, 4)
n<-c(15, 9, 26, 5)
binom95CI(x, n)

## [1] 0.1633643 0.2992951 0.7486971 0.2835821 0.6771302 0.9251454 0.9905446
## [8] 0.9949492

errorbars_fig4<-binom95CI(x, n)

# How many selfers mated as a female, how many as a male?
table(d2$category, d2$cop.type)

##
##                       both never only fem only male
##   Selfer                 6     1        2         6
##   Plastic mixer          6     0        3         0
##   Plastic switcher      24     0        1         1
##   Outcrosser             4     0        1         0

(6+2)/(6+1+2+6) # 0.5333333, i.e. 53.3% mated as a female

## [1] 0.5333333

(6+6)/(6+1+2+6) # 0.8, i.e. 80.0% mated as a male

## [1] 0.8

# How many snails with non-zero selfing rates post-isolation mated as a femal
e?
# I.e., how many selfers and plastic mixers mated as a female?
table(d2$category, d2$cop.type)
```

```
##
##                       both never only fem only male
##   Selfer                 6     1        2         6
##   Plastic mixer          6     0        3         0
##   Plastic switcher      24     0        1         1
##   Outcrosser             4     0        1         0
```

```
(6+2+6+3)/(6+1+2+6+6+0+3+0) # 0.7083333,i.e. 70.8% of selfers and plastic mix
ers mated as a female

## [1] 0.7083333
```

#### 11.2 Figure 4

```
#tiff(filename="Fig4.tif", width=85, height=85, units="mm",
#      res=600, pointsize=11)

par(mar=c(4.5,3.5,2,0), xpd=T)
par(bty="n")
a<-barplot(height=c(6/(6+1+2+6), 6/(6+3), 24/(24+1+1), 4/(4+1)),
        beside=TRUE, xlab="", col=c("red", "lightskyblue4", "lightskyblue1",
"green1"),
        ylab="", cex.axis=1, cex.lab=1, cex.names=1, las=1,
        mgp=c(3.5,1,0), xaxt="n", yaxt="n", space=c(0.2))

title(ylab="% snails that mated in both roles", mgp=c(2.2, 1, 0), cex.lab=1)
title(xlab="Propensity for selfing", mgp=c(3.5, 1, 0), cex.lab=1)
axis(1, tick=F, at=c(0.7, 1.9, 3.1, 4.3), labels=c("Selfer\n\n",
                                        "Plastic\nmixer\n",
                                        "Plastic\nswit-\ncher",
                                        "Out-\ncros-\nser"),
     cex.axis=1, las=1, mgp=(c(2, 2, 0)))
axis(2, tick=T, at=seq(0, 1, 0.2), labels=c("0.0", "0.2", "0.4", "0.6", "0.8"
, "1.0"),
     cex.axis=1, las=1, mgp=c(2,0.75,0))
segments(a, errorbars_fig4[1:4], a, errorbars_fig4[5:8], lwd=1)
arrows(a, errorbars_fig4[1:4], a, errorbars_fig4[5:8], lwd=1, angle=90, code=
3, length=0.05)
```

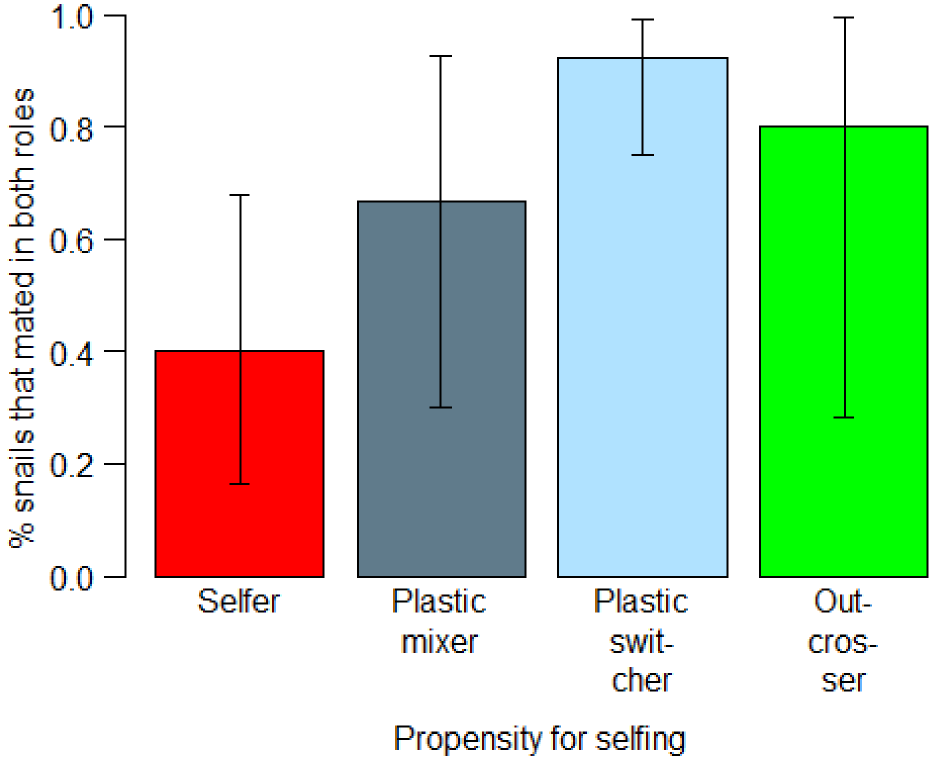

```
#dev.off()
```

### 12. Model 10: Do snails that mated in both roles differ in their selfing propensity?

#### 12.1 Model 10: Probability of mating in both sexual roles

```
m10<-glmmTMB(cop.both~category + shell.size + (1|P0mother), data=d3, family=b
inomial)
summary(m10)

##  Family: binomial  ( logit )
## Formula:          cop.both ~ category + shell.size + (1 | P0mother)
## Data: d3
##
##      AIC      BIC   logLik deviance df.resid
##     55.7     65.3    -22.8     45.7       45
##
## Random effects:
##
## Conditional model:
##  Groups   Name        Variance  Std.Dev.
##  P0mother (Intercept) 2.769e-09 5.263e-05
## Number of obs: 50, groups:  P0mother, 22
##
## Conditional model:
```

```
##                             Estimate Std. Error z value Pr(>|z|)
## (Intercept)                  0.32576    3.08912   0.105  0.91602
## categoryPlastic mixer        1.11967    0.88754   1.262  0.20712
## categoryPlastic switcher     2.88076    0.90618   3.179  0.00148 **
## shell.size                  -0.05095    0.21242  -0.240  0.81042
## ---
## Signif. codes:  0 '***' 0.001 '**' 0.01 '*' 0.05 '.' 0.1 ' ' 1
```

```
summary(m10)$coef
```

```
## $cond
##                               Estimate Std. Error    z value    Pr(>|z|)
## (Intercept)                  0.32576242  3.0891250  0.1054546 0.916015097
## categoryPlastic mixer        1.11966797  0.8875432  1.2615364 0.207115664
## categoryPlastic switcher     2.88076067  0.9061844  3.1790005 0.001477838
## shell.size                  -0.05095445  0.2124175 -0.2398787 0.810424269
##
## $zi
## NULL
##
## $disp
## NULL
```

```
# Diagnostic plot
res_m10<-resid(m10)
fitted_m10<-fitted(m10)
plot(res_m10~fitted_m10, las=1, xlab="Fitted values", ylab="Residuals",
     cex.lab=1.6, cex.axis=1.6, cex=1.6)
```

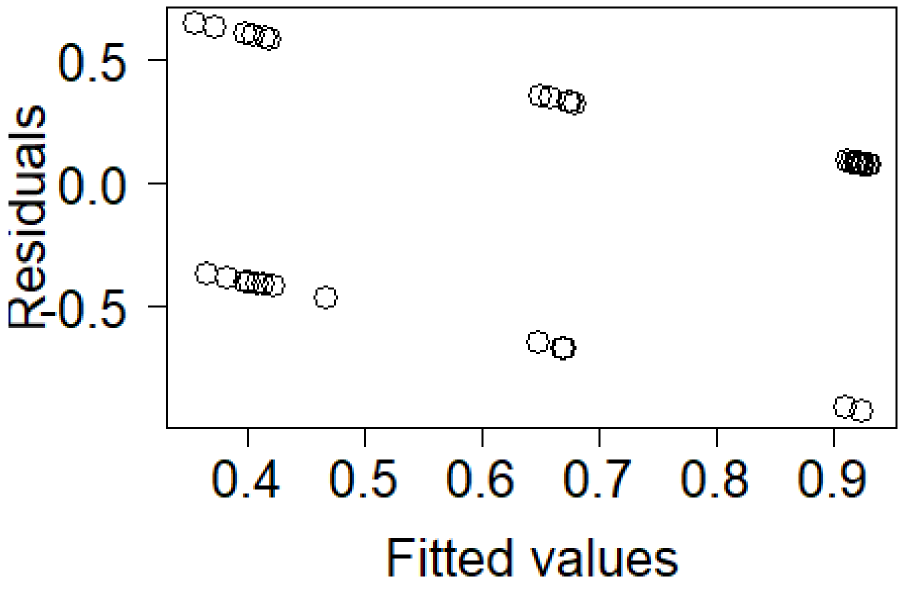

```
# Test the model fit using package DHARMa
  # simulate residuals
simres_m10<-simulateResiduals(fittedModel=m10, plot=F)
simres_m10all<-residuals(simres_m10)
  # make the plots
plot(simres_m10)
```

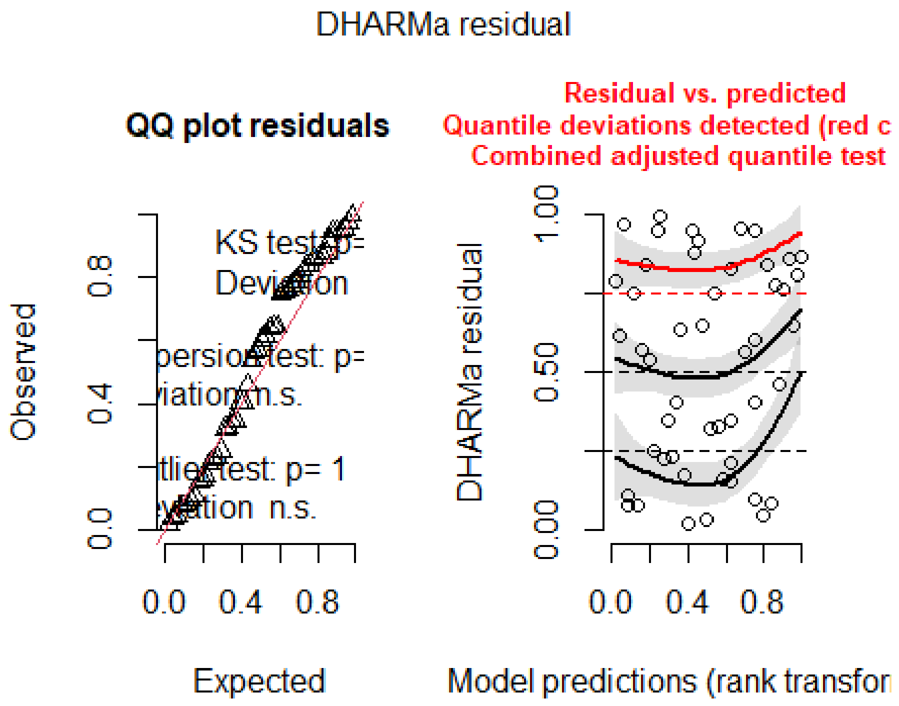

```
par(mfrow=c(2,2), mar=c(5,5,5,4))

# Significance of random effects: P0mother
m10b<-glmmTMB(cop.both~category + shell.size, data=d3, family=binomial)
anova(m10, m10b)

## Data: d3
## Models:
## m10b: cop.both ~ category + shell.size, zi=~0, disp=~1
## m10: cop.both ~ category + shell.size + (1 | P0mother), zi=~0, disp=~1
##      Df    AIC    BIC  logLik deviance Chisq Chi Df Pr(>Chisq)
## m10b  4 53.692 61.340 -22.846   45.692
## m10   5 55.692 65.252 -22.846   45.692     0      1          1

table(d3$cop.both, d3$category)

##
##     Selfer Plastic mixer Plastic switcher Outcrosser
##   0      9             3                2          0
##   1      6             6               24          0
```
