## Supporting Information for "Selfing-outcrossing as a gradient, not a dichotomy: propensity for selfing varies within a population of hermaphroditic animals"

### Supporting Methods: Paternity analyses using COLONY

We repeated all paternity analyses using COLONY version 2.0.5.9 (1), which simultaneously infers sibship and parentage from individual multilocus genotypes. Analyses were run separately for each of the 56 F1 mothers of which juveniles were genotyped. In each analysis, we entered both the mother and all her potential mating partners as candidate fathers, and the maternal family identity as maternal sibship. We also specified two types of locus-specific error rates, one for errors caused by null alleles and one for other types of genotyping errors (for details see 2). Parameters for paternity analyses were set as female and male polygamy, with inbreeding, in a monoecious species, short length of run, full-likelihood analysis at medium precision, without updating allele frequencies. The probability that the correct father was among the candidate fathers was set to 1.

### Figure S1

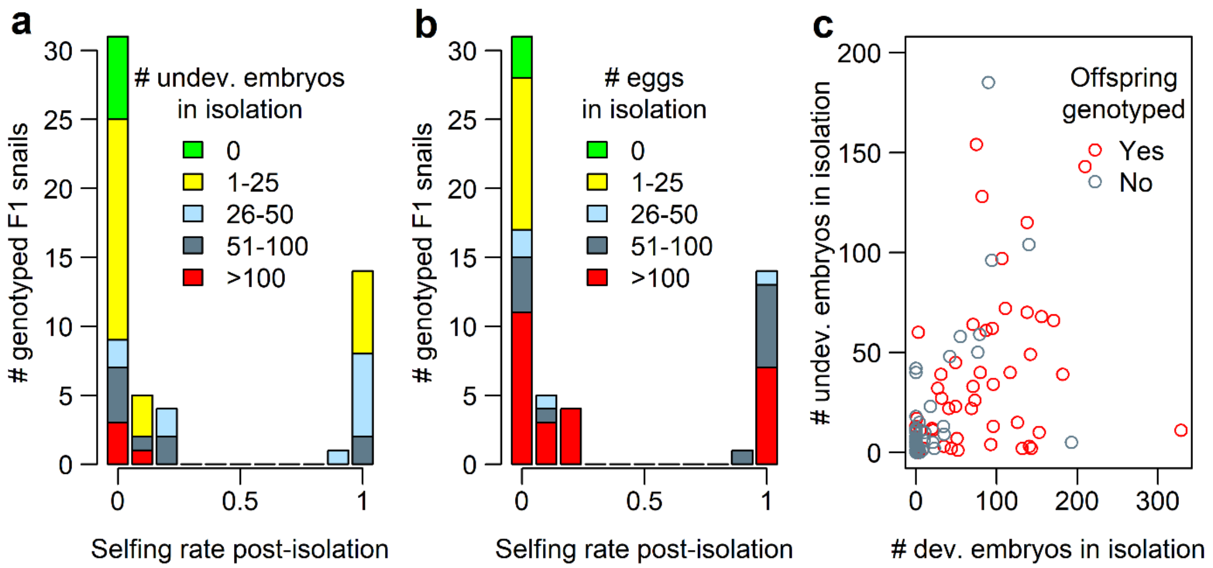


**Fig. S1. Relationship between post-isolation selfing rate and the production of developed and undeveloped embryos in isolation.** Among snails with selfing rate estimates (n = 55), both the production of undeveloped embryos (a) or developed and undeveloped embryos (i.e., eggs) in isolation (b) predicted continued selfing once snails could mate. (c) The similarity between these figures and Fig. 2b is due to a positive correlation between the number of developed and undeveloped embryos produced in isolation ($r = 0.61$, $t_{266} = 12.53$, $p < 0.0001$; n = 268 snails).

### Figure S2

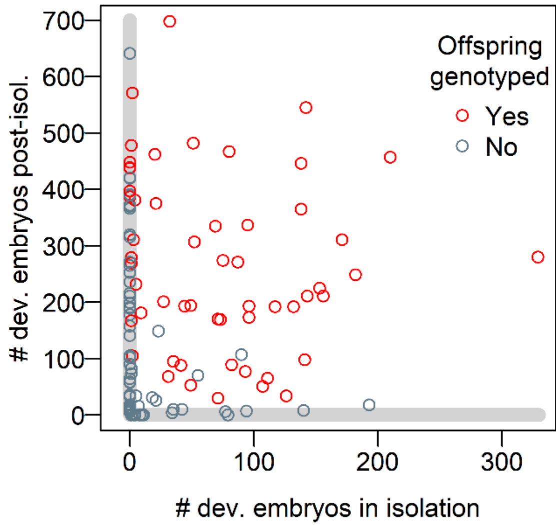


**Fig. S2. Female reproductive success in isolation and post-isolation in female fertile snails (n = 124).** The subset of snails whose offspring were genotyped (n = 55) underrepresents snails that only reproduced before or after the start of mating trials (areas shaded in grey).

### Figure S3

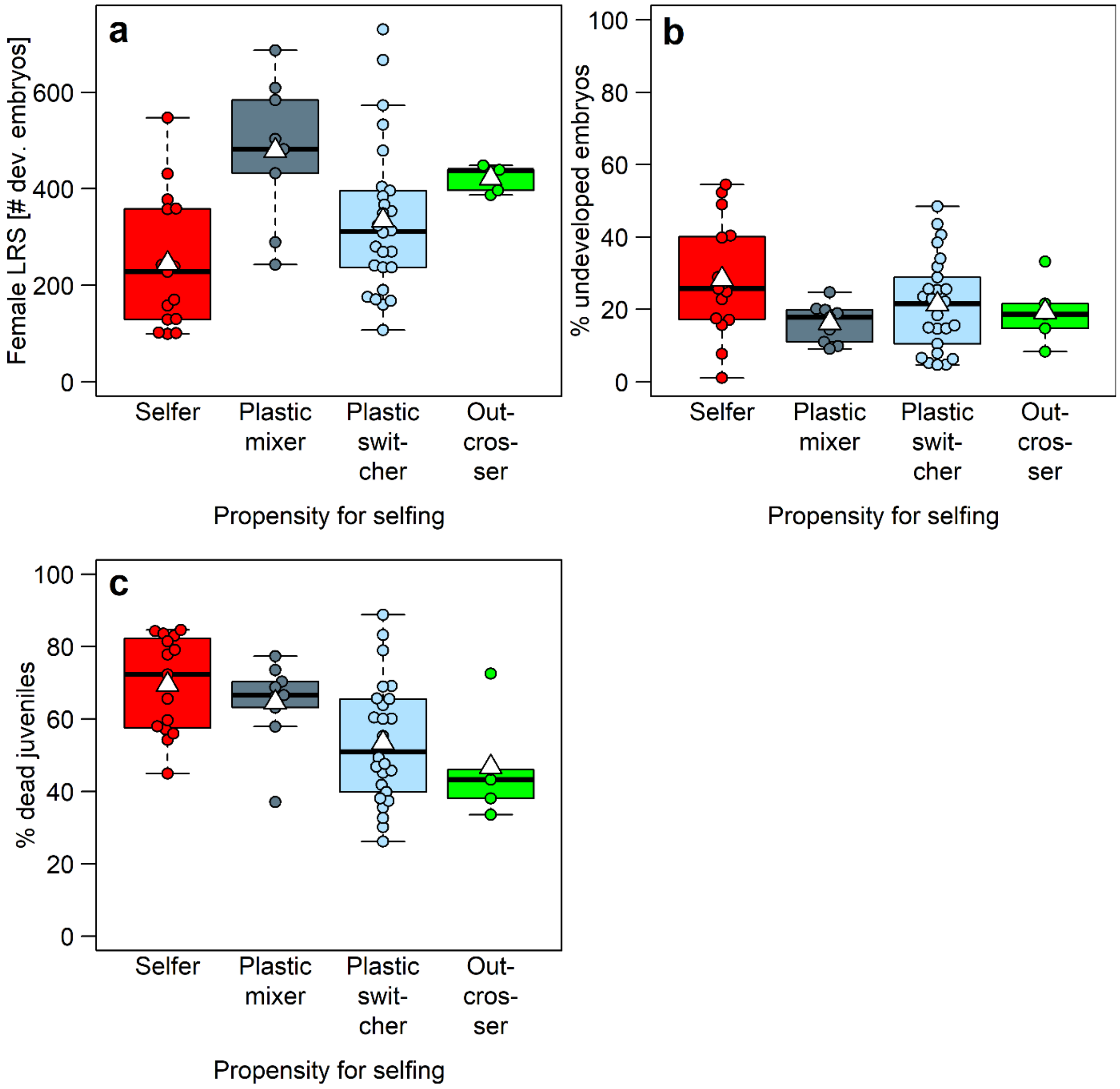


**Fig. S3. Reduced female LRS and increased offspring mortality in preferential selfers when considering only snails with successful selfing rate estimates (n = 55).** Selfers had the fewest developed embryos (a), the highest proportion of embryos that failed to develop (b), and the highest mortality rate among their juvenile offspring. White triangles on boxplots show group means. Outcrossers were excluded from statistical analysis because of low sample size (n = 5).

### Table S1

**Polymorphism data, population-level inbreeding coefficients, and number of loci with mother-offspring mismatches in three generations of snails.** P0 snails were collected in the field on 24 April 2013; F1 and F2 snails are their first- and second-generation laboratory-born descendants, respectively. P0 and F1 snails were genotyped as adults, and F2 snails as juveniles. Forty-one F2 snails with low-quality genotypes were excluded (see Methods for details). The mean number of alleles per locus, the observed and expected heterozygosity, and the population-level inbreeding coefficient $F_{IS}$ over all loci with a 95% CI were estimated using Genetix, version 4.05.2 (3). $F_{IS}$-values in bold are significantly different from zero. We sequenced the mitochondrial cytochrome oxidase subunit I (COI) gene, identified as suitable for species delineation in *Radix* (4), of all 86 P0 snails. All but one of these snails proved to be *R. balthica*, reducing the number of P0 snails to 85. Abbreviations: N, number of snails genotyped; $N_P$, number of polymorphic microsatellite loci; $N_L$, mean number of loci successfully genotyped; $N_A$, mean number of alleles per locus; $H_O$, mean observed heterozygosity; $H_E$, mean expected heterozygosity calculated without bias (5); $F_{IS}$, multilocus inbreeding coefficient according to Weir and Cockerham (6) with a 95% confidence interval based on 10 000 bootstrap iterations (*CI95*); $N_{Lmatoff}$, mean number of loci successfully genotyped in both mother and offspring; $N_{Lmismatch}$, mean number of loci without a maternal allele (i.e., number of loci with a mother-offspring mismatch), $N_{Lnonmat}$, mean number of loci with a single non-maternal allele (i.e., number of loci incompatible with being selfed).

| | N | $N_P$ | $N_L$ ± SD | $N_A$ | $H_O$ | $H_E$ | $F_{IS}$ | *CI95* | $N_{Lmatoff}$ ± SD | $N_{Lmismatch}$ ± SD | $N_{Lnonmat}$ ± SD |
|---|---|---|---|---|---|---|---|---|---|---|---|
| P0 (all) | 85 | 9 | 8.4 ± 1.1 | 11.4 | 0.662 | 0.707 | **0.064** | (0.017, 0.099) | - | - | - |

| | N | $N_P$ | $N_L$ ± SD | $N_A$ | $H_O$ | $H_E$ | $F_{IS}$ | *CI95* | $N_{Lmatoff}$ ± SD | $N_{Lmismatch}$ ± SD | $N_{Lnonmat}$ ± SD |
|---|---|---|---|---|---|---|---|---|---|---|---|
| P0 (grandmothers of F2) | 22 | 9 | 8.7 ± 0.5 | 8.0 | 0.677 | 0.683 | 0.009 | (-0.010, 0.068) | - | - | - |
| F1 (all) | 274 | 9 | 8.9 ± 0.4 | 12.9 | 0.679 | 0.682 | 0.004 | (-0.018, 0.024) | 8.8 ± 0.5 | 0.2 ± 0.5 | 4.6 ± 1.4 |
| F1 (mothers of F2) | 56 | 9 | 8.9 ± 0.4 | 9.7 | 0.688 | 0.684 | -0.006 | (-0.072, 0.049) | 8.8 ± 0.5 | 0.2 ± 0.7 | 4.8 ± 1.5 |
| F2 (genotyped) | 727 | 9 | 8.7 ± 0.7 | 10.8 | 0.572 | 0.665 | **0.140** | (0.120, 0.159) | 8.6 ± 0.8 | 0.1 ± 0.4 | 3.2 ± 2.2 |
| F2 (assigned to a father) | 717 | 9 | 8.7 ± 0.7 | 10.7 | 0.574 | 0.665 | **0.137** | (0.117, 0.155) | 8.6 ± 0.8 | 0.1 ± 0.3 | 3.2 ± 2.2 |

### Table S2

**Number of genotyped F2 offspring per family and quality of paternity assignments.** Genotyping was deemed unsuccessful if fewer than six loci of both mother and offspring were genotyped successfully, OR the offspring genotype lacked a maternal allele at more than two loci, OR six loci of both mother and offspring were genotyped successfully but two of these lacked a maternal allele. Paternity analyses were conducted using a custom-built R routine and COLONY version 2.0.5.9 (1). The probabilities of F2 offspring being selfed (P(selfed)) or sired by a given non-maternal father (P(father)) were estimated using COLONY. Abbreviations: $N_{Fath}$, likely number of (non-maternal) fathers among the genotyped progeny of a F1 snail.

| F1 snail | All genotyped F2 offspring | Successfully genotyped F2 offspring | Selfed F2 offspring | | Outcrossed F2 offspring | | | | Unassigned F2 offspring |
|---|---|---|---|---|---|---|---|---|---|
| | N | N | N | P(selfed) ± SD | N | P(selfed) ± SD | $N_{Fath}$ | P(father) ± SD | N |
| 9_4.2 | 15 | 15 | 0 | - | 15 | 0.00 ± 0.00 | 1 | 0.98 ± 0.01 | 0 |
| 9_5.1 | 5 | 4 | 4 | 0.96 ± 0.02 | 0 | - | - | - | 0 |
| 11_4.1 | 4 | 4 | 0 | - | 4 | 0.00 ± 0.00 | 1 | 0.99 ± 0.00 | 0 |
| 14_3.7 | 15 | 15 | 0 | - | 15 | 0.00 ± 0.00 | 1 | 0.91 ± 0.06 | 0 |
| 17_1.10 | 15 | 15 | 15 | 0.45 ± 0.44[1] | 0 | - | - | - | 0 |
| 22_2.3 | 15 | 15 | 3 | 0.95 ± 0.01 | 12 | 0.00 ± 0.00 | 2 | 0.99 ± 0.01[2] | 0 |
| 22_2.6 | 20 | 18 | 1 | 1.00 | 17 | 0.00 ± 0.00 | 1 | 0.89 ± 0.22 | 0 |

| F1 snail | All genotyped F2 offspring | Successfully genotyped F2 offspring | Selfed F2 offspring | | Outcrossed F2 offspring | | | | Unassigned F2 offspring |
|---|---|---|---|---|---|---|---|---|---|
| | N | N | N | P(selfed) ± SD | N | P(selfed) ± SD | $N_{Fath}$ | P(father) ± SD | N |
| 22_4.1 | 16 | 15 | 0 | - | 15 | 0.00 ± 0.00 | 2 | 0.91 ± 0.11 | 0 |
| 22_5.2 | 15 | 15 | 1 | 0.97 | 14 | 0.00 ± 0.00 | 2 | 0.97 ± 0.04 | 0 |
| 31_2.1 | 15 | 15 | 0 | - | 15 | 0.00 ± 0.00 | 1 | 0.90 ± 0.13 | 0 |
| 31_2.2 | 16 | 16 | 16 | 0.83 ± 0.11 | 0 | - | - | - | 0 |
| 31_3.3 | 15 | 15 | 0 | - | 15 | 0.00 ± 0.00 | 1 | 0.99 ± 0.02 | 0 |
| 31_3.4 | 15 | 15 | 0 | - | 15 | 0.00 ± 0.00 | 2 | 1.00 ± 0.00[2] | 0 |
| 31_3.7 | 8 | 7 | 7 | 0.82 ± 0.05 | 0 | - | - | - | 0 |
| 31_4.2 | 15 | 15 | 0 | - | 14 | 0.00 ± 0.00 | 2 | 0.85 ± 0.25 | 1 |
| 31_4.8 | 10 | 9 | 0 | - | 9 | 0.00 ± 0.00 | 1 | 0.83 ± 0.12 | 0 |
| 34_2.6 | 22 | 13 | 1 | 0.86 | 11 | 0.00 ± 0.00 | 1 | 0.88 ± 0.26 | 1 |
| 34_2.8 | 17 | 13 | 0 | - | 13 | 0.00 ± 0.00 | 1 | 0.84 ± 0.16 | 0 |
| 38_1.1 | 16 | 16 | 0 | - | 16 | 0.00 ± 0.00 | 1 | 0.81 ± 0.25 | 0 |
| 38_2.1 | 14 | 14 | 14 | 0.79 ± 0.11 | 0 | - | - | - | 0 |
| 38_6.1 | 15 | 15 | 0 | - | 14 | 0.00 ± 0.00 | 1 | 0.99 ± 0.01[2] | 1 |
| 40_1.25 | 22 | 18 | 0 | - | 17 | 0.00 ± 0.00 | 1 | 0.96 ± 0.10 | 1 |
| 40_1.6 | 15 | 15 | 3 | 0.96 ± 0.04 | 12 | 0.00 ± 0.00 | 1 | 0.76 ± 0.18 | 0 |
| 40_1.7 | 17 | 15 | 0 | - | 15 | 0.00 ± 0.00 | 1 | 0.93 ± 0.12 | 0 |
| 40_1.8 | 15 | 15 | 0 | - | 15 | 0.00 ± 0.00 | 1 | 1.00 ± 0.00 | 0 |

| F1 snail | All genotyped F2 offspring | Successfully genotyped F2 offspring | Selfed F2 offspring | | Outcrossed F2 offspring | | | | Unassigned F2 offspring |
|---|---|---|---|---|---|---|---|---|---|
| | N | N | N | P(selfed) ± SD | N | P(selfed) ± SD | $N_{Fath}$ | P(father) ± SD | N |
| 40_1.9 | 17 | 16 | 1 | 1.00 | 15 | 0.00 ± 0.00 | 1 | 0.93 ± 0.11 | 0 |
| 40_3.10 | 15 | 15 | 0 | - | 15 | 0.00 ± 0.00 | 1 | 0.99 ± 0.01 | 0 |
| 40_3.16 | 15 | 15 | 0 | - | 15 | 0.00 ± 0.00 | 2 | 1.00 ± 0.00 | 0 |
| 40_3.7 | 13 | 13 | 13 | 0.76 ± 0.14 | 0 | - | - | - | 0 |
| 40_4.2 | 15 | 15 | 0 | - | 15 | 0.00 ± 0.00 | 1 | 0.96 ± 0.04 | 0 |
| 43_5.4 | 11 | 11 | 0 | - | 11 | 0.00 ± 0.00 | 1 | 0.93 ± 0.05 | 0 |
| 43_5.5 | 15 | 15 | 0 | - | 15 | 0.00 ± 0.00 | 1 | 0.93 ± 0.09 | 0 |
| 45_1.2 | 16 | 16 | 16 | 0.84 ± 0.05 | 0 | - | - | - | 0 |
| 45_1.3 | 4 | 4 | 4 | 0.68 ± 0.11 | 0 | - | - | - | 0 |
| 45_2.3 | 15 | 15 | 0 | - | 15 | 0.00 ± 0.00 | 2 | 1.00 ± 0.01 | 0 |
| 45_9.1 | 15 | 15 | 15 | 0.84 ± 0.05 | 0 | - | - | - | 0 |
| 46_2.4 | 7 | 6 | 5 | 0.98 ± 0.02 | 1 | 0.00 | 1 | 0.63 | 0 |
| 51_3.2 | 15 | 15 | 0 | - | 15 | 0.00 ± 0.00 | 1 | 0.91 ± 0.14 | 0 |
| 52_4b.4 | 15 | 15 | 0 | - | 15 | 0.00 ± 0.00 | 1 | 0.97 ± 0.02 | 0 |
| 52_8.4 | 3 | 3 | 0 | - | 0 | - | - | - | 3 |
| 54_3a.1 | 15 | 15 | 2 | 0.99 ± 0.00 | 13 | 0.00 ± 0.00 | 2 | 0.99 ± 0.02 | 0 |
| 61_3.1 | 7 | 4 | 4 | 0.48 ± 0.35[3] | 0 | - | - | - | 0 |
| 61_7.2 | 6 | 6 | 6 | 0.97 ± 0.01 | 0 | - | - | - | 0 |

| F1 snail | All genotyped F2 offspring | Successfully genotyped F2 offspring | Selfed F2 offspring | | Outcrossed F2 offspring | | | | Unassigned F2 offspring |
|---|---|---|---|---|---|---|---|---|---|
| | N | N | N | P(selfed) ± SD | N | P(selfed) ± SD | $N_{Fath}$ | P(father) ± SD | N |
| 63_1.7 | 19 | 17 | 0 | - | 16 | 0.00 ± 0.00 | 1 | 0.95 ± 0.09 | 1 |
| 67_2.7 | 7 | 7 | 0 | - | 7 | 0.00 ± 0.00 | 1 | 0.50 ± 0.31[4] | 0 |
| 70_1.5 | 15 | 15 | 2 | 1.00 ± 0.00 | 13 | 0.00 ± 0.00 | 1 | 1.00 ± 0.01 | 0 |
| 70_1.6 | 12 | 11 | 0 | - | 10 | 0.00 ± 0.00 | 1 | 0.99 ± 0.03[2] | 1 |
| 70_3.14 | 15 | 15 | 0 | - | 15 | 0.00 ± 0.00 | 1 | 0.96 ± 0.04 | 0 |
| 70_3.18 | 13 | 13 | 13 | 0.72 ± 0.20 | 0 | - | - | - | 0 |
| 70_3.20 | 15 | 15 | 0 | - | 15 | 0.00 ± 0.00 | 1 | 1.00 ± 0.00 | 0 |
| 70_3.26 | 15 | 12 | 1 | 0.92 | 11 | 0.00 ± 0.00 | 3 | 0.97 ± 0.06[5] | 0 |
| 70_9.5 | 19 | 16 | 0 | - | 16 | 0.00 ± 0.00 | 1 | 0.97 ± 0.04[2] | 0 |
| 75_7.1 | 17 | 16 | 0 | - | 15 | 0.00 ± 0.00 | 1 | 0.98 ± 0.03 | 1 |
| 79_2.1 | 15 | 15 | 0 | - | 15 | 0.00 ± 0.00 | 1 | 0.72 ± 0.27 | 0 |
| 79_4.4 | 5 | 4 | 4 | 0.99 ± 0.01 | 0 | - | - | - | 0 |
| 83_3.4 | 15 | 15 | 15 | 0.78 ± 0.21 | 0 | - | - | - | 0 |

[1]Seven offspring had a single, identical allele absent in both the mother and her single mating partner. Four of these offspring were left unassigned by COLONY. Apart from this allele, offspring genotypes were fully compatible with being selfed, suggesting that the non-maternal allele resulted

from a mutation in the maternal germ line. Meanwhile, offspring each lacked alleles from the non-maternal father at 3-6 loci, making outcrossing exceedingly unlikely. Among offspring without the mother-offspring mismatch, P(selfed) was 0.83 ± 0.10.

[2]In these five families, one offspring each was left unassigned by our R-routine but was assigned by COLONY with P(father) ≥ 0.91. Because of the high P(father) and because in all five families other offspring were assigned, by both COLONY and our R-routine, to the respective father as well, we accepted COLONY's assignment.

[3]All four offspring had one locus lacking a maternal allele, thereby decreasing P(selfed). However, all offspring had three loci lacking an allele from the single potential non-maternal father, rendering outcrossing unlikely.

[4]The lower P(father) likely resulted from the relative similarity of the two parents' genotypes, with only four alleles private to the father. However, all offspring possessed 2-4 of these private paternal alleles, rendering selfing unlikely.

[5]In this family, three offspring were assigned to different, yet always non-maternal, fathers by our R-routine and COLONY. The COLONY assignments involved a putative father who lacked an allele from the three offspring at 2-4 loci each, while the R-routine assigned the offspring to fathers with at most a single father-offspring mismatch. We thus gave preference to the assignment of our R-routine.

### Table S3

**GLMMs on female LRS, its components, and juvenile mortality when considering only snails with estimated selfing rate.** Results are shown for four models using only snails with offspring that were genotyped and successfully assigned to a father (n = 50; five outcrossers were excluded due to low sample size). Response variables were the lifetime number of developed embryos (female LRS) and eggs, the lifetime proportion of undeveloped embryos, and the proportion of developed embryos that died before they could be genotyped (*i.e.*, before reaching the juvenile age of 12.3 ± 0.5 weeks). For response variables 1 and 2, negative binomial errors were fitted, and results are provided on the log scale. For the two response variables that are proportions, Gaussian errors were fitted, as they did not deviate significantly from normality and as model assumptions were fulfilled. Fixed effects were propensity for selfing (reference level: selfers) and body size (shell length). As random intercepts we included P0 mother identity (non-significant: $\chi^2_1 \leq 0.94$, $p \geq 0.33$) and, in all models except that of the number of eggs, pair identity on mating opportunity 1 (non-significant: $\chi^2_1 \leq 1.17$, $p \geq 0.28$). S.E.: standard error, PS.: propensity for selfing.

| Response | Predictor | Estimate (S.E.) | *z*-value | *p*-value |
|---|---|---|---|---|
| # dev. embryos (LRS) | Intercept | 4.14 (0.57) | 7.27 | < 0.0001 |
| | PS (plastic mixer) | 0.67 (0.16) | 4.11 | <0.0001 |
| | PS (plastic switcher) | 0.36 (0.14) | 2.53 | 0.0115 |
| | Body size | 0.09 (0.04) | 2.46 | 0.0138 |
| # eggs | Intercept | 4.30 (0.51) | 8.45 | < 0.0001 |
| | PS (plastic mixer) | 0.51 (0.15) | 3.46 | 0.0005 |
| | PS (plastic switcher) | 0.28 (0.13) | 2.19 | 0.0285 |
| | Body size | 0.10 (0.03) | 3.10 | 0.0019 |
| % undev. embryos | Intercept | 17.55 (16.89) | 1.04 | 0.30 |
| | PS (plastic mixer) | -12.59 (5.37) | -2.34 | 0.0192 |

| Response | Predictor | Estimate (S.E.) | *z*-value | *p*-value |
|---|---|---|---|---|
| | PS (plastic switcher) | -5.35 (4.25) | -1.26 | 0.21 |
| | Body size | 0.72 (1.14) | 0.63 | 0.53 |
| % dead juveniles | Intercept | 116.78 (18.04) | 6.47 | < 0.0001 |
| | PS (plastic mixer) | -3.40 (5.53) | -0.62 | 0.54 |
| | PS (plastic switcher) | -17.76 (4.49) | -3.95 | < 0.0001 |
| | Body size | -3.30 (1.22) | -2.69 | 0.0070 |
